# Transposon- and genome dynamics in the fungal genus *Neurospora*: insights from nearly gapless genome assemblies

**DOI:** 10.1101/2020.09.27.311811

**Authors:** Diem Nguyen, Valentina Peona, Per Unneberg, Alexander Suh, Patric Jern, Hanna Johannesson

**Author notes:** Corresponding author: Hanna Johannesson, Department of Organismal Biology, Evolutionary Biology Centre, Uppsala University, SE-75236 Uppsala, Sweden. Email addresses: DN VP PU AS PJ HJ.

## Abstract

**Background:** A large portion of nuclear DNA is composed of transposable element (TE) sequences, whose transposition is controlled by diverse host defense strategies in order to maintain genomic integrity. One such strategy is the fungal-specific Repeat-Induced Point (RIP) mutation that hyper-mutates repetitive DNA sequences. While RIP is found across Fungi, it has been shown to vary in efficiency. To date, detailed information on the TE landscapes and associated RIP patterns exist only in a few species belonging to highly divergent lineages.

**Result:** We investigated 18 nearly gapless genome assemblies of ten *Neurospora* species, which diverged from a common ancestor about 7 MYA, to determine genome-wide TE distribution and their associated RIP patterns. We showed that the TE contents between 8.7-18.9% covary with genome sizes that range between 37.8-43.9 Mb. Degraded copies of Long Terminal Repeat (LTR) retrotransposons were abundant among the identified TEs, and these are distributed across the genome at varying frequencies. In all investigated genomes, TE sequences had signs of numerous C-to-T substitutions, suggesting that RIP occurred in all species. RIP signatures in all genomes correlated with TE-dense regions.

**Conclusions:** Essentially gapless genome assemblies allowed us to identify TEs in *Neurospora* genomes, and reveal that TEs contribute to genome size variation in this group. Our study suggests that TEs and RIP are highly correlated in *Neurospora*, and hence, the pattern of interaction is conserved over the investigated evolutionary timescale. We show that RIP signatures can be used to facilitate the identification of TE-rich region in the genome.

## Background

Advanced sequencing technologies have yielded numerous genome assemblies of high quality from diverse species across the tree of life (from NCBI Assembly database, [1]). Despite the availability of these assemblies, there are still difficult-to-sequence genomic regions, such as repetitive sequences and centromeres [2, 3]. The inability to sequence these regions has limited studies of genome architecture and evolution, which depend on data from multiple genomes. Thus, availability of high-quality gapless genome assemblies from multiple individuals and species are critically needed, particularly to understand how repetitive sequences contribute to genome size evolution and how genome integrity is maintained.

Genomes are dynamic entities, subject to large and small-scale rearrangement events that can lead to gains and losses of genomic DNA sequences. Transposable elements (TEs) are genetic elements present in eukaryotic and prokaryotic genomes, characterized by their ability to propagate in the genome. While TEs may contribute to evolutionary change and innovation [4], TE activity can lead to deleterious effects including insertional disruption of functionally important sequences such as promoters or genes themselves, gene miss-expression, and silencing of adjacent genes [5, 6]. TEs can also mediate ectopic recombination between distant chromosomal regions, inversions, and deletion of genomic sequences [7, 8].

Based on the nature of their transposition intermediates, TEs can be classified as class I RNA retrotransposons (RNA intermediate) or class II DNA transposons (DNA intermediate) [9, 10]. They are further divided into subclasses, orders, and super-families based on their specific structural and coding features. Class I elements, consisting of LTR retroelements, LINEs (Long Interspersed Nuclear Elements) and SINEs (Short Interspersed Nuclear Elements), are the most frequent class of TEs in many animals (e.g., *Drosophila melanogaster*), plants (e.g., *Arabidopsis thaliana* and maize), and fungi [11-14].

To counterbalance potential detrimental side effects of genome-damaging agents, genome defense mechanisms have evolved to ensure the maintenance of genome integrity [15-17, 18. Eukaryotes have evolved ways to defend their genomes against TEs {Gladyshev, 2016 #2852]. For example, plants and animals utilize methylation and RNA interference (RNAi)-based mechanisms [19, 20]. In fungi, one such mechanism is sequence homology-based Repeat-Induced Point (RIP) mutation [21, 22]. First discovered in *Neurospora crassa*, RIP permanently mutates duplicated sequences, such as TEs, and introduces multiple C-to-T transition mutations, predominantly at CpA dinucleotides, in both copies of the sequences, skewing dinucleotide frequencies towards an over-representation of TpA in mutated sequences [21, 22]. For RIP mutations to occur, there are requirements for minimal duplicated sequence length of about 400 base pairs (bp), though sequences as short as 155 bp have been reported, and sequence identity of greater than 80% [23-25]. In evolutionarily diverged fungal species where RIP has been experimentally demonstrated, RIP is “leaky”, and intact and active TEs are present in the genomes, reviewed in [26]. In contrast, RIP is considered highly efficient in *N. crassa* and has resulted in the essential absence of intact and active TEs [27].

Here we study the distribution of TEs and RIP in the filamentous fungal genus *Neurospora*. The motivation of our study is fourfold. First, the genus *Neurospora* represents a tractable system; it has small genomes relative to other eukaryotes (∼43 Mb) [28, 29], and the model organism *Neurospora crassa* is well-studied and has a well-annotated genome [30]. Second, despite its relatively small genome, about 10% of the *Neurospora* genome is comprised of repetitive DNA sequences [29]. Third, the efficiency of RIP varies across the kingdom Fungi. In *N. crassa* RIP is thought to be efficient at controlling transposons, while in distantly related fungi, RIP is “leaky” resulting in the persistence and presence of active TEs [27]; comparisons over short evolutionary time is needed to understand the evolution of this key genomic feature. Fourth, in addition to the well-studied *N. crassa*, the phylogenetic relationship between species whose last common ancestor of the clade of *Neurospora* diverged within the last 7 million years, is well resolved [31-34], providing a phylogenetic framework for studies of genome evolution in this group of fungi [35]. Altogether, *Neurospora* genomes present a well-suited model system for studying the relationship between TEs and genome defense.

Our study of the TE landscape in *Neurospora* was facilitated by high-quality nearly gapless genome assembly of long-read sequenced genomes. We thus comprehensively surveyed TEs based on their diversity, sequence abundance, and chromosomal distribution in each genome assembly, and compared these features between and within species to better describe the dynamics of TE variability. Furthermore, we determined how RIP associated with TEs among the different genomes in this *Neurospora* clade to better understand whether RIP efficiency in *N. crassa* is a general feature of *Neurospora*. With our study, we provide novel insights into the application of genome-wide RIP index to identify TE enriched locations in the genome, when identifying TEs in genomes heavily affected by RIP is not possible.

## Results

### *Nearly gapless* Neurospora *assemblies*

To study the TE and RIP landscapes of closely-related *Neurospora* species (**Figure 1**) [31, 36, 37], we generated PacBio assemblies from four genomes, of which two previously lacked genomic information [*N. hispaniola* (FGSC 8817), and *N. metzenbergii* (FGSC 10397)] (**Table S1**). Genomic data from *N. intermedia* (FGSC 8767) and *N. discreta* (FGSC 8579), generated using short-read sequencing technologies, have previously been published [38, 39]. Additionally, we used 11 previously published PacBio-sequenced genome assemblies [i.e. from the genomes of *N. intermedia* (FGSC 8761, 8767), *N. sitophila* (FGSC 5940, 5941, W1426, and W1434), and *N. tetrasperma* (FGSC 2503, 2504, 9045, 9046, and 10752)] [39-43]. Reference assemblies are available for *N. crassa* (FGSC 2489) and *N. tetrasperma* L6 (FGSC 2508, 2509) [29, 44]. In total, we analyzed 18 genomes from 10 *Neurospora* species, of which 15 genomes were PacBio-sequenced, resulting in dense taxon sampling both across the phylogeny and within specific species (*N. sitophila* (4 strains) and *N. intermedia* (3 strains)) (**Figure 1, Table S1**).

**Figure 1.**
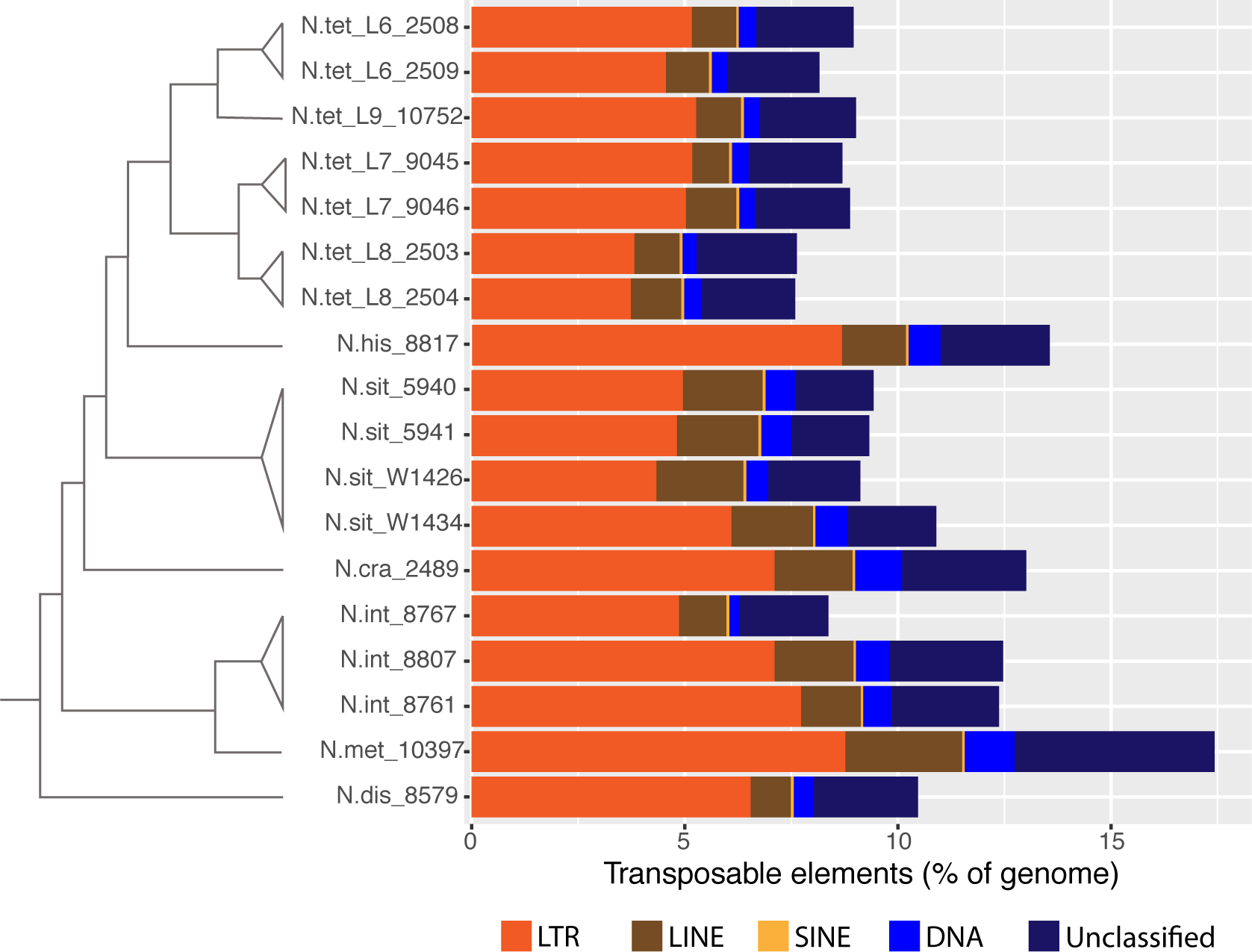
Schematic *Neurospora* phylogeny and TE-distributions. The left panel phylogeny was modified from [31, 36, 99]. The right panel indicate TE composition in each genome determined by calculating the number of TE nucleotides relative to the assembly length (in percent) for each of the TE families (Class I: LTR, SINE, and LINE; Class II: DNA transposons; Unclassified TEs).

The assemblies of the PacBio-sequenced genomes were nearly gapless. We recovered contigs that could be anchored to the seven chromosomes in each of the PacBio genome assemblies that were collinear with the seven chromosomes in *N. crassa* [45], indicating that our assemblies were at the chromosome level. One assembly (strain FGSC 5940) had no gaps [the one excess contig is mitochondrial DNA], and from three other strains (W1434, and FGSC 8807 and 9046) we identified up to 8 unplaced contigs that represent gaps in the assembly resulting from the repetitive ribosomal RNA genes. Two strain of *N. intermedia* (FGSC 8767 and 8761) had particularly more “gappy” assemblies, where 24 and 10 contigs mapped to the 7 *N. crassa* chromosomes, respectively. For all PacBio assemblies, the 1 to 29 unplaced contigs that could not be mapped to the *N. crassa* reference genome were mitochondrial DNA and ribosomal RNA genes. The genome assemblies for examined *Neurospora* genus ranged between 37.8 and 43.9 Mb, and *N. crassa* was 41 Mb (**Table S1**).

### The curated Neurospora TE library

TE identification in the nearly gapless genome assemblies involved sequence similarity searches based on a modified version of an existing query repeat sequence library [46]. The original 978 repeats library was curated and reduced to 844 repeats. Specifically, we removed cellular genes and sequences lacking transposon features such as transposase, reverse transcriptase, RNaseH, integrase, *gag*, protease. Consensus sequences for elements previously identified by RepeatModeler that had no identified domains in the NCBI Conserved Domain Database were retained in the library. However, full-length LTR retrotransposon candidates determined by LTRharvest that had no identifiable domains or had only CHROMO domains, or were cellular genes, were removed. An additional 41 *N. crassa* TE sequences obtained from NCBI GenBank, from Selker et al. [47], and from Wang et al. [48] were added to the updated TE query library for a total of 885 repeats for RepeatMasker analyses, and the library has been made available at: https://doi.org/10.6084/m9.figshare.c.4310996 [49].

### Genomic TE content rich in LTR retrotransposons

The *Neurospora* genomes were comprised of both Class I RNA retrotransposon and Class II DNA transposon sequences. These TE sequences represented 7.66 Mb of genomic DNA (17.43% of genome) in *N. metzenbergii*, the species with the largest genome. In two of the species with the smallest genomes, these TEs represented less than 4 Mb of the genomes (*N. sitophila*: 3.45 Mb, 9.11% of genome; and *N. discreta*: 3.95 Mb, 10.46% of genome) (**Figure 1**; **Table S2**). We found that TE comprised 13.02% (5.34 Mb) of the *N. crassa* genome, which is similar to a previous report [29].

The proportion of genomes occupied by the different TE families (e.g., LTR retrotransposon, LINE, or DNA elements) varied considerably among the *Neurospora* species (**Figure 1, Figure S1**). However, the LTR retrotransposon sequences were most abundant in all *Neurospora* genomes (**Figure 1**). They made up the largest fraction of the TE sequences (1.44 - 3.85 Mb or 3.74 - 8.77% of genomes), followed by LINE elements (0.34 - 1.20 Mb or 0.86 - 2.74% of genomes), DNA elements (0.09 - 0.51 Mb or 0.24 - 1.17% of genomes), and Unclassified repeats (0.70 - 2.06 Mb or 1.82 - 4.69% of genomes) (**Table S2, Figure S1**).

### Genome size correlation with TE content

In *Neurospora* species, TE content correlated positively with genome size (**Figure 2**), suggesting TE contribution to genome size expansion in *Neurospora*. A number of genomes fell outside the 95% confidence interval of the regression; these genomes either had more or fewer TEs than expected. For example, *N. discreta* and *N. sitophila* (strains FGSC 5940 and W1434, W1426) had more TE sequences than expected relative to their genome sizes, as predicted using the correlation between the remaining species. Alternatively, both strains of each of the *N. tetrasperma* L6, L7 and L8 had fewer TEs than expected relative to their genome sizes (**Figure 2**). When zooming in on the different TE families, we found that LTR retrotransposon and the Unclassified repeats positively correlated with larger genome sizes, while other TEs were weakly or not at all correlated to genome size (**Figure S2**).

**Figure 2.**
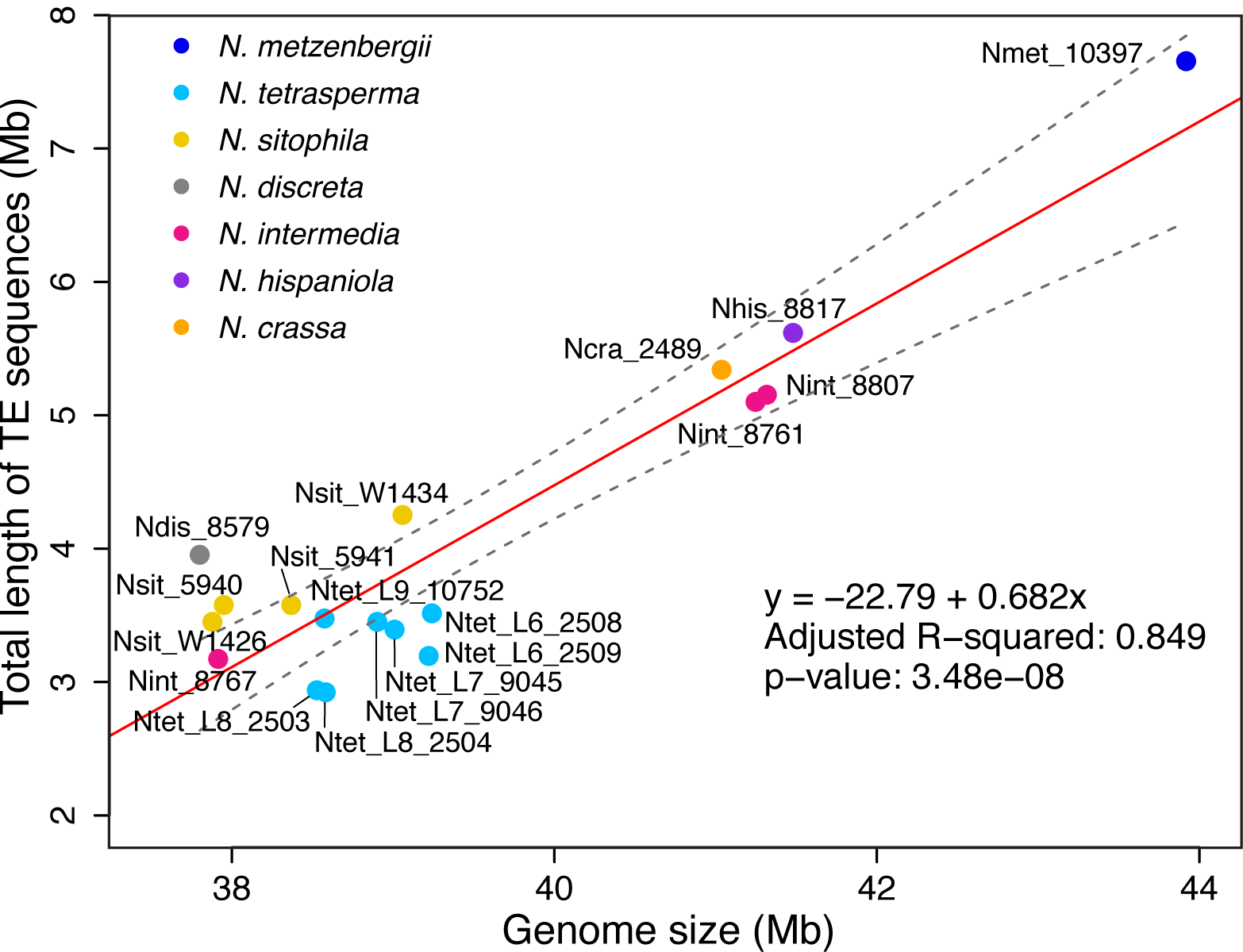
*Neurospora* genome size correlation with TE content. Total genomic TE lengths (in base pairs) correlated positively with total *Neurospora* genome assembly lengths. Each *Neuropspora* species was color coded, and multiple strains from different species of *N. tetrasperma* were grouped together.

### TEs locate across all seven Neurospora chromosomes

Numerous TE sequences were identified on each of the seven *Neurospora* chromosomes, but sliding window analyses suggest certain regions per chromosome were particularly enriched (log2 observed/expected values >2) (**Figure 3, Figure S3**, top panels). Similar patterns of enrichment were observed between orthologous chromosomes in each of the *Neurospora* species (**Figure S3**, top panels).

**Figure 3.**
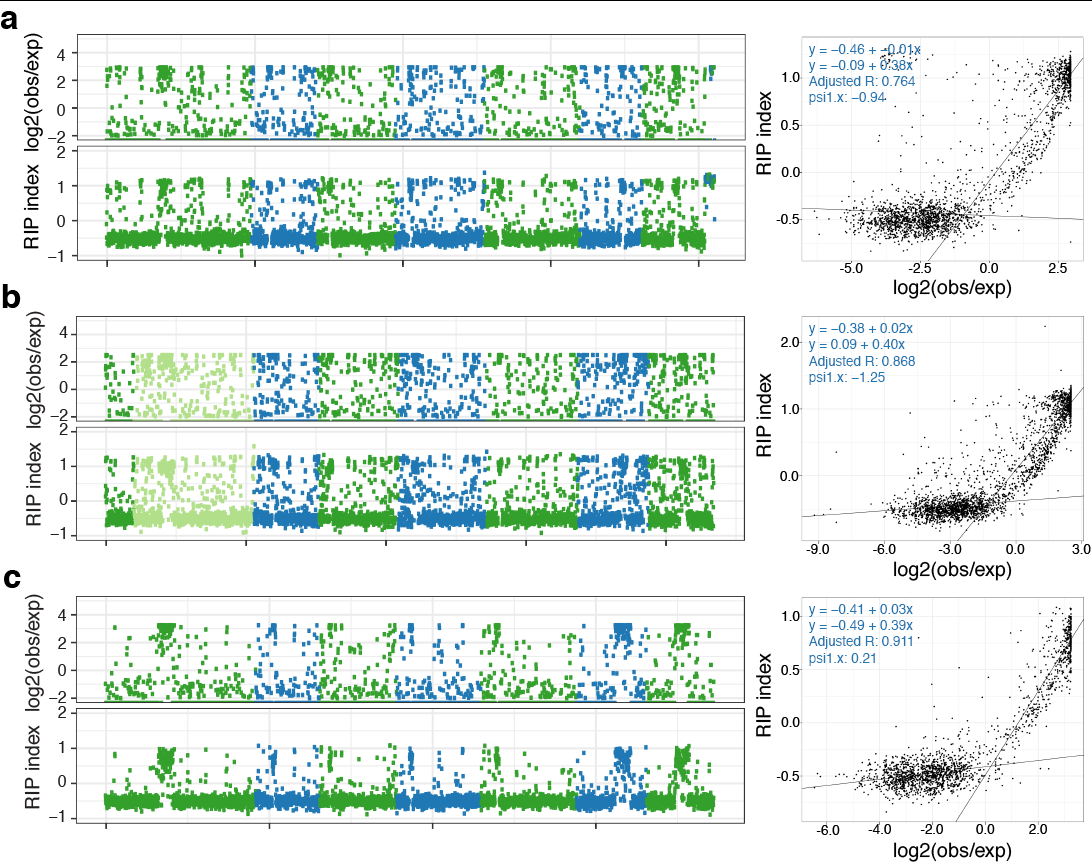
Genome-wide TE landscape and RIP signature along *Neurospora* chromosomes. a) *Neurospora crassa* (model organism), b) *N. metzenbergii* (strain with largest genome), c) *N. discreta* (strain with one of the smallest genome). Top panel for each species indicates the enrichment of transposable element (TE) sequences (log2(observed repeat (bp)/expected repeats (bp)) determined in 10 kb windows. Each dot represents a 10-kb window. Values above 2 are herein described as enriched in TE sequences. Bottom panel for each species indicates the genome-wide composite RIP index, independent of underlying genomic content, determined in 10 kb windows. Each dot represents a 10-kb window. Positive values were herein described as a window contained sequences that experienced RIP mutation. For all plots, the alternating colors between blue and green indicate the alternation between chromosomes, with chromosome numbers following the alignment to *N. crassa*. The lighter shading indicates the presence of multiple contigs for the respective chromosome. Right panel for each species indicates the correlation between a 10-kb window for TEs and the corresponding 10-kb window for composite RIP index. Regression lines were fit using a segmented linear regression model in which the breakpoint (“psi”) is also estimated. The top regression line is the estimated fit for log2 scores < psi value (left line) and the bottom regression line is the estimated fit for log2 scores > psi value (right line). The adjusted R^2^ (“adjusted R”) value calculates fit of the regression lines to the data.

### RIP signatures were detected in all Neurospora species

The composite RIP index can be used as a proxy for the RIP activity in a genomic region; positive values indicate that RIP has occurred [50]. For this study, we developed a method to determine genome-wide RIP indices independent of the underlying genomic context. We use the *N. crassa* genome assembly as an example to explain the method that was also applied to all genome assemblies. First, to validate that the composite RIP index could be applied across a genome to indicate regions of RIP activity, we used the *N. crassa* genome as reference and generated mock genomes with equal length to the reference. The mock genomes were generated either by scrambling the sequenced genome or selecting nucleotides based on the observed nucleotide frequencies. Observed TE intervals were randomly placed on the mock genome to calculate composite RIP index. As expected, no RIP signature was observed in the mock genomes (**Figure S4**, “shuffle” and “frequency”), whereas the reference *N. crassa* genome (**Figure S4**, “obs”) indicated RIP activity in some sections but not in others that likely represent gene-rich regions. Similar patterns were observed for the other 17 genomes in this study (**Figure S4**).

Composite RIP index values were also calculated for 10 kb windows, delineated independently of the sequence context or genomic content (e.g., genic regions, repetitive regions) for each genome assembly, and plotted over the length of the chromosome (**Figure 3, Figure S3**, bottom panels). Positive values indicated that a region had experienced RIP activity. Many TEs have RIP signature, based on visual inspection of sequence alignments. Note that these RIPed regions coincided with windows enriched in TEs (**Figure 3, right panels, Figure S5**). Taken together, on a genome-wide scale, TE sequences were observed to have experienced RIP.

### TE partly accounts for intra-species genome size variation

We observed genome size variation between strains of *N. sitophila* and *N. intermedia* (**Table S1**). To better understand the variation within species, particularly whether TEs accounted for the gain or loss of DNA, we compared the high-quality nearly gapless genome assemblies of four independent *N. sitophila* and three independent *N. intermedia* strains (**Table S1**).

We found genome size differences among *N. intermedia* as the FGSC 8767 assembly was approximately 3.4 Mb smaller than FGSC 8807 and FGSC 8761 (**Table S1**), and the difference was not due to poor data quality, since we confirmed the data quality with Illumina sequenced data. TEs contributed to 1.95 Mb of this size difference, leaving 1.4 Mb unaccounted for the remaining differences (**Table S2**). The FGSC 8767 assembly was, however, more fragmented and 24 contigs mapped to *N. crassa* where each chromosome had between 2 and 7 mapped contigs (e.g., chromosome 1 was assembled from 7 contigs), and an excess of 11 contigs representing gaps in the assembly, due to mitochondrial DNA and ribosomal RNA genes. In contrast, the assemblies for FGSC 8761 and 8807 had 10 and 7 contigs mapped to *N. crassa*, respectively, and correspondingly 17 and 7 gaps, respectively. Therefore, the difference in genome size may be due to unassembled repetitive sequences in the FGSC 8767 genome, demonstrating the importance of high-quality and nearly gapless assemblies for more specific/detailed whole-genome comparisons within and across species.

Among the four *N. sitophila* strains, W1434 had approximately 1 Mb larger genome than the other three strains (W1426, FGSC 5940, FGSC 5941: 37.9-38.4 Mb) (**Table S1**). The W1434 assembly had 7 contigs mapped to *N. crassa* where each chromosome was fully mapped, though there were 7 unmapped contigs that represent mitochondrial DNA and ribosomal RNA genes. The better assembled FGSC 5940 had 7 contigs mapping to *N. crassa* where each chromosome was fully mapped, and 1 mitochondrial DNA contig. In contrast, the W1426 and FGSC 5941 assemblies had 12 and 18 unplaced contigs, respectively, with 8 and 7 contigs mapping to the chromosomes in *N. crassa*. The larger genome size may be a consequence of different genome rearrangements, for example by accumulation of certain TE families, rather than assembly quality due to fragmentation. Indeed, the size difference could be accounted for by an excess in repetitive sequences that were dispersed along the seven chromosomes (**Figure S3**). The TE content of W1434 was about 0.7 Mb larger (4.25 Mb compared with 3.45-3.58 Mb in W1426, FGSC 5940, FGSC 5941), of which .589 Mb (82%) of sequences were contributed by LTR retrotransposons alone (**Table S2**).

### TEs were recently active in Neurospora

The current understanding of TEs in *N. crassa* is that they are not active. Whether this dogma is true for other *Neurospora* species could be tested by searching for lineage-specific TE insertions. We scored pairwise alignments of individual TE-flanking regions for TE presence and absence. These insertions indicate recent TE accumulation, and refer to them as “pairwise lineage-specific” herein. In general, few pairwise lineage-specific TEs could be detected; we found between 0 and 23 insertions per genome (**Table S3**). These were either LTR and LINE retrotransposons from 82 subfamilies (**Table S4**). In total, we found 593 pairwise lineage-specific LTR retrotransposons (of which 406 putative full-length insertions; **Tables S3** and **S4**) belonging to 76 subfamilies and 509 LINE belonging to 6 subfamilies (**Tables S3** and **S4**) suggesting a diversification of LTRs. Most of the insertions (32%) were represented by ncra_Tad1_01 LINE subfamily, followed by ncra_LTR_69, ntet_Tad1_01, Tad1.1 (4% each), ntet_LTR_18, ncra_LTR_49, ncra_Tad1_06 (3% each; **Table S5**).

## Discussion

Genomes integrity is maintained by different mechanisms to ensure genome function and successful transmission of genetic material to the next generation. However, variation in genomic DNA content can still be observed. Extensive intra- and inter-species variation in genomic content can be found across eukaryotic lineages (e.g., in [51, 52]). The study of genome content variations has been made possible by availability of high-quality genome assemblies via the employment of next generation and newer sequencing technologies, increasingly available at low costs and with increasing efficiency and accuracy.

The first report on the 38 Mb genome assembly of *N. crassa* indicated that 10% of genome consisted of repetitive elements, which were detected based on filtering of alignments longer than 200 bp [29]. In our study, we found that 13% of the genome in *N. crassa* consists of TE sequences. The difference between our studies may be due to the underrepresentation of SINE elements and truncated elements in the original draft genome. We now add to the body of knowledge that closely-related *Neurospora* species have similarly large proportions of their genomes comprised of repetitive elements. Nevertheless, we found variation in repetitive content, both between and also within species. Among our investigated *Neurospora* genomes, TE contents vary between 8.7 and 18.9% of the genome. TE content variation among fungal genomes have been reviewed previously [53], but comparisons have typically been studied in species that have diverged over long evolutionary time scales. For example, *Saccharomyces cerevisiae* had 3.1% of the 12 Mb genome consisting of TEs from five families, all LTR retrotransposons [54], which is in contrast to other species with high TE contents, including *Blumeria graminis f. sp. tritici* (90% TEs, 97.4 Mb genome) and *Tuber melanosporum* (60.1% TEs, 123.6 Mb genome) [11, 55]. Here in this study we characterize the variation on a shorter time scale.

Our study involves broad sampling of *Neurospora* species, of which two are represented by multiple strains providing a brief look into population-level variation. These types of data can provide a broader picture of TE dynamics between strains of a species in a population, and over a time scale where TEs can transpose to new sites, rather than a snapshot provided by one (up to four) strains per species. Large genome size variation among individual species in a population has been attributed to TEs (e.g. [56] and [57]). With sampling of up to four strains in *N. sitophila* and *N. intermedia*, we also saw variation in TE content, though these strains were sampled from diverse locales (*N. sitophila*: Italy and French Polynesia; *N. intermedia*: Taiwan, India, Indonesia). For instance, the larger *N. intermedia* FGSC 8767 is a resistant spore killing Sk-2 strain, while the shorter FGSC 8807 and FGSC 8761 are both sensitive strains to spore killing, and it is possible that TE content correlate with this function, warranting further investigation of the effects of TEs on host genome function and evolution.

Our view of TE dynamics over short evolutionary time is supported by our identified specific TE insertions. These numbers are small, and primarily represented by LTR retrotransposons and LINE elements. We used a conservative analysis pipeline based on identification of orthologous regions flanking the TE and queried whether the corresponding region contains a TE or not (thus intact or uninterrupted by a TE) in the contrasting genome. TEs often clusters and/or nest within other TEs over time [58]. Furthermore, RIP mutations have been shown to extend beyond the duplicated target region [59], complicating identification of orthologous TE flanking sequences between the compared genomes. Thus, the low number of observed genomic pairwise TE differences could be hampered by nested insertion and RIP activities indicating use of broad sampling and sequencing to resolve most individual TE loci along the host phylogeny.

Taken together, our results on variation of TE content among strains and species of *Neurospora*, indicate that, albeit the genome being well protected [60], TEs have been actively transposing in *Neurospora* genomes over the relatively short evolutionary time scale. However, we cannot at this point exclude the possibility that the variation in TE content between *Neurospora* strains and species is a result of differential loss of an ancestral pool of TEs in the different lineages.

All sampled *Neurospora* genomes were rich in LTR retrotransposons. The abundance of this group of repetitive sequences has also been observed in yeasts, plants and animals [12-14]. One may speculate that LTR retrotransposon sequences are especially numerous in *Neurospora* genomes, because they propagate via an RNA intermediate and copy-paste mechanism, and that they are longer than other numerous elements utilizing similar replication strategies including LINEs and SINEs, thereby contributing larger fractions of nuclear DNA. LTR retrotransposons can also form numerous short solitary LTRs (solo-LTRs) [61]. Furthermore, the fraction of Unclassified repeats was quite large and likely contains solo-LTRs and (non-autonomous) DNA transposons, highlighting a classification problem that contribute to an annotation and detection of TEs problem.

Genome size has been reported to be positively correlated with the abundance of TEs in diverse lineages of eukaryotes [62, 63]. While these studies have been compared across distant and diverged lineages, few have been conducted with closely-related species [64-66]. In our study with *Neurospora* species that diverged from a common ancestor within the last 7 million years, we observed similar genome size-TE correlations. The correlation found in our study is interesting given that TEs are expected to be inactive in *Neurospora*, and indeed, we have verified a conserved genome defense system in this genus. As mentioned above, we cannot rule out that the difference is due to shared common TEs that have been differentially deleted in the different lineages. As previously demonstrated, TEs contribute to genome size evolution, though their net effect on genome size depends on TE accumulation relative to the overall deletion rate [67].

To maintain genome integrity, different organisms have evolved different methods to defend their genomes against parasitic elements [16]. In *Neurospora crassa*, four genome defense mechanisms control TEs, either post-transcriptionally via RNAi-based mechanisms (e.g. quelling and meiotic silencing of unpaired DNA) or transcriptionally (e.g. RIP) and DNA methylation [15, 68]. RIP has been demonstrated experimentally in other filamentous ascomycetes: *Podospora anserina* [69, 70], *Magnaporthe grisea* [71], *Leptosphaeria maculans* [72] and *Nectria haematococca* [73]. Repetitive regions in *N. crassa* coincided with RIP mutated regions and previously observed to be methylated regions [21, 22].

Our study demonstrates that RIP mutation signatures can be computationally assessed in *Neurospora* similar to previous reports on a wide diversity of ascomycete and basidiomycete fungi [74-76]. We developed a method to assess RIP genome-wide, independent of the underlying genomic content, available in the R package ripr (https://github.com/NBISweden/ripr). This resource can prove useful to identify genomic regions, e.g., centromeres that are TE rich [77, 78], without identification of TEs first, which can be a difficult task depending on the methods used for detection [79, 80]. The emerging pattern from the present study demonstrating correlation between TEs and RIP will become useful tools in several lines of fungal genome assessments from determining TE content by analyzing a less complicated RIP index. Also, under the assumption that repetitive sequences such as TE frequently accumulate at centromeric regions, RIP screening could facilitate identification of such TE-associated genomic regions.

Detected RIP signatures does not however directly demonstrate RIP activity, as other mutational sources present confounding factor, as observed in retroviruses where A- and T-mutations accumulate following reverse transcription [81, 82]. Additionally, RIP index has been developed from *N. crassa* and dinucleotide contexts in other fungal species can vary, and even include trinucleotide contexts [71, 76, 83], and further tests are needed to establish RIP activity in these species. Comparison of RIP efficiency between fungal species will benefit from experiments to quantify the extent of RIP mutations over a number of sexual generations in *Neurospora* as has been tested in a related fungal species, *Podospora anerina* (unpublished results).

## Conclusions

Genome evolution studies benefit greatly from availability of nearly gapless genome assemblies. Here, we have generated, analyzed, and made available high-quality nearly gapless fungal (*Neurospora*) genome assemblies. In spite of the conserved RIP machinery in *Neurospora*, we see that TEs are likely transposing in *Neurospora* and variation in TE content within genomes, between genomes of the same species, and among species, that contributes to genome size evolution. Importantly, we developed a method to determine the RIP indices across a genome assembly to identify regions rich in TE sequences.

## Methods

### Strains investigated in the study

*Neurospora* genomes used in this study were either generated in house or collected from public databases. All strains sequenced for this project were obtained from the Fungal Genetics Stock Center (http://www.fgsc.net/) and *N. sitophila* W1434, which was provided by Jacobson et al. [41]. All strains used in this study (**Table S1**) are referred to by their FGSC identification numbers, unless otherwise noted (e.g. W1426 and W1434, which were kindly provided by D. J. Jacobson [41]). We used three publicly available, well-annotated *Neurospora* genome assemblies that included *N. crassa* (*N. crassa* OR74A version 12 (FGSC 2489) [29], sequenced by Broad Institute, corrected for the assembly error detected by Galazka et al. [30]), and *N. tetrasperma* (FGSC 2508 *mat A* and FGSC 2509 *mat a* [44]). Other PacBio generated assemblies were previously published [39, 40, 42, 43]. The remaining genomes were generated following previously published PacBio sequencing and assembly protocols [39, 42] and are reported for the first time in this study (**Table S1**).

### TE sequence detection

Repetitive DNA and TEs were identified using RepeatMasker (Version 4.0.8, http://www.repeatmasker.org) with Dfam database of repetitive DNA families obtained 20171107 [84], and RepBase [85], as well as a curated *Neurospora*-specific TE library generated in this study (https://doi.org/10.6084/m9.figshare.c.4310996) [49].

To curate the *Neurospora*-specific TE library, we utilized an existing *Neurospora*-specific repetitive sequence library [46] to identify repetitive sequences (TEs, simple repeats and low complexity sequences) in each of the *Neurospora* assemblies and collected results into and updated TE library that included TEs called from of *N. crassa* [29], *N. discreta* (http://genome.jgi-psf.org/Neudi1/) and *N. tetrasperma* (http://genome.jgi-psf.org/Neute1/) and *Neurospora* species sister to the clade containing *N. discreta* [46]. In summary, the original TE library [46] was compiled with RepeatModeler and LTRharvest, which yielded a total 978 repeats (http://fungalgenomes.org/public/neurospora/data/repeatlib/Gioti_neurospora_repeats.renamed.lib). The TE library was further curated manually because many previously LTRharvest-identified elements were incorrect. All 978 elements were re-assessed, including 274 sequences that were previously identified by LTRharvest, and 566 sequences that were classified by RepeatModeler as “Unknowns”, which may include multicopy genes. LTRharvest-identified repeats were queried against the conserved domain database (CDD, *v 3.17, 52910 PSSMs e-value setting 0.01.*) to determine the presence of LTR retrotransposon-related protein domains (belonging to e.g. *gag, pol* (including RNaseH, reverse transcriptase, and integrase), and CHROMO domain) [86, 87]. Sequences were kept in the curated TE library if one of these protein domains were present, except CHROMO domains that required presence of an additional *gag* or *pol* sequence. Cellular genes, with or without transposon-related domains, were removed. RepeatModeler-classified “Unknown” repeats were queried against the CDD. Repeats with sequence similarity to cellular genes were removed from the curated TE library. Sequences without similarity matches to transposon-related protein domains (or cellular genes) were kept in the library as these could represent non-autonomous transposable elements such as SINEs normally lacking identified protein domains.

RepeatModeler-derived “DNA”, “LTR”, and “LINE” repeats (n=8, 61, and 23 sequences, respectively) were confirmed for presence of transposase-[DNA] or retrotransposon-[LTR, LINE] related protein domains in CDD. RepeatModeler-derived “SINE” repeats (n=40 sequences) were confirmed in the genomic tRNA database (Data Release 18.1 (August 2019)), using tRNAscan-SE webserver (http://trna.ucsc.edu/tRNAscan-SE/) and SINEBase [88, 89]. SINE repeats with 100% sequence similarity to a tRNA gene were removed from the curated TE library. The non-redundant nucleotide database (blastn [90]) and Repbase (CENSOR: [91, 92]) were also used to identify transposon similarities for a subset of repeat sequences without conclusive results.

The curated TE library used in this study can be accessed at Figshare (https://doi.org/10.6084/m9.figshare.c.4310996) [49]. We analyzed each *Neurospora* assembly using the updated “Gioti_neurospora_repeats.renamed.v2_191004.lib library” (844 TE sequences), the available *Neurospora crassa* TEs from NCBI Genbank (9 TE sequences), as well as additional TEs described by Selker et al. [47] (28 TE sequences) and Wang et al. [48](4 TE sequences) by using RepeatMasker (RepeatMasker version 4.0.8, run with rmblastn version 2.6.0+, and combined database: Dfam_Consensus 20171107 and RepBase Update 20181026). We limited our query library to include TEs from *Neurospora* and thus exclude other fungal repeats reported in RepBase [93]. The RepeatMasker “.tbl” outputs were used to quantify proportions of the assemblies that contained TEs. The RepeatMasker “.out” outputs were used to calculate the composite RIP index for each of the repeat sequences. Fragmented TEs were not stitched together to create intact TEs, and nested TEs were not disentangled. Here we utilized the RepeatMasker repeat class/family categorization (column 11 in the .out file: DNA, LTR, LINE, SINE, Unclassified, Simple repeat and Low complexity).

### Identification of lineage-specific TE insertions

In order to identify lineage-specific TEs, we modified the methodology illustrated in [94] for presence and absence alignments of TE insertion flanking DNA. We considered a TE insertion to be species-specific when the flanking DNA (occupied integration site or presence state) in the first species are found unambiguously in close proximity in the second species (unoccupied pre-integration site or absence state). All the occupied integration sites for which we found multiple pre-integration sites or undetermined orthology (e.g. when the species are too evolutionary far away from one another) were classified as unresolved integration sites.

Identification involves two steps. First, we extracted 500 bp flanking the TE insertions from each *Neurospora* assembly (based on the RepeatMasker annotation) and aligned to all other *Neurospora* assemblies. The alignment was filtered for putative unoccupied pre-integration sites by identifying TE flanking DNA that aligned less than 50 bp apart and with at least 70% similarity (to take RIP mutations into account). Second, we required the putative pre-integration sites to be unique and therefore aligned these to the first species. If the putative pre-integration site mapped uniquely and at least for 80% of the locus length to the TE integration site, the insertion was classified as a species-specific insertion. In case the putative pre-integration sites had ambiguous alignments in the first comparison assembly, the insertion was classified as an unresolved site. Finally, when both TE flanking sequences failed to map confidently, the corresponding insertions were classified as unresolved sites. Scripts can be found in the Github repository https://github.com/ValentinaBoP/NeurosporaSpecificTE.

### TE distribution in the genome

To determine the distribution of TEs along the *Neurospora* genomes, we calculated log2-ratios of observed to expected TE content in 10-kb non-overlapping sliding windows. For each window, observed TE content was calculated by summing the intersection of the window coordinates with the TE coordinates, as defined in the RepeatMasker output file “.out”. The expected TE content was calculated as the window size times the genome-wide TE fraction.

### Composite RIP index calculations

RIP indices were calculated for each of the transposon fragments. Frequencies of TpA, ApT, CpA, TpG, ApC, and GpT dinucleotides in each of the fragments were tabulated in the RepeatMasker outputs “.out” file; sequences were extracted using the EMBOSS seqret function [95]. From these frequencies, the ratios TpA/ApT (“RIP product index”) and (CpA+TpG)/(ApC+GpT) (“RIP substrate index”) and the composite RIP index [(TpA/ApT) - (CpA+TpG)/(ApC+GpT)] were calculated [50, 96]. TEs considered to have experienced RIP mutations are described as TpA/ApT > 0.89 and (CpA+TpG)/(ApC+GpT) < 1.03, as well as positive values of the composite RIP index [29, 50, 96].

To assess the significance of composite RIP scores for TEs in a *Neurospora* genome, we generated mock sequences on which to calculate null distributions of RIP scores. Mock sequences were generated in two ways, either by shuffling all genome positions or drawing nucleotides at random from the observed nucleotide frequency distribution, preserving genome length in both cases. For each panel in **Figure S4**, three distributions were plotted. First, the observed RIP distribution for a genome was calculated using the repeats determined by RepeatMasker (“obs”). Second given the positions of the repeats in the genome, RIP scores were recalculated by shuffling the repeat positions for two different cases. A random genome was obtained by shuffling the bases (“shuffle”), and third, generated by sampling nucleotides from the observed nucleotide frequency distribution (“frequency”), Cases 2 and 3 will disrupt all repeat regions such that no RIP signal should be observed. Functions to calculate RIP were implemented in R and are available in the R package ripr (https://github.com/NBISweden/ripr).

The correlation between windows with RIPed sequences and windows enriched in TEs were visualized with the segmented package (version 1.2-0) in R (version 3.6.0) [97, 98]. Linear regression models were estimated with two segmented relationships. Estimates of the slopes and breakpoints are provided. The psi (“psi1”) value indicates the breakpoint. The adjusted R^2^ (“adjusted R”) value gives an indication of how well the linear models fit the data, adjusted for the number of parameters, and considers both regression lines.

## Declarations

### Ethics approval and consent to participate

Not applicable

### Consent for publication

Not applicable

### Availability of data and materials

All previously unpublished raw sequencing reads generated in this study have been deposited at the Sequence Read Archive as BioProject PRJNA622402. All genome assemblies based on PacBio sequencing and reference assemblies, and RepeatMasker tables of repeats for all genomes (“.tbl” and “.out” files) and the curated *Neurospora* TE library have been deposited at Figshare (https://doi.org/10.6084/m9.figshare.c.4310996) [49]. The pipeline to determine RIP and TE landscape patterns, and RIP background are contained in an R script also deposited in Figshare.

### Completing interests

The authors declare that they have no competing interests.

### Authors’ contributions

DN, VP, PU analyzed data and all authors participated in data and results interpretation. DN and HJ drafted the manuscript and VP, PU, AS, PJ contributed to writing the manuscript. All authors read and approved the final manuscript.

## Acknowledgements

Magdalena Grudzinska-Sterno for preparing the strains for sequencing. Mats Pettersson and Douglas Scofield for scripts and discussion, Björn Nystedt at the National Bioinformatics Infrastructure Sweden at SciLifeLab and the Uppsala TE Jamboree group for helpful discussions.

