## Supplementary figures and images for "Transposon- and genome dynamics in the fungal genus *Neurospora*: insights from nearly gapless genome assemblies"

### Figure S1

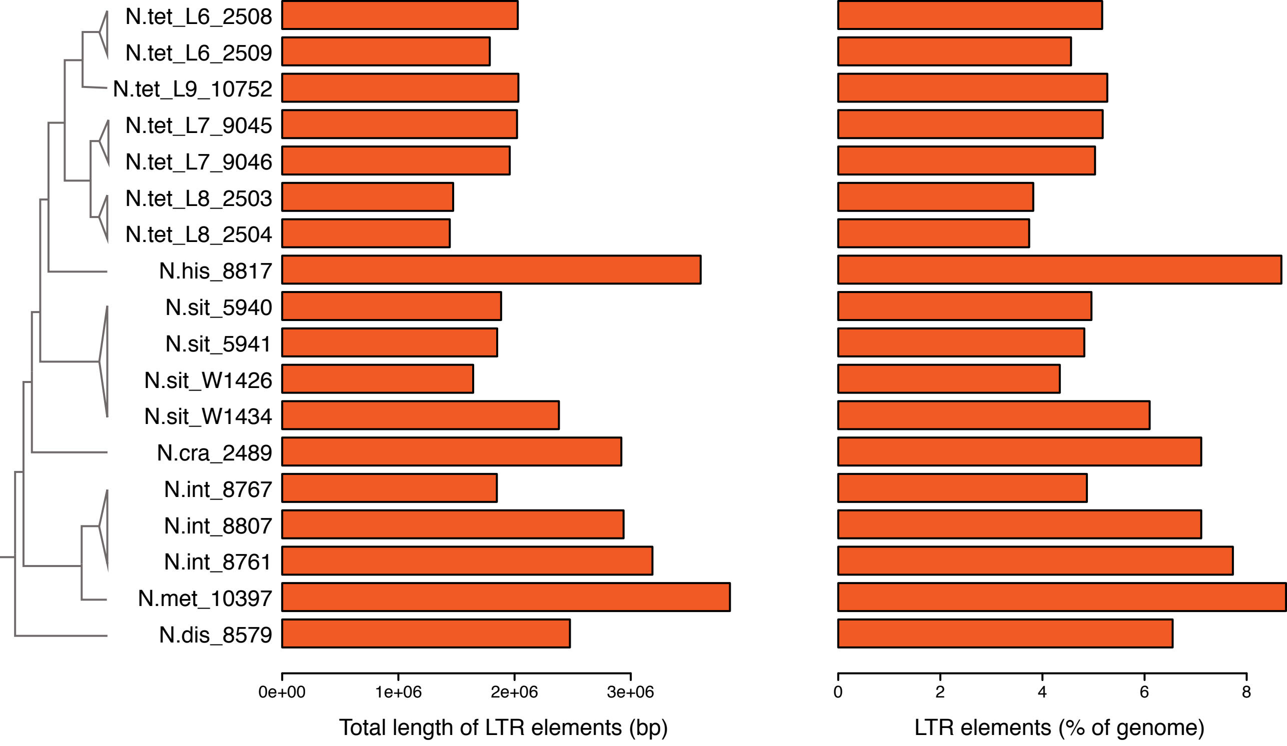

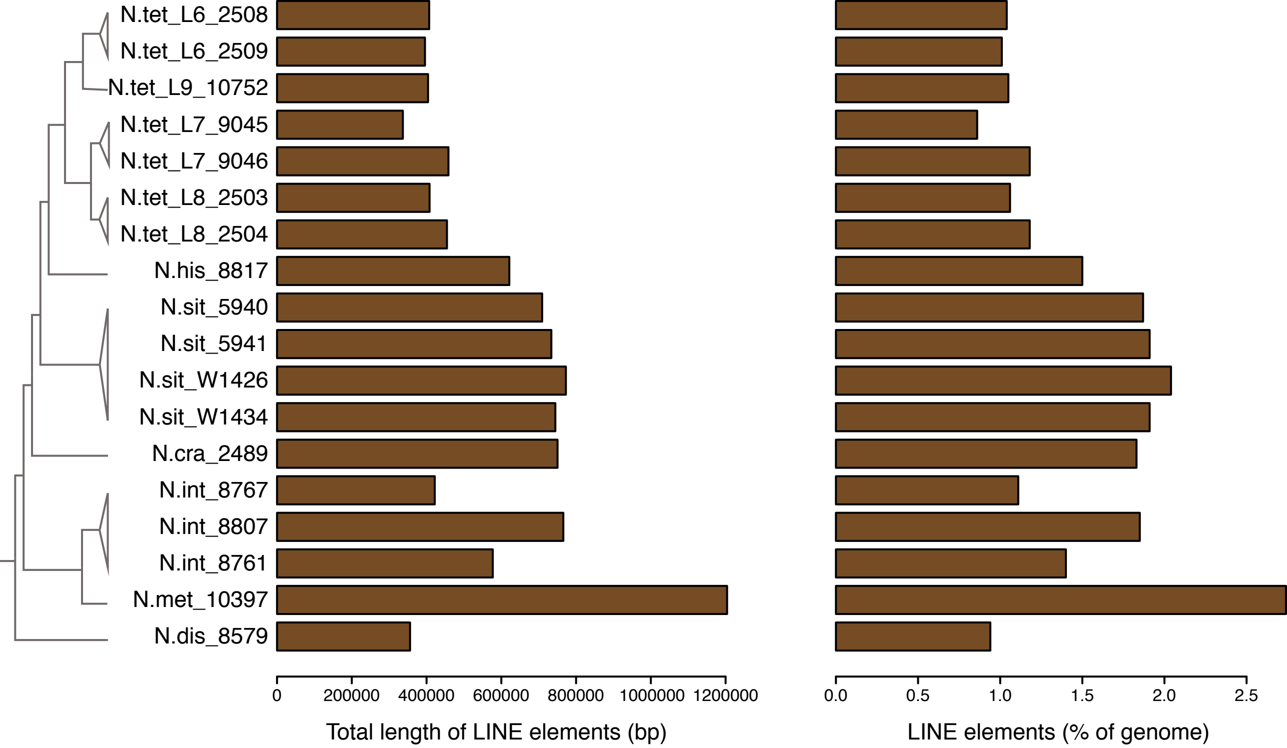

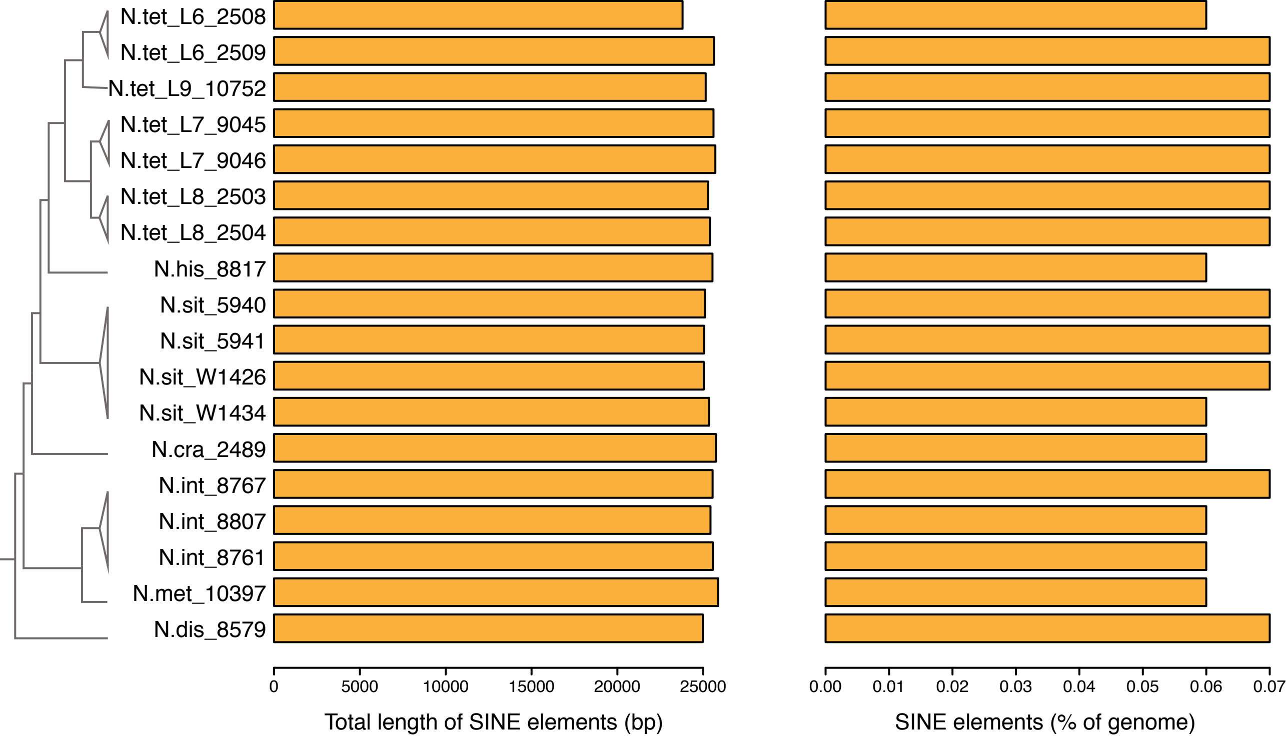

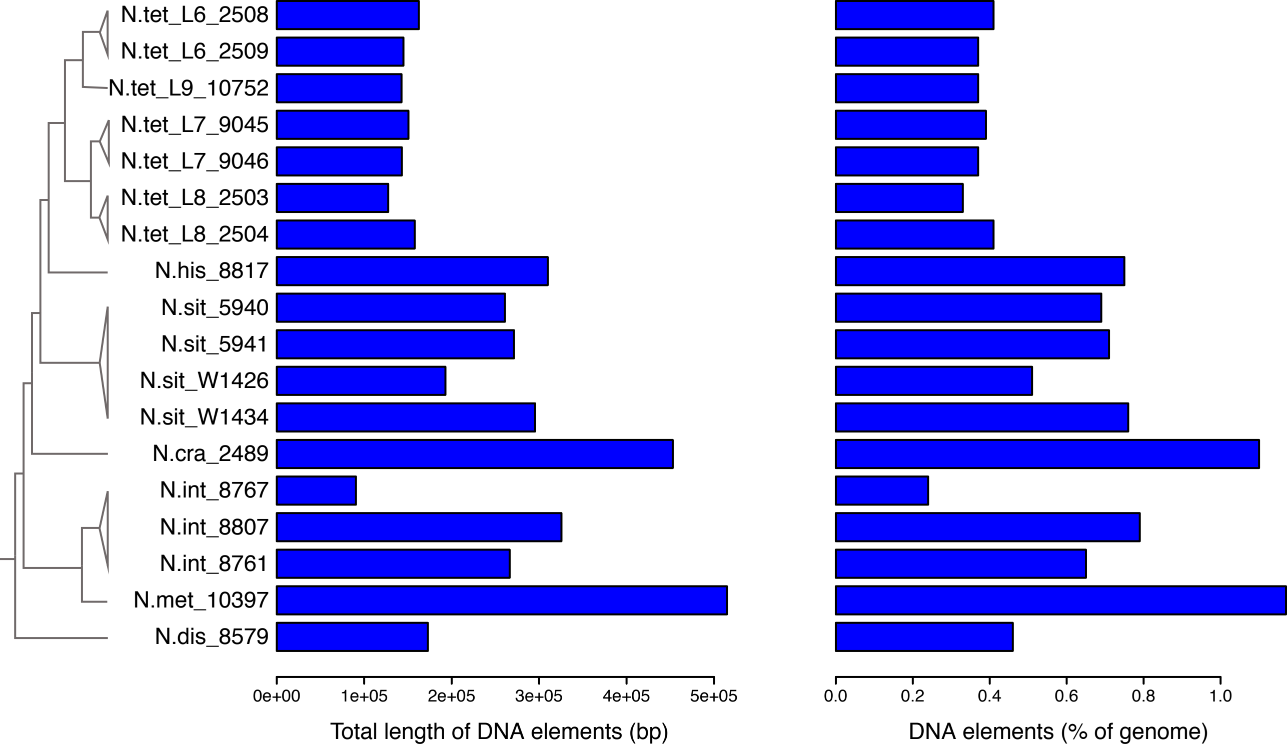

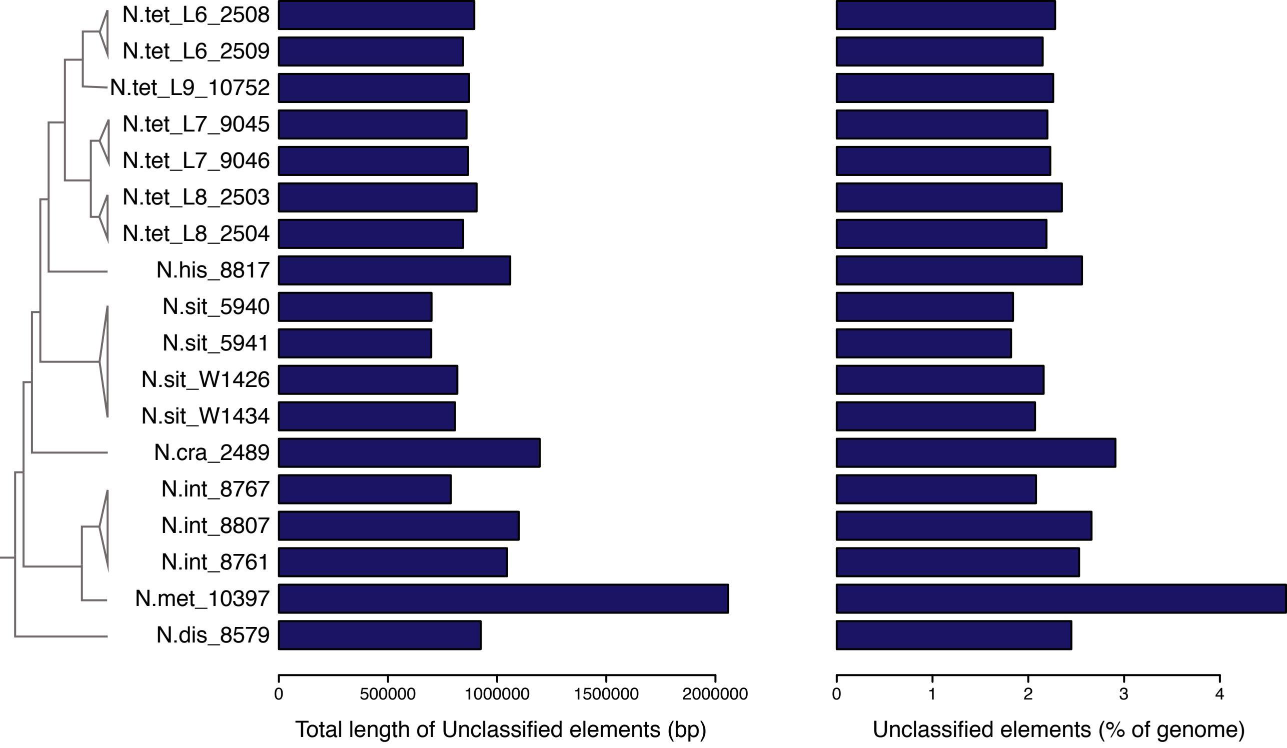

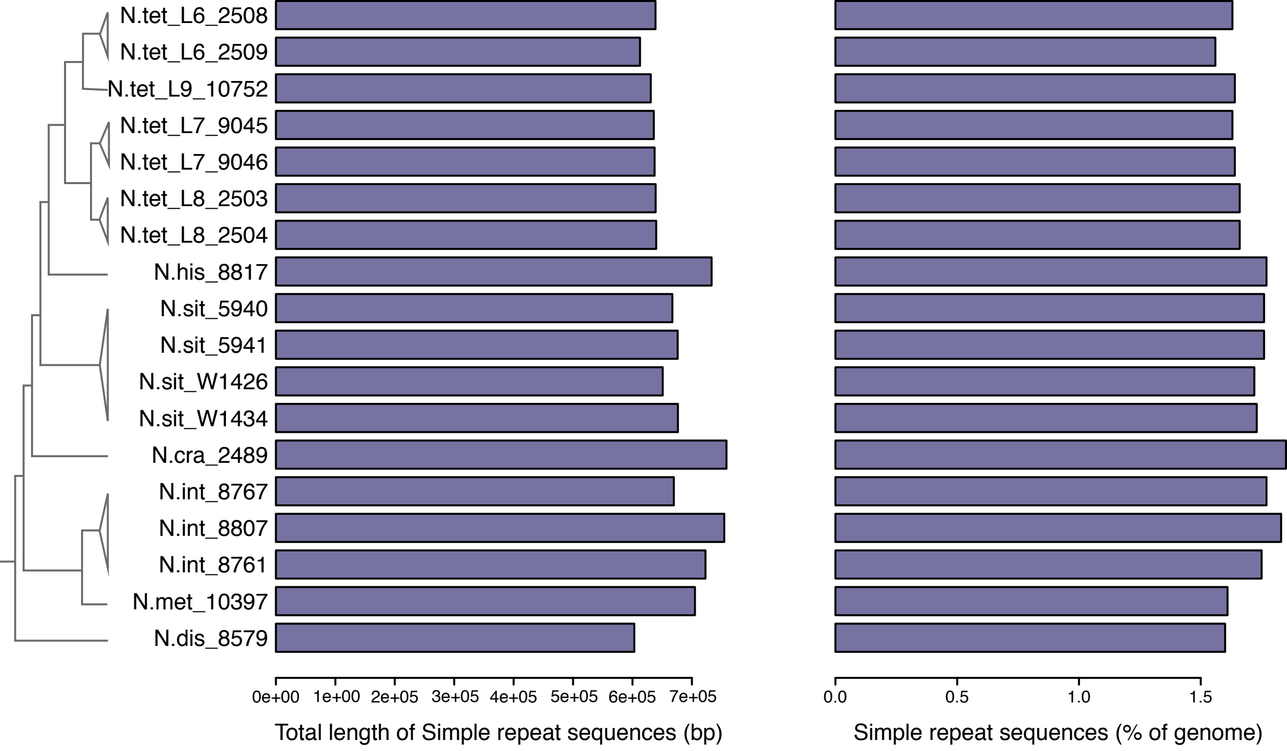

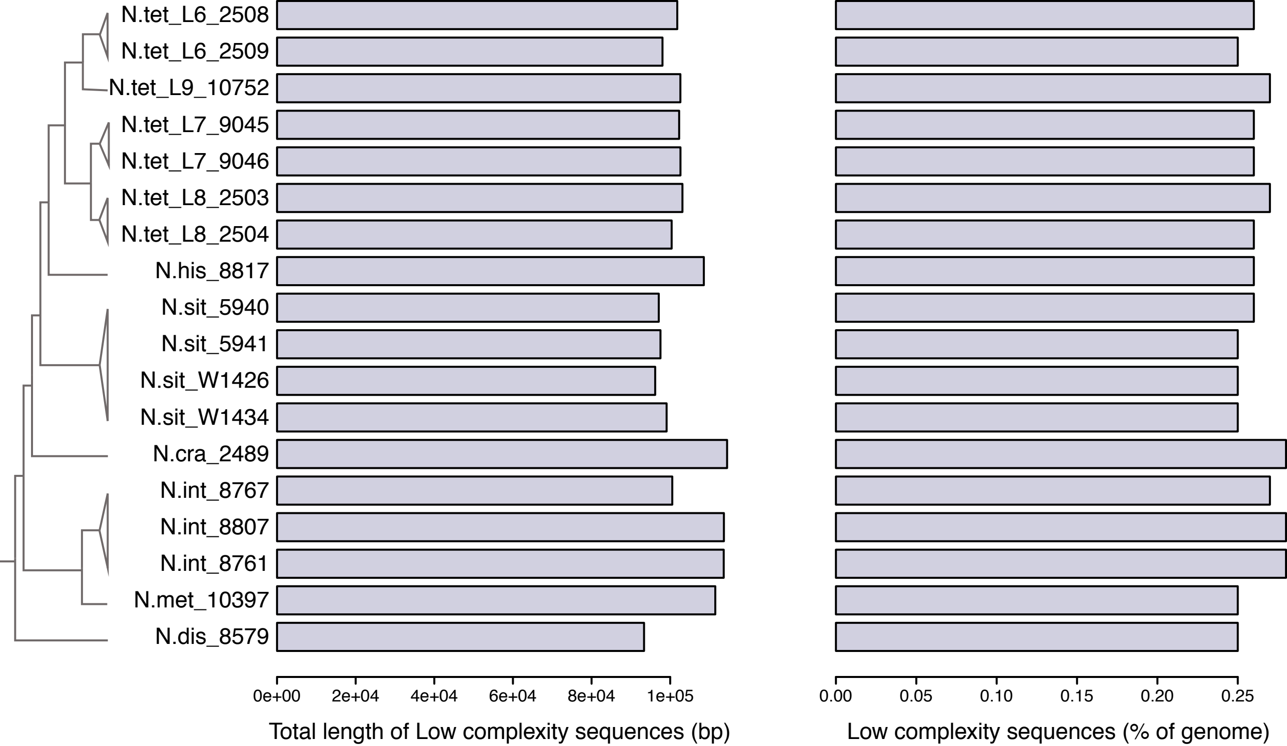

### Figure S2

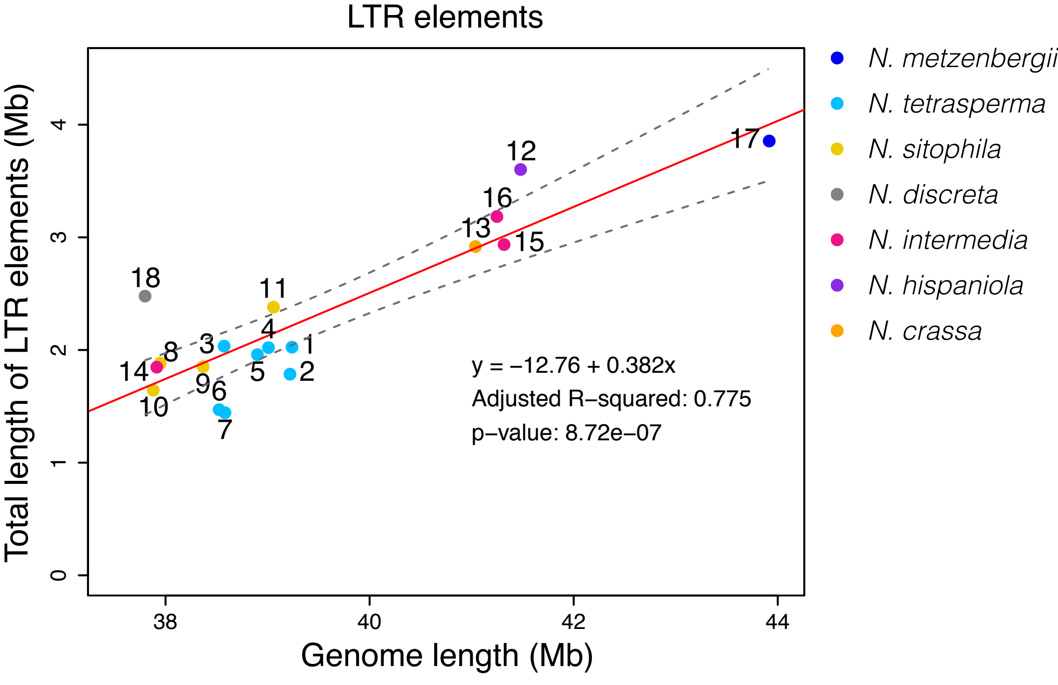

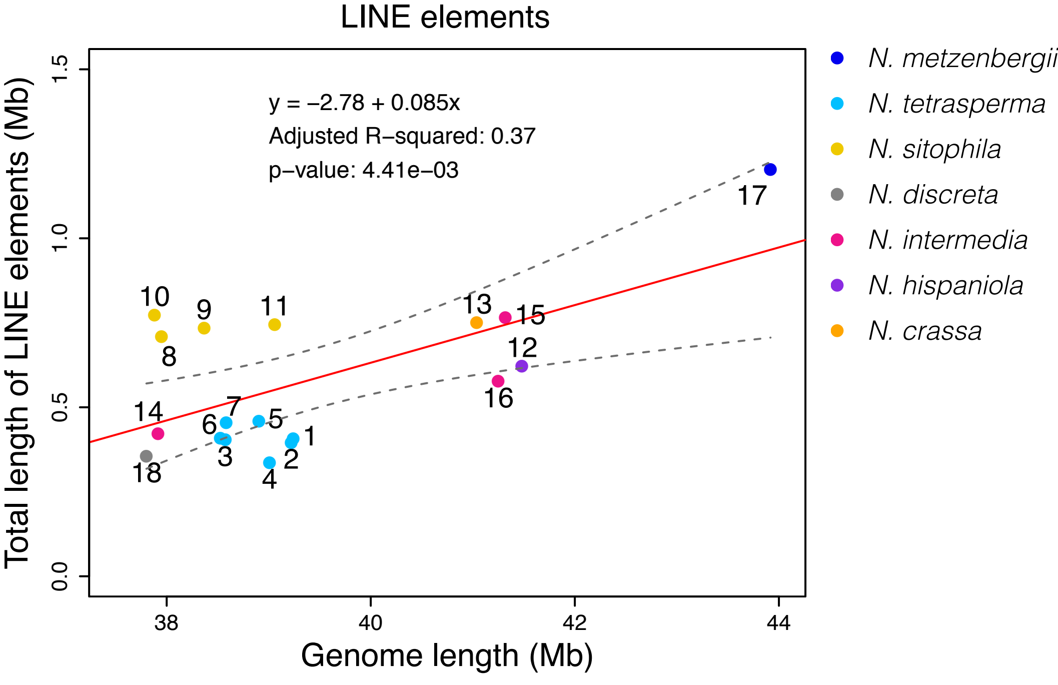

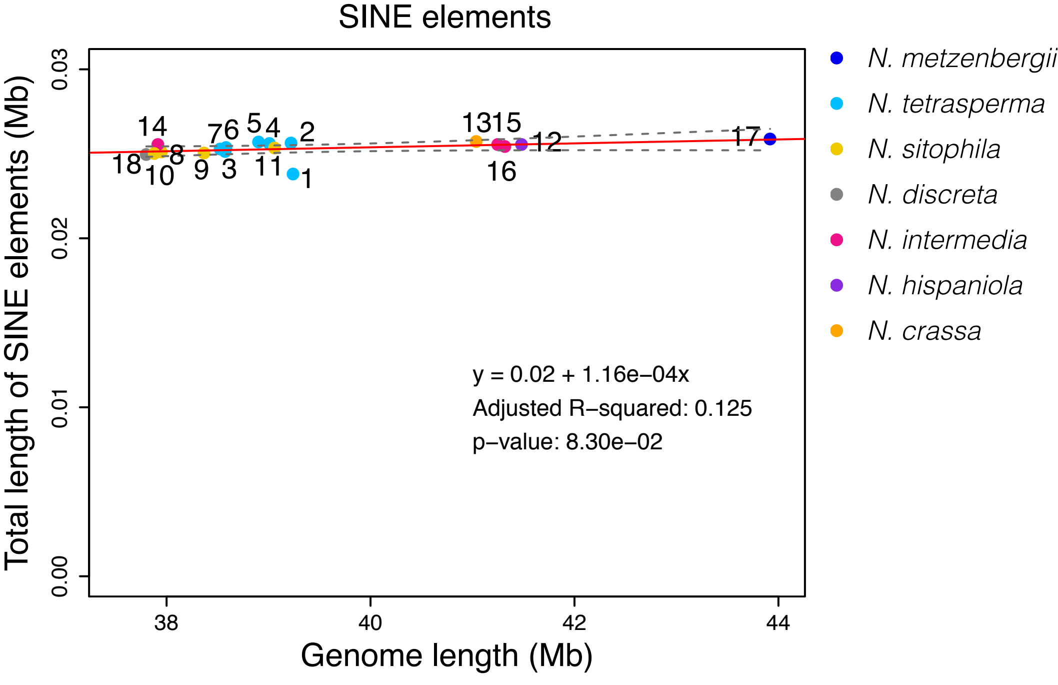

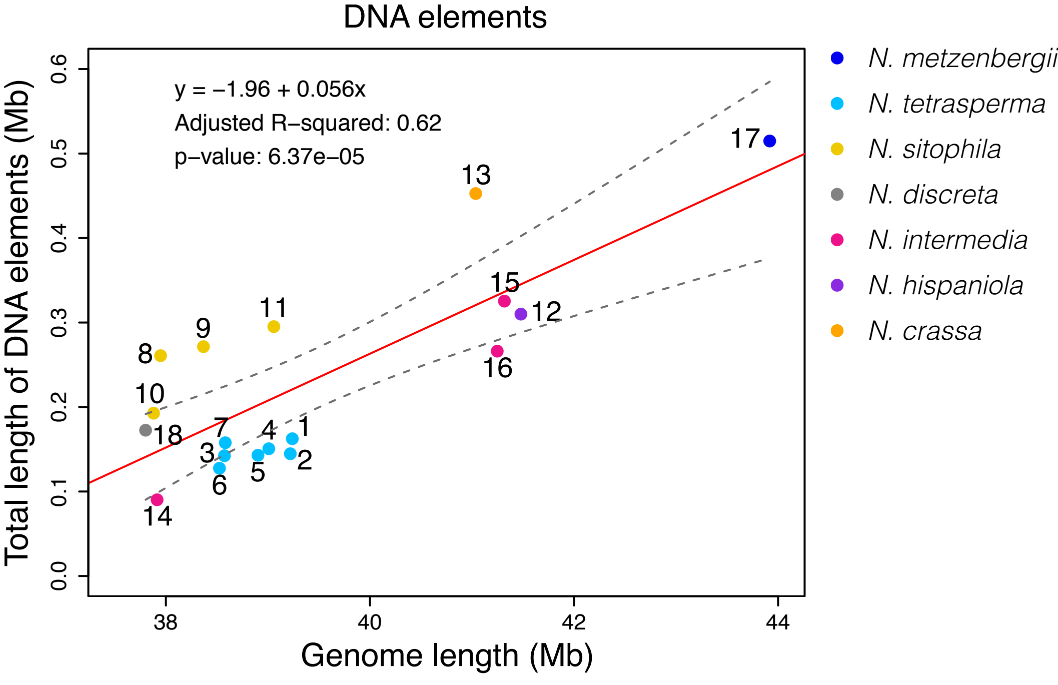

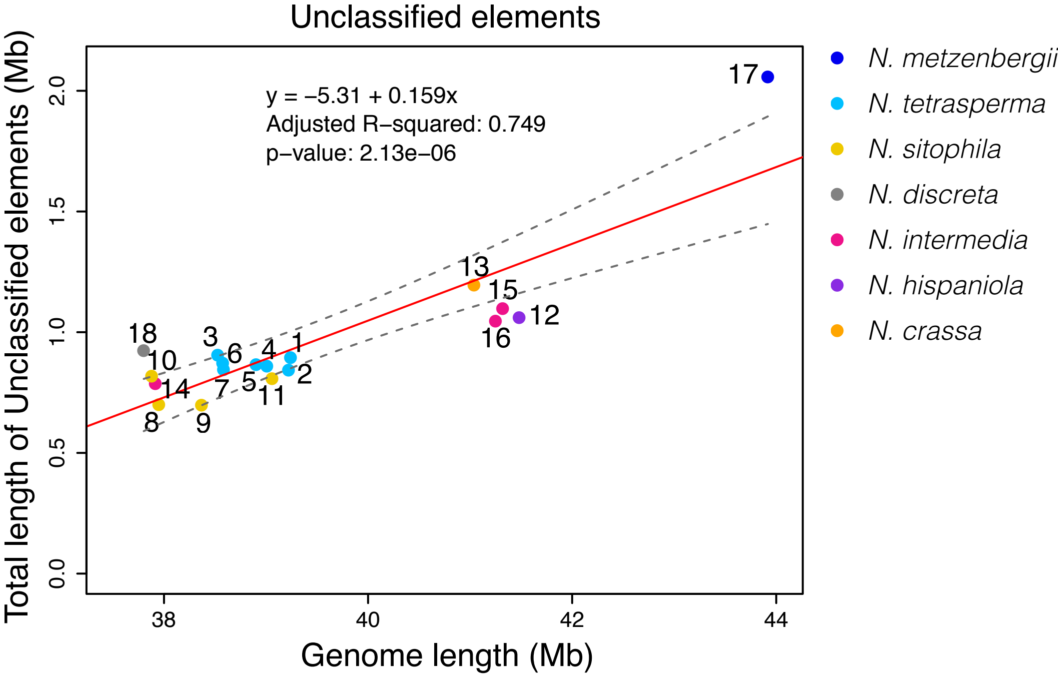

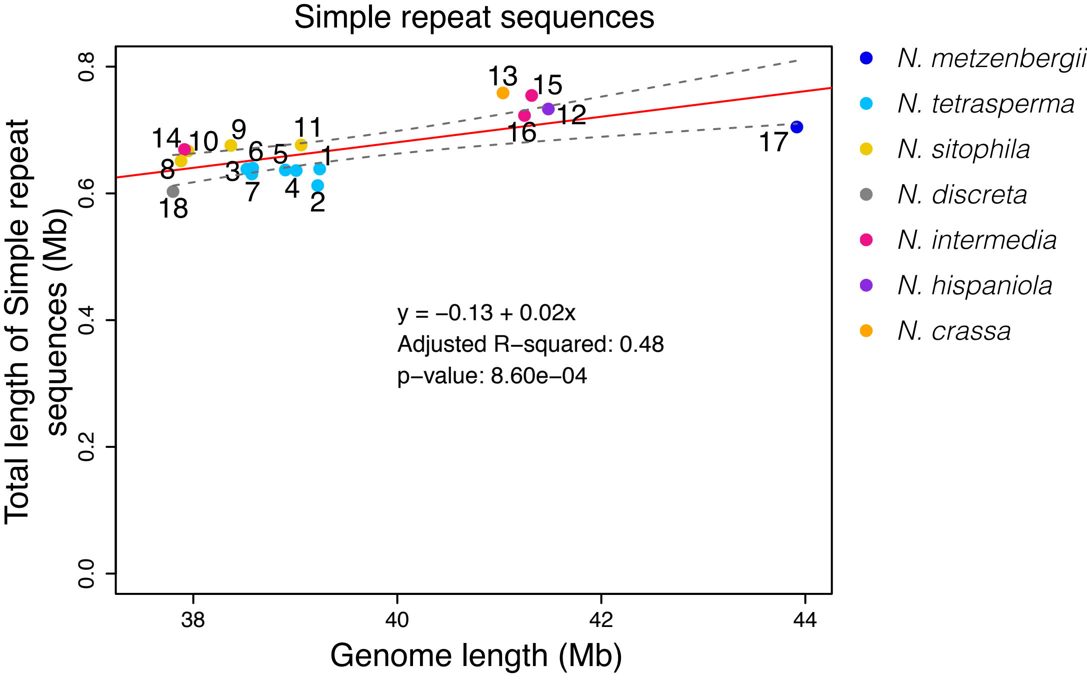

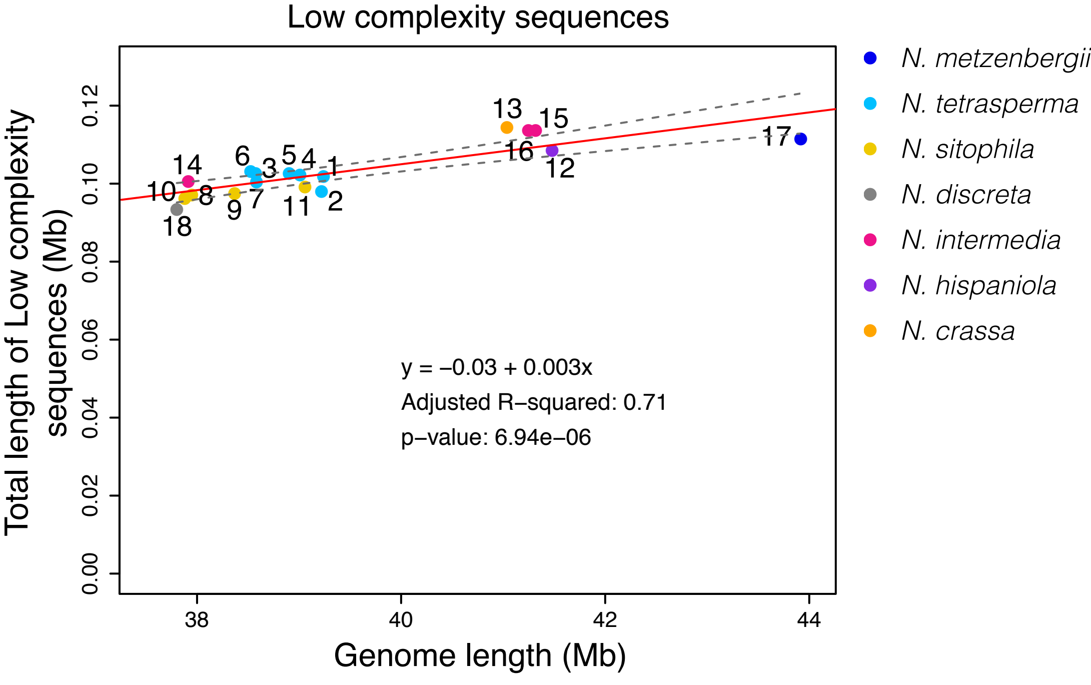
