## Supplementary material for "Transposon- and genome dynamics in the fungal genus *Neurospora*: insights from nearly gapless genome assemblies": Figure S3

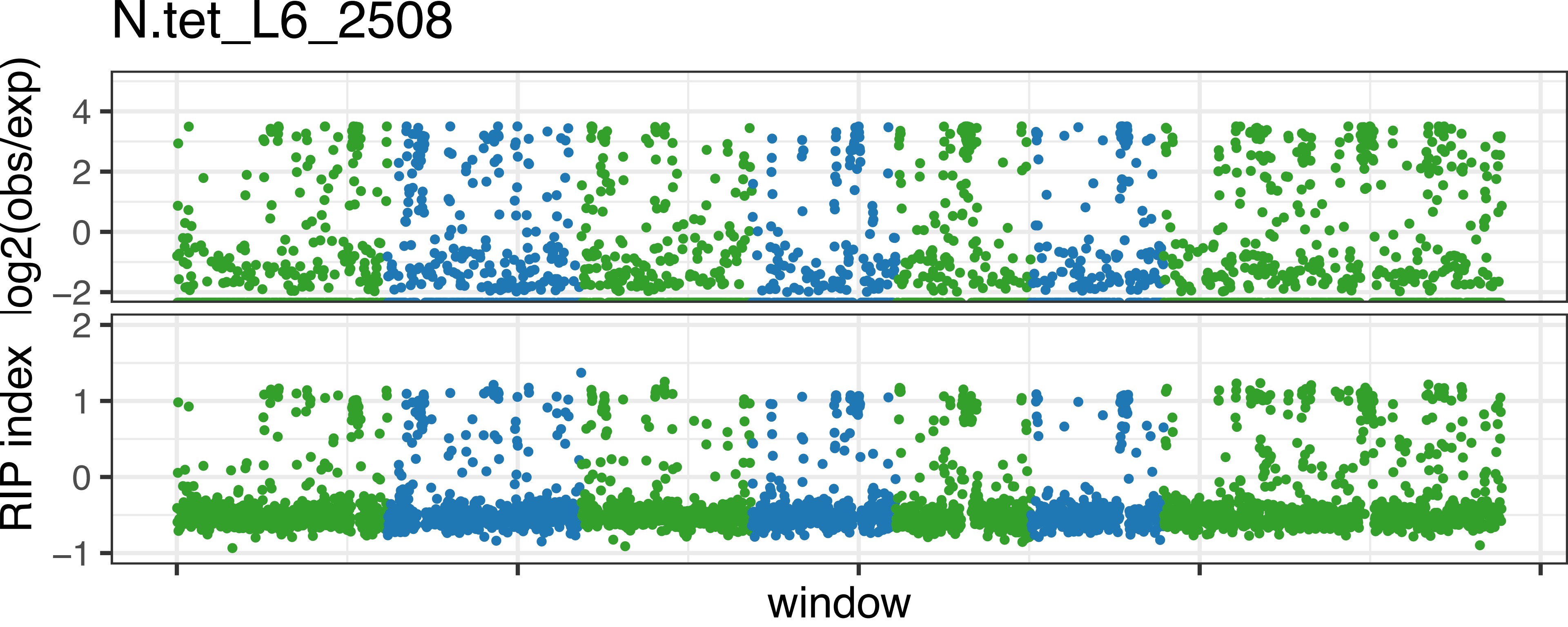
N.tet_L6_2508
log2(obs/exp)
4
2
0
−2
RIP index
2
1
0
−1
window

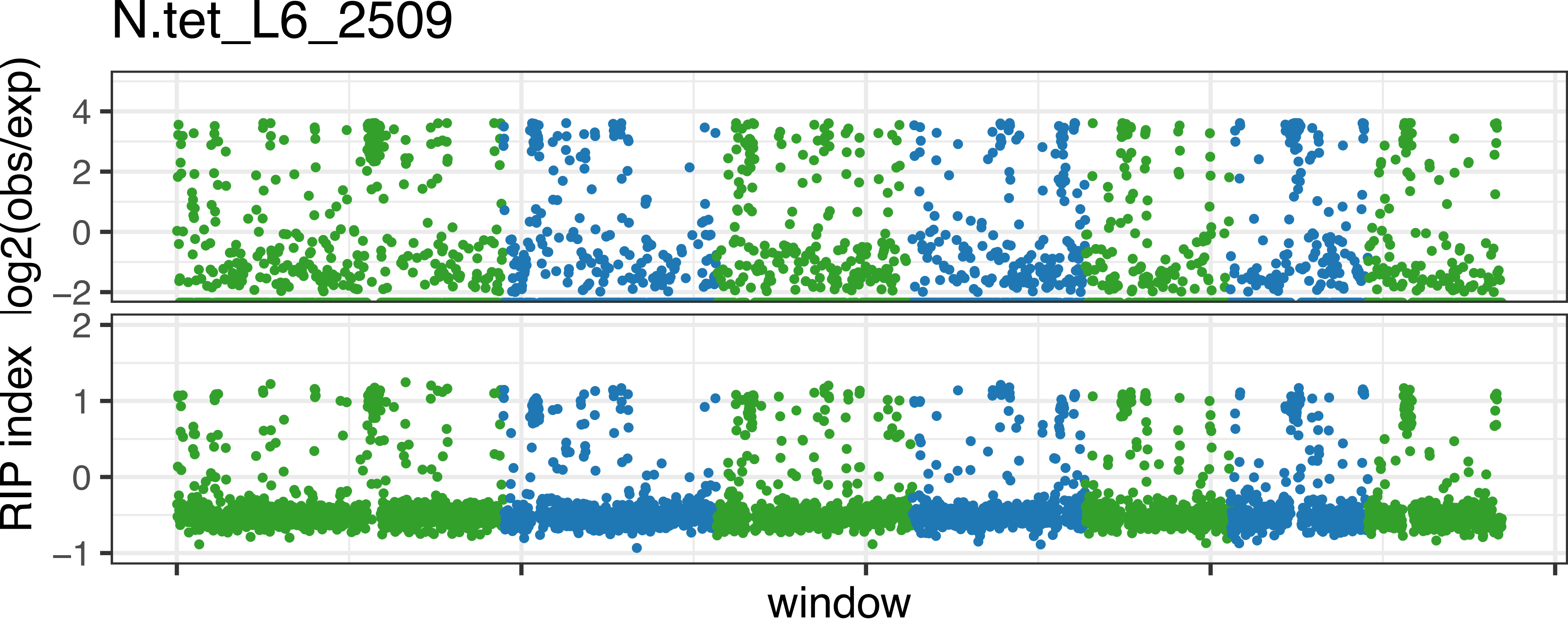
N.tet_L6_2509
log2(obs/exp)
4
2
0
−2
RIP index
2
1
0
−1
window

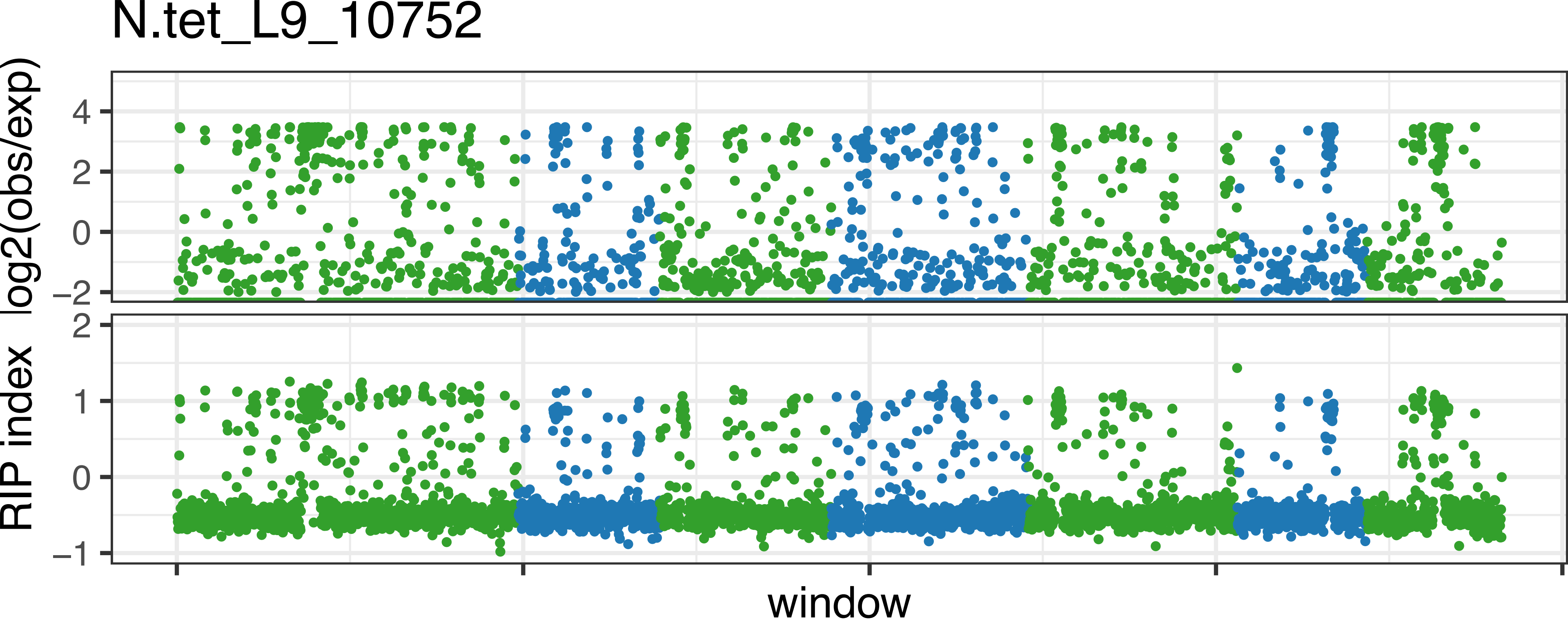
N.tet_L9_10752
log2(obs/exp)
4
2
0
−2
RIP index
2
1
0
−1
window

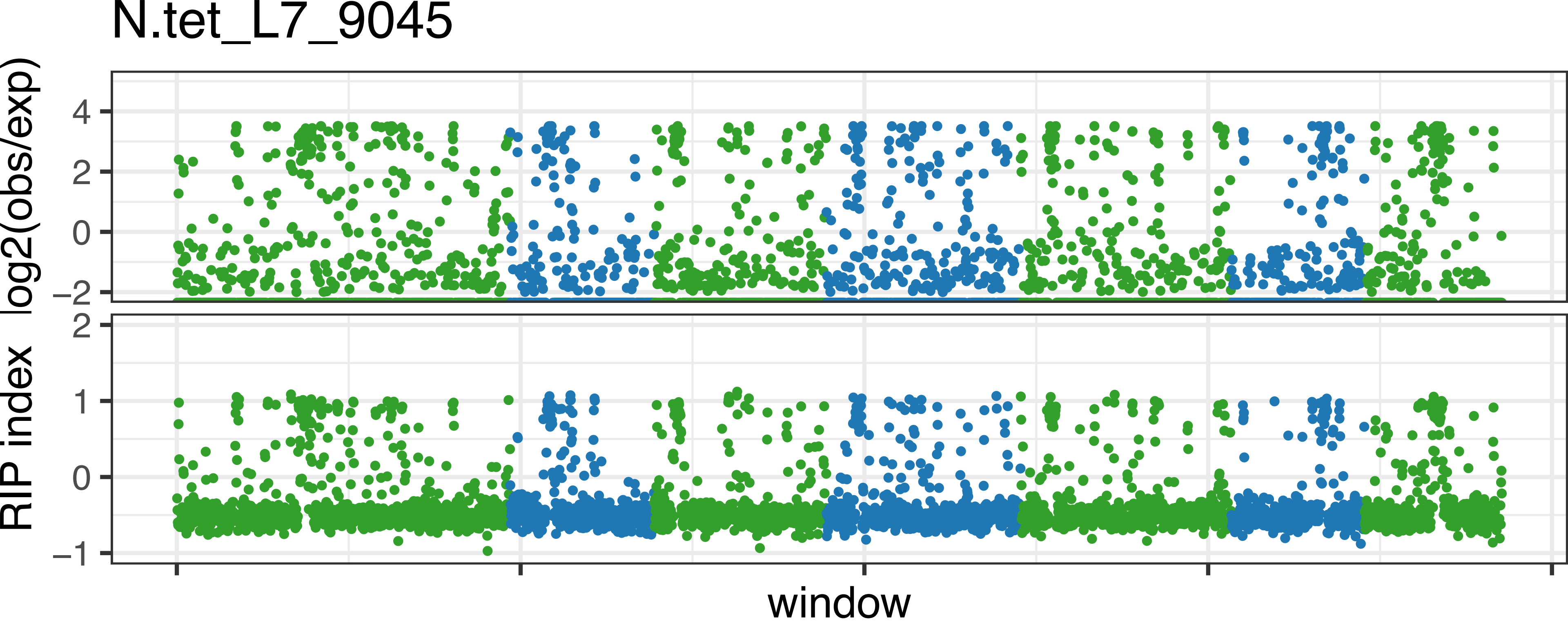
N.tet_L7_9045
log2(obs/exp)
4
2
0
−2
RIP index
2
1
0
−1
window

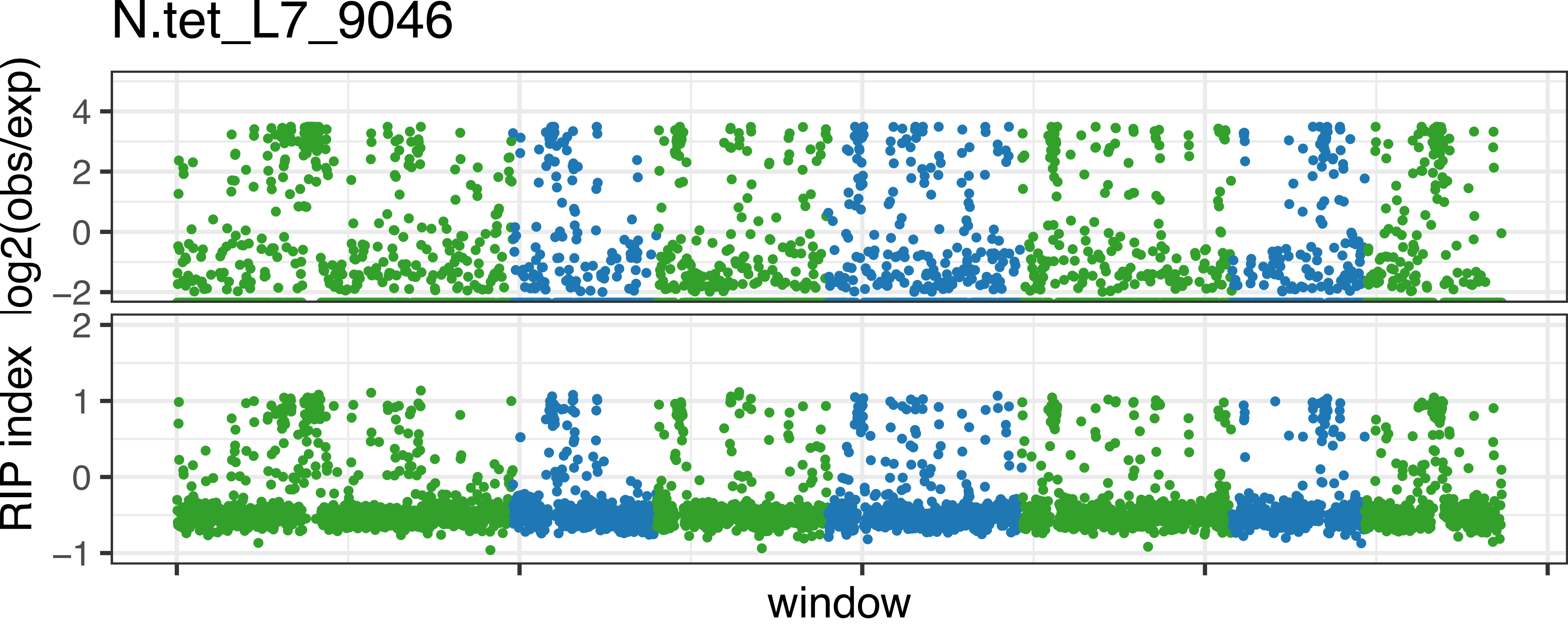

N.tet_L7_9046
log2(obs/exp)
4
2
0
−2
RIP index
2
1
0
−1
window

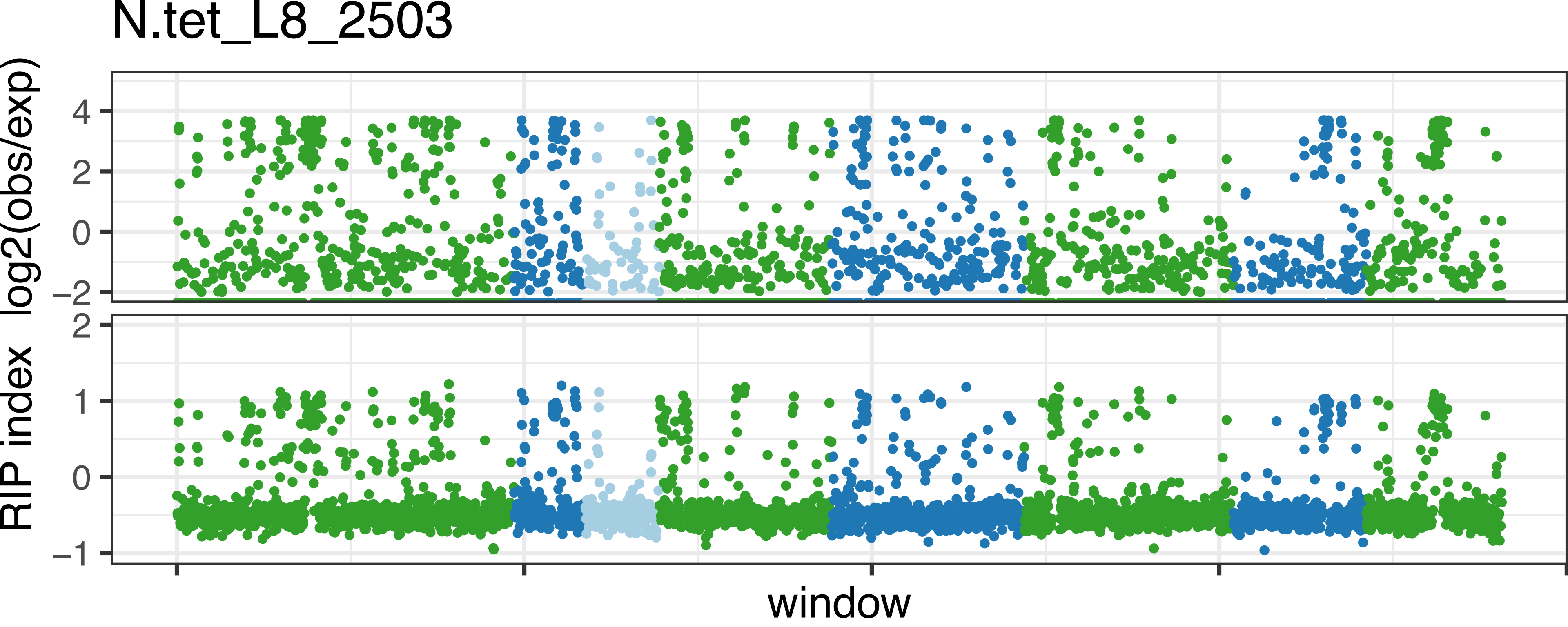

N.tet_L8_2503
log2(obs/exp)
4
2
0
−2
RIP index
2
1
0
−1
window

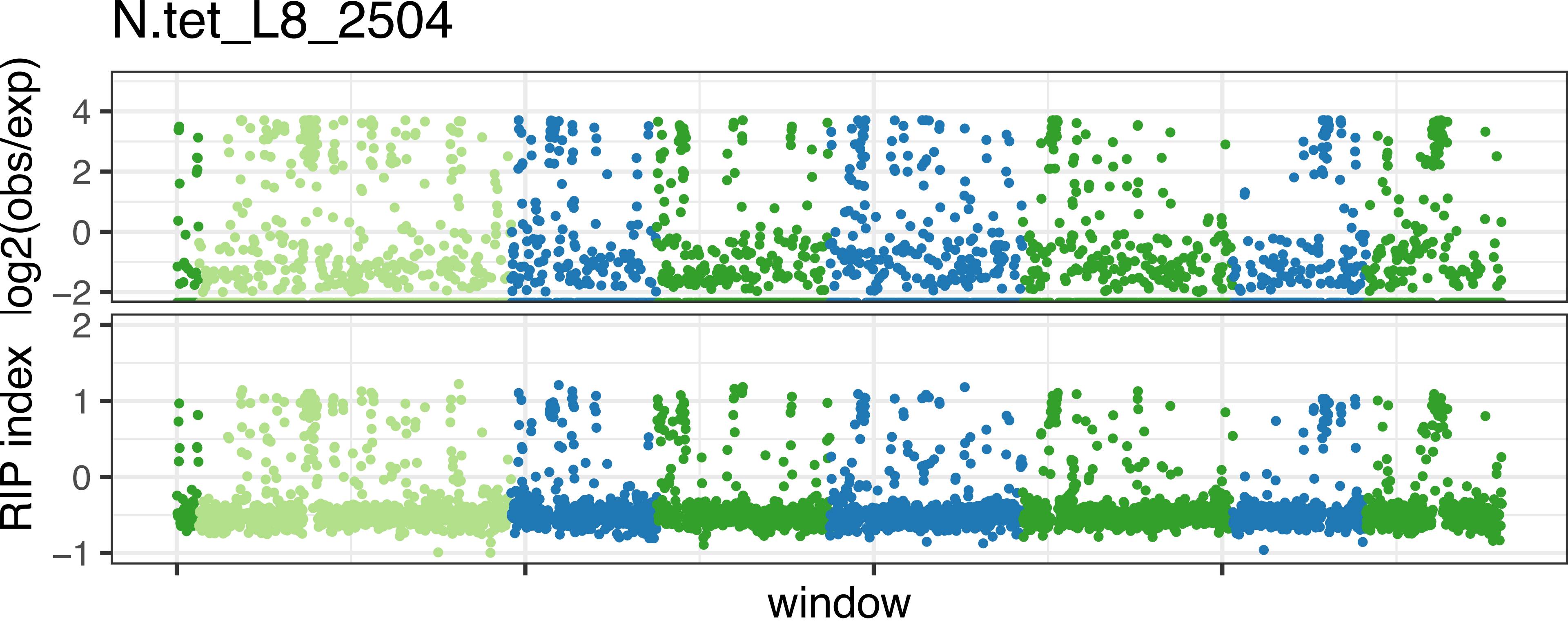
N.tet_L8_2504
log2(obs/exp)
4
2
0
−2
RIP index
2
1
0
−1
window

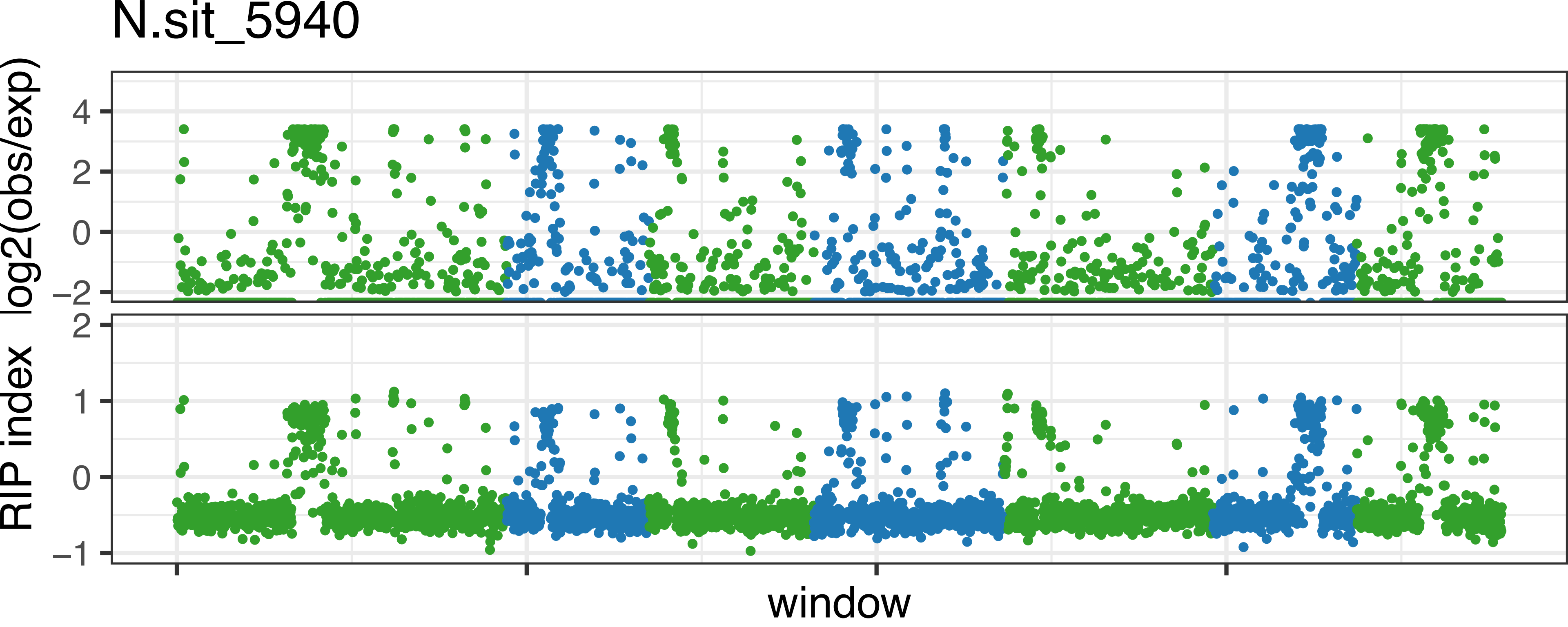
N.sit_5940
log2(obs/exp)
4
2
0
−2
RIP index
2
1
0
−1
window

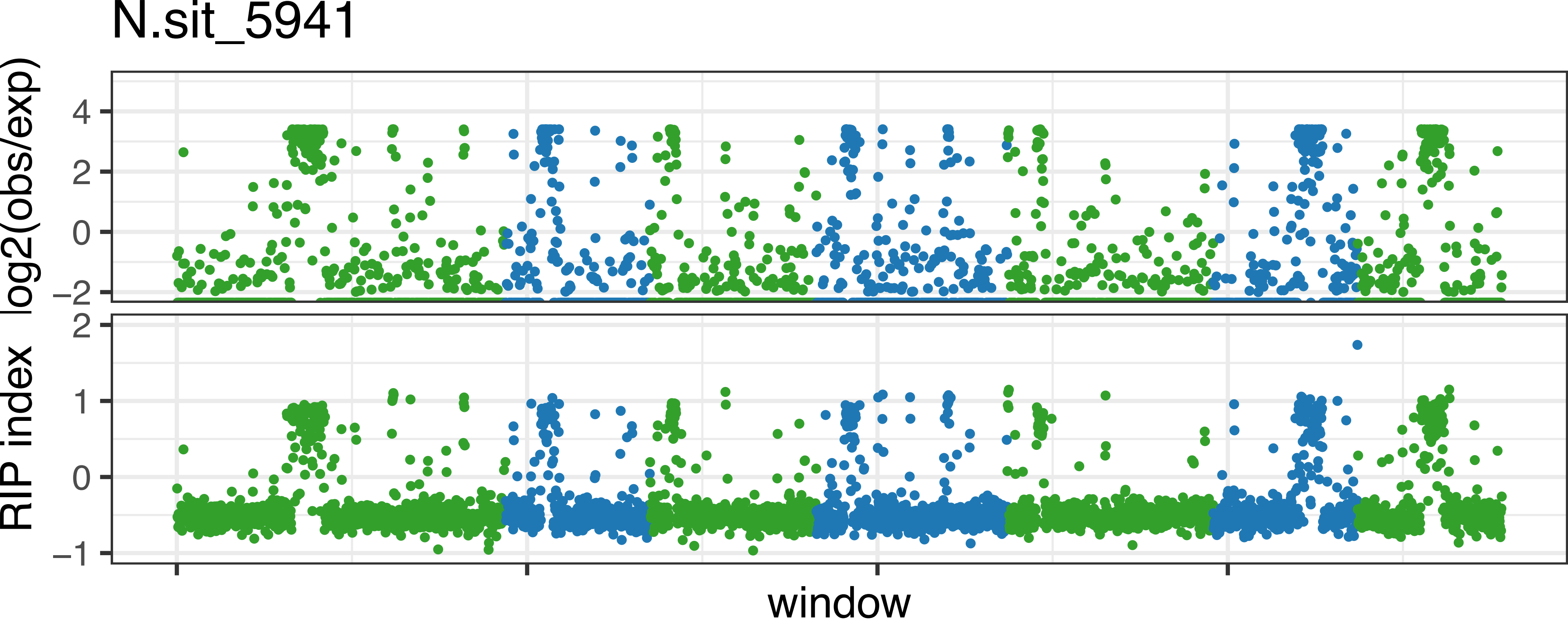
N.sit_5941
log2(obs/exp)
4
2
0
−2
RIP index
2
1
0
−1
window

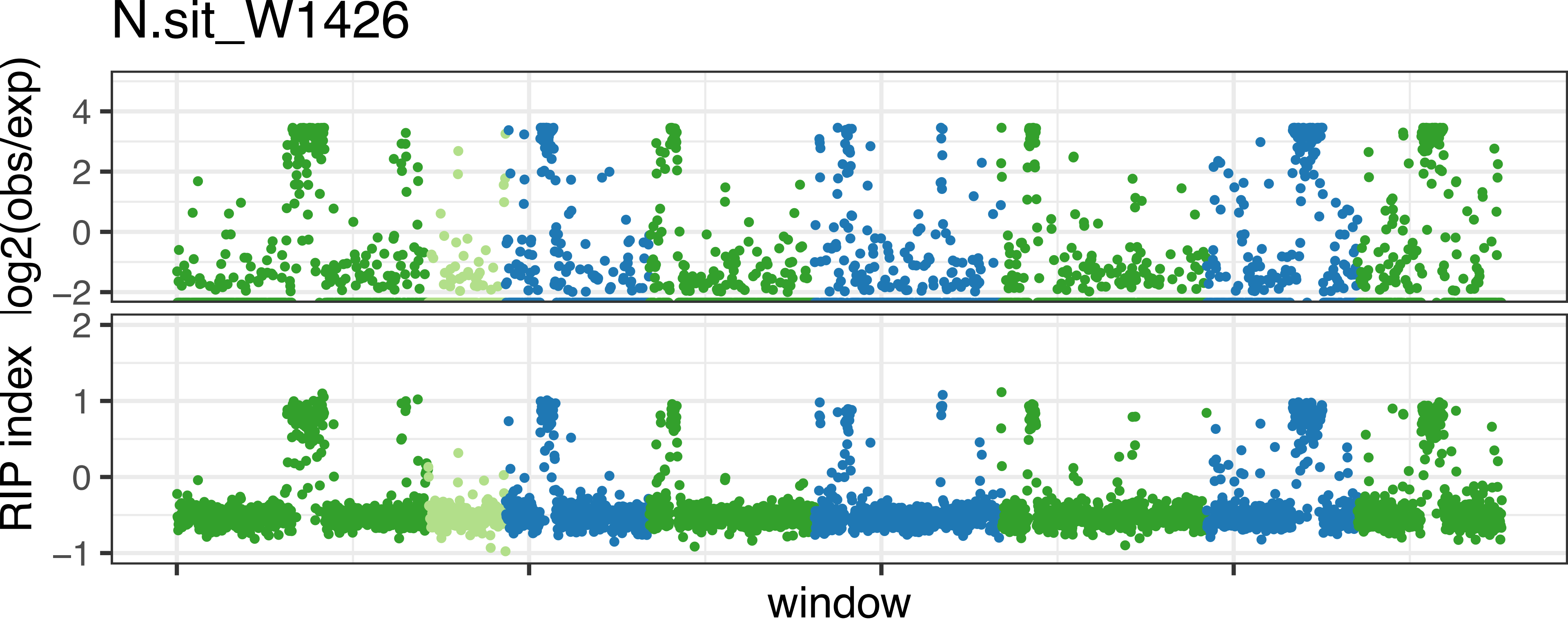

N.sit_W1426
log2(obs/exp)
4
2
0
−2
RIP index
2
1
0
−1
window

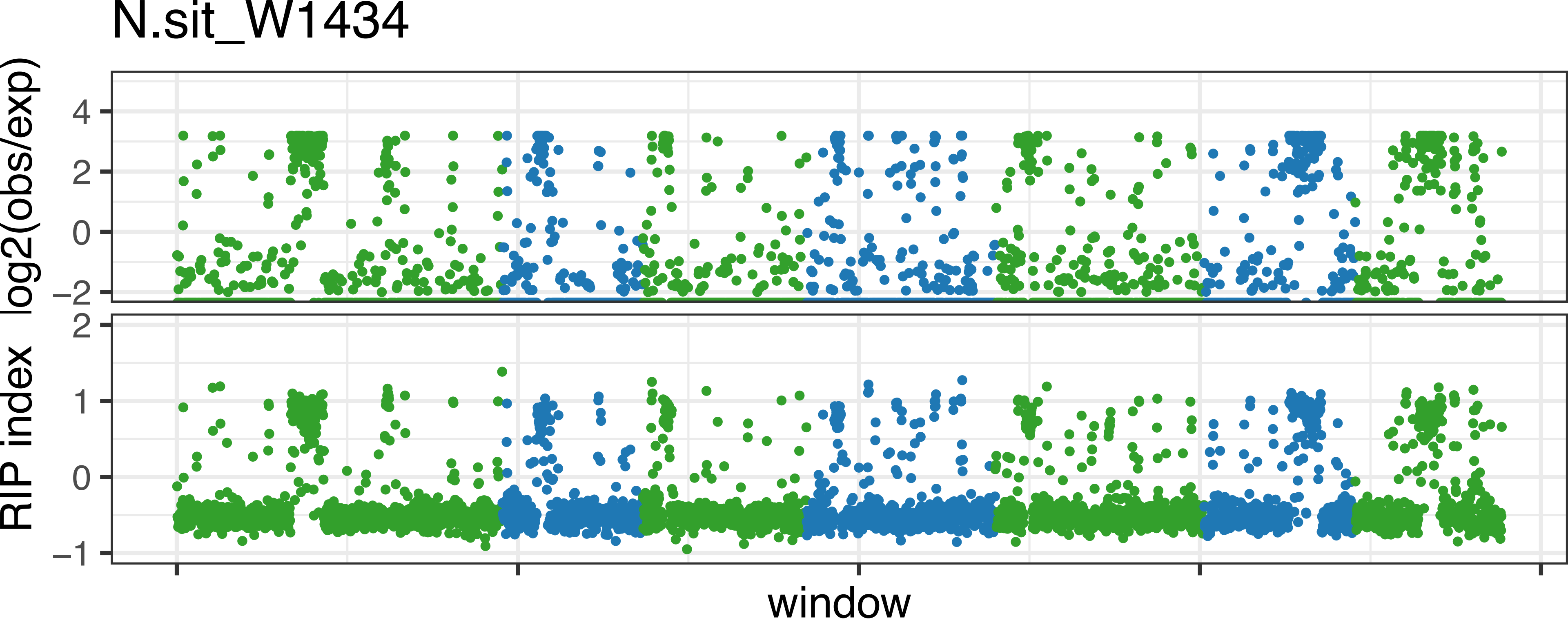
N.sit_W1434
log2(obs/exp)
4
2
0
−2
RIP index
2
1
0
−1
window

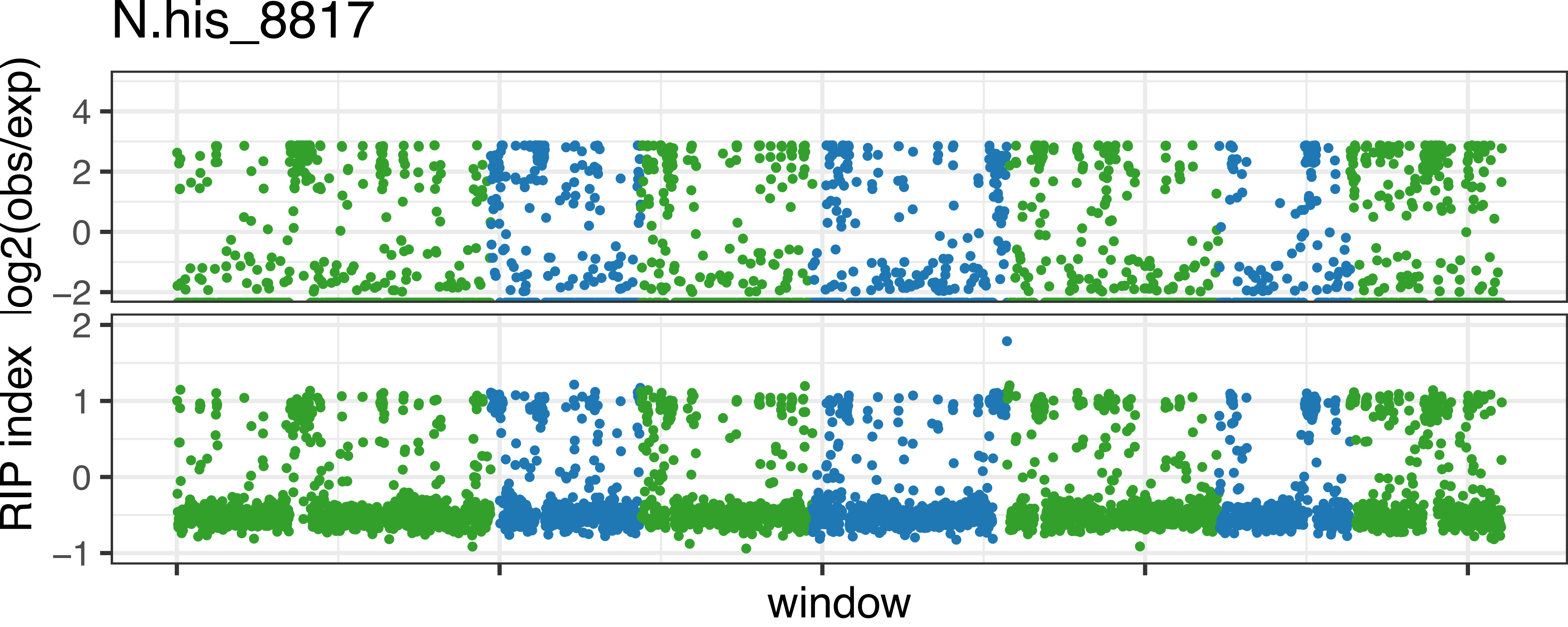

N.his_8817
log2(obs/exp)
4
2
0
−2
RIP index
2
1
0
−1
window

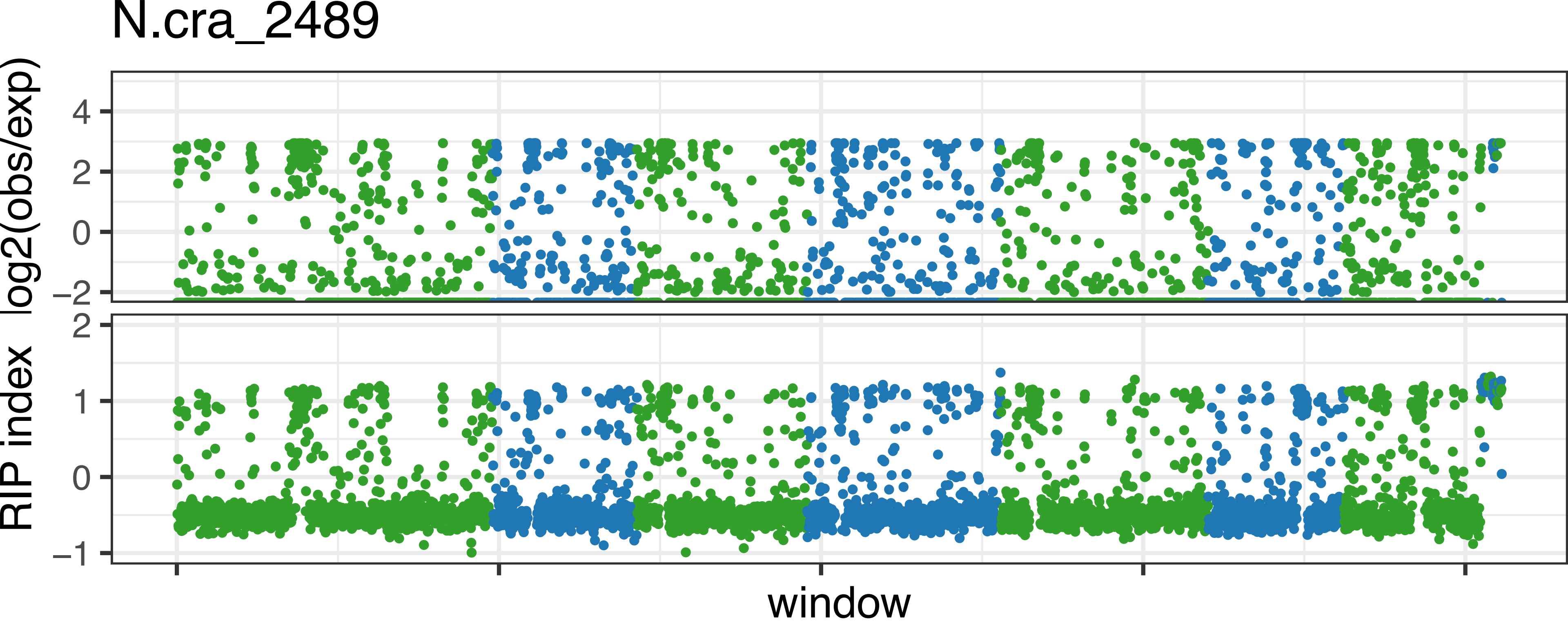

N.cra_2489
log2(obs/exp)
4
2
0
−2
RIP index
2
1
0
−1
window

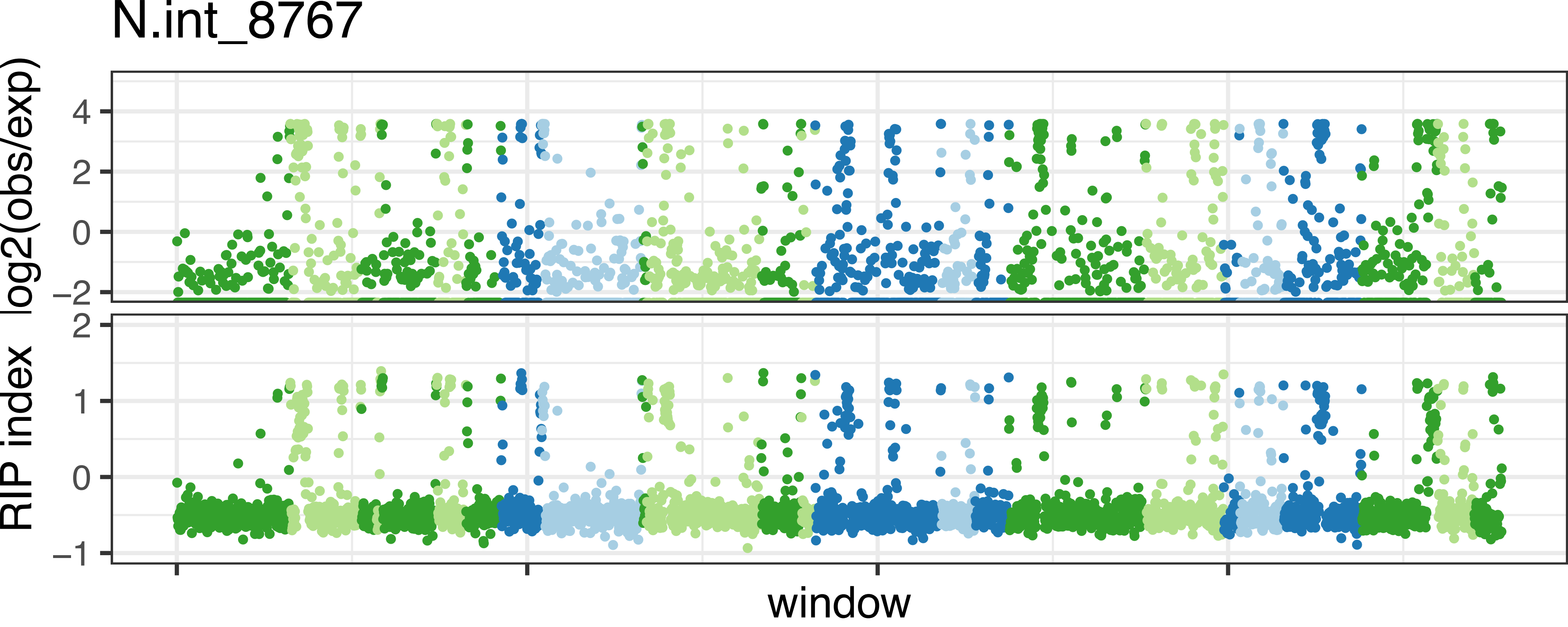

N.int_8767
log2(obs/exp)
4
2
0
−2
RIP index
2
1
0
−1
window

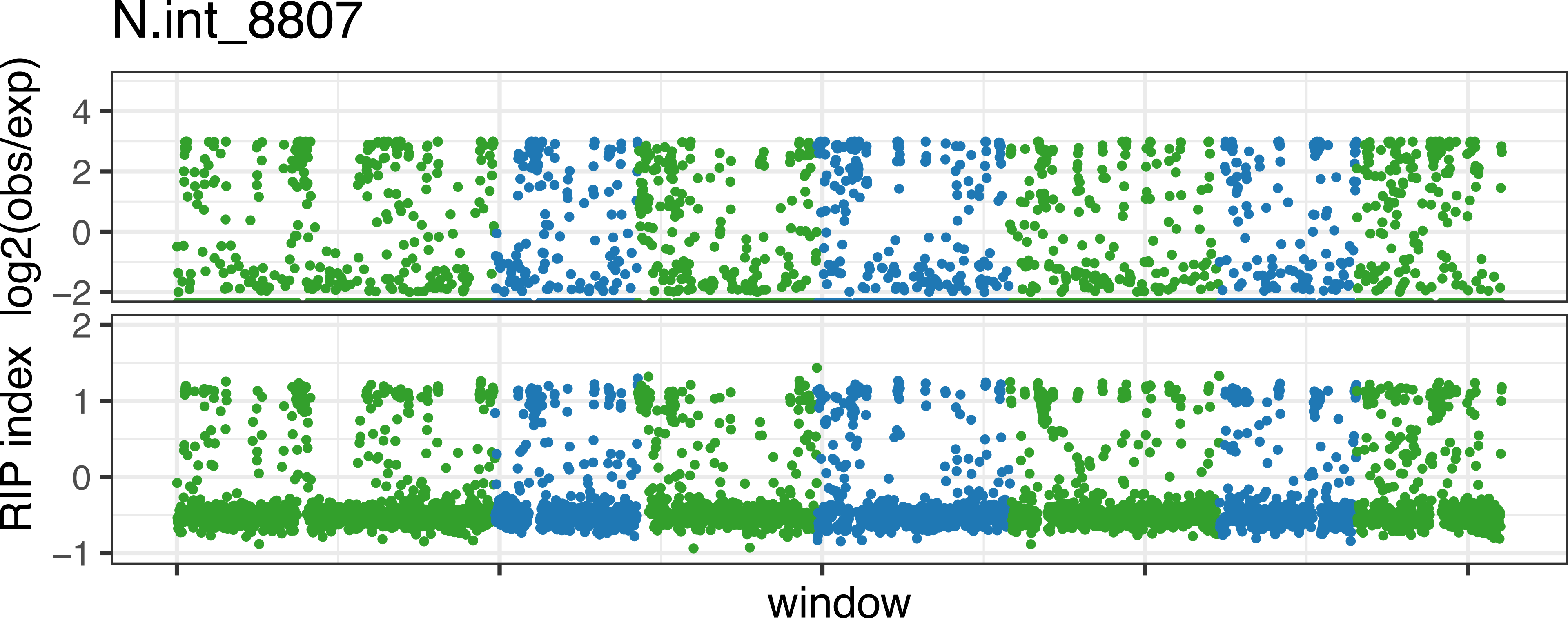

N.int_8807
log2(obs/exp)
4
2
0
−2
RIP index
2
1
0
−1
window

N.int_8761

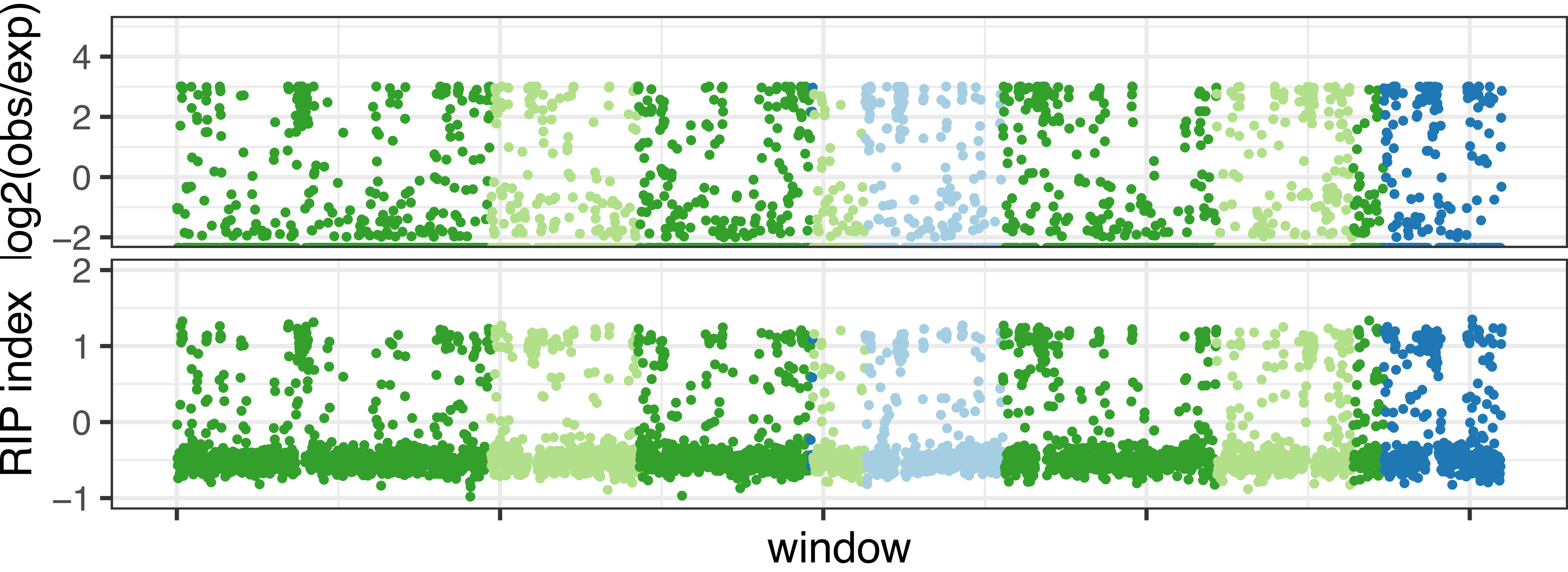

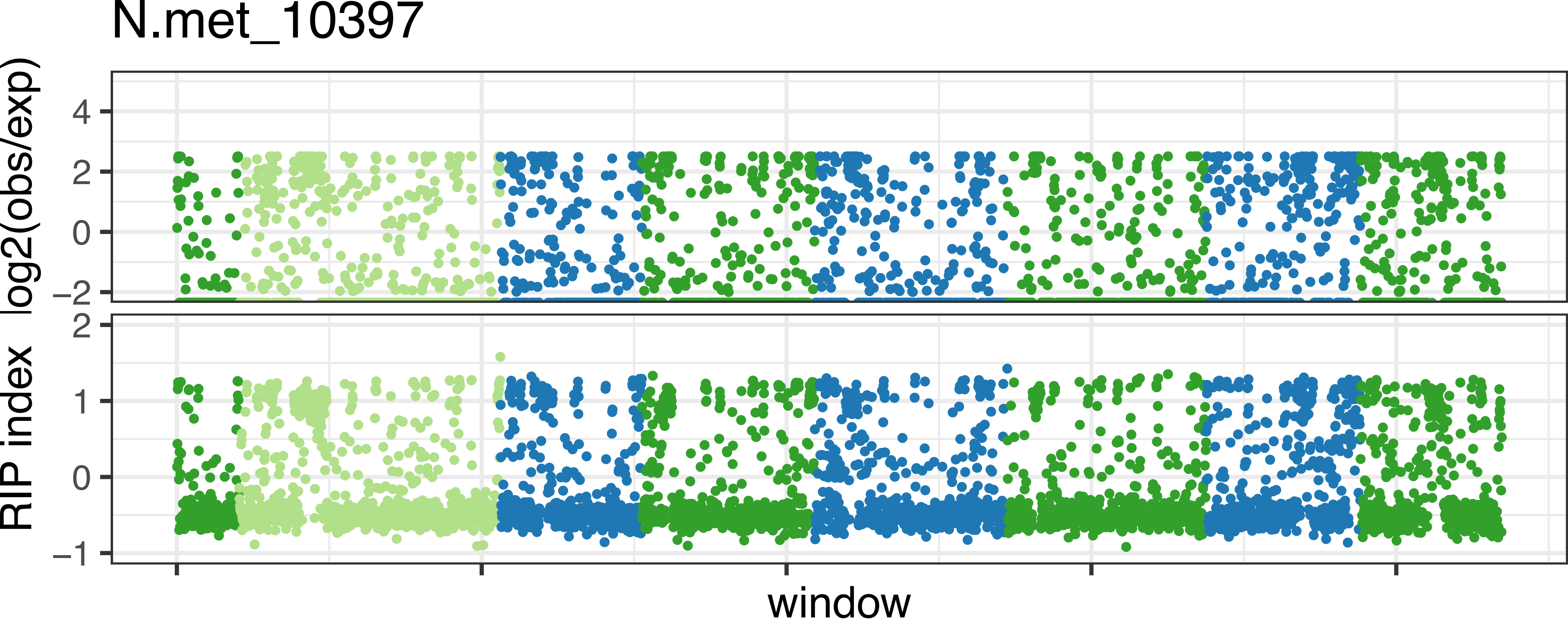
N.met_10397
log2(obs/exp)
4
2
0
−2
RIP index
2
1
0
−1
window

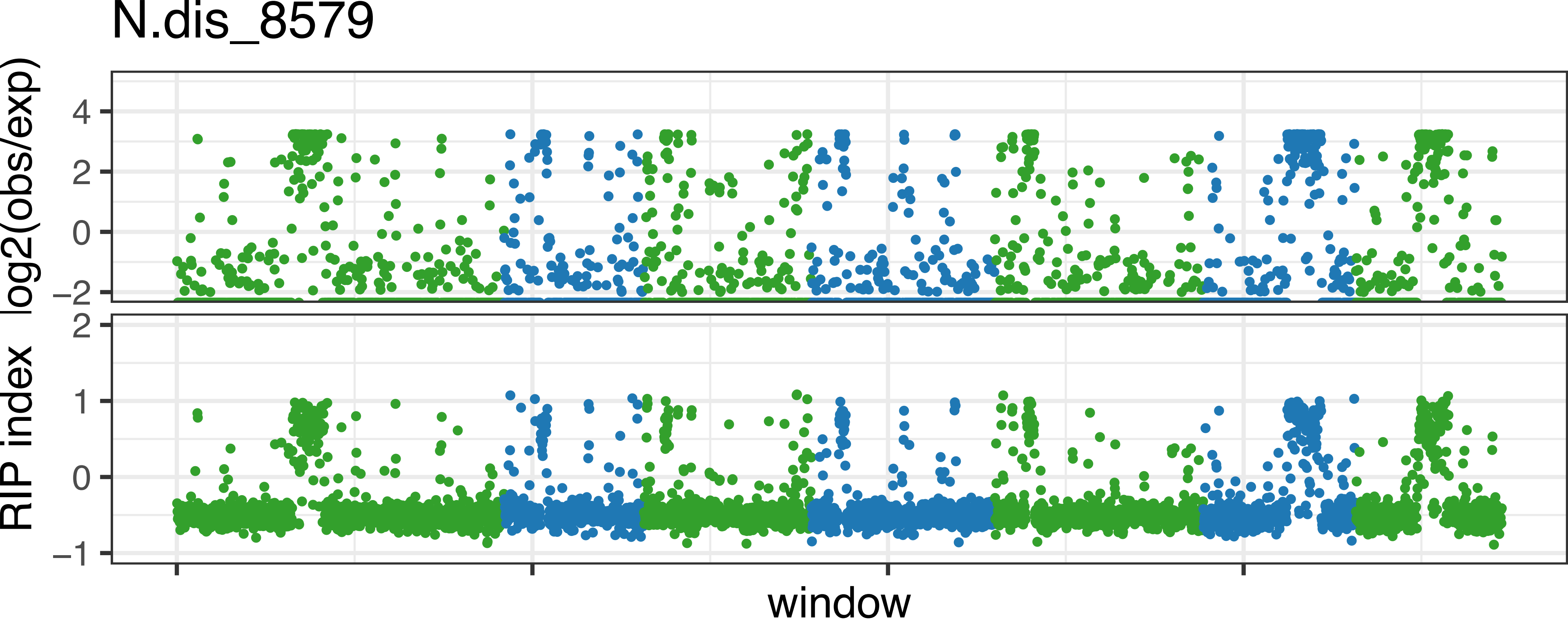

N.dis_8579
log2(obs/exp)
4
2
0
−2
RIP index
2
1
0
−1
window
