## Supplementary material for "Transposon- and genome dynamics in the fungal genus *Neurospora*: insights from nearly gapless genome assemblies": Figure S4

N.tet_L6_2508

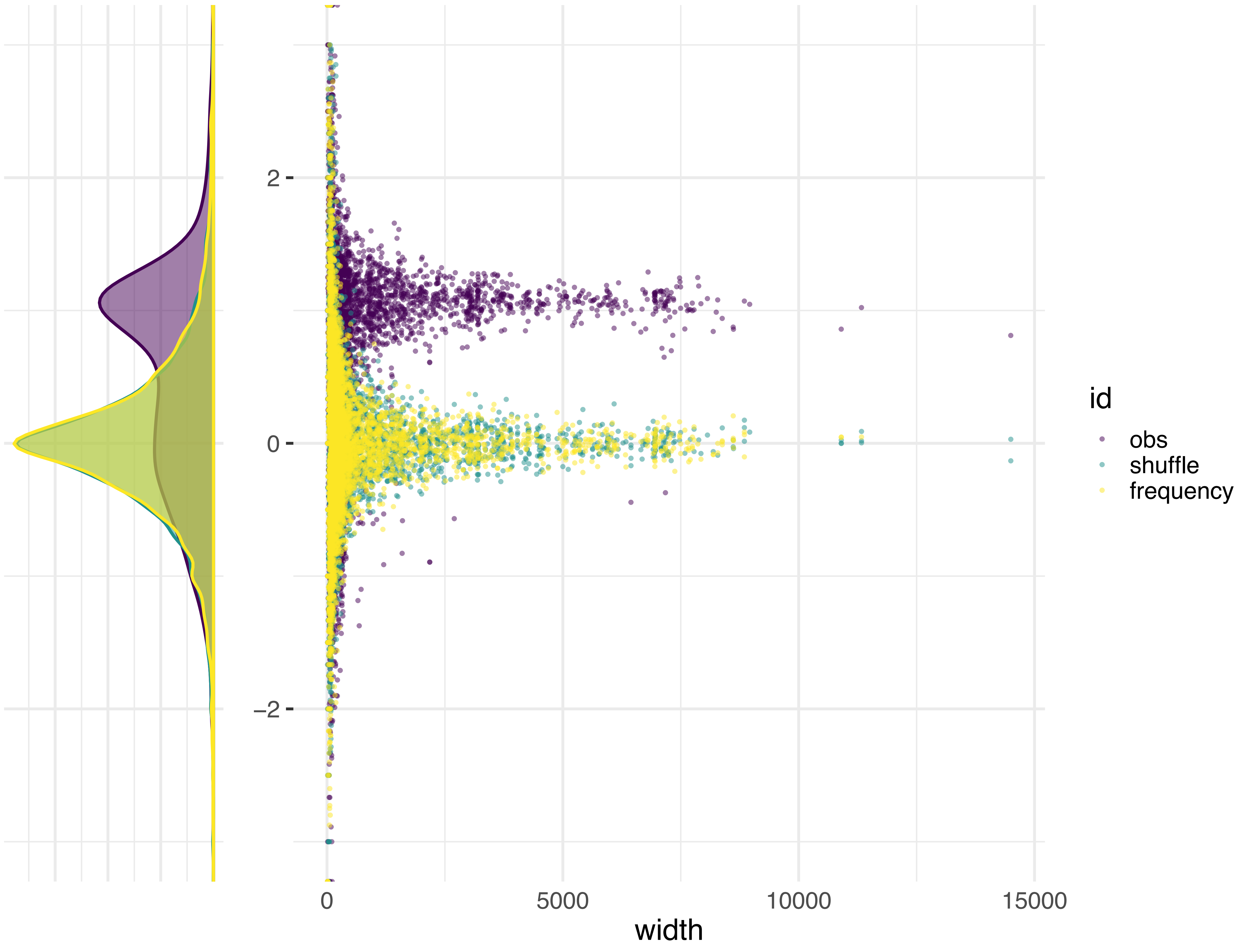

N.tet_L6_2509

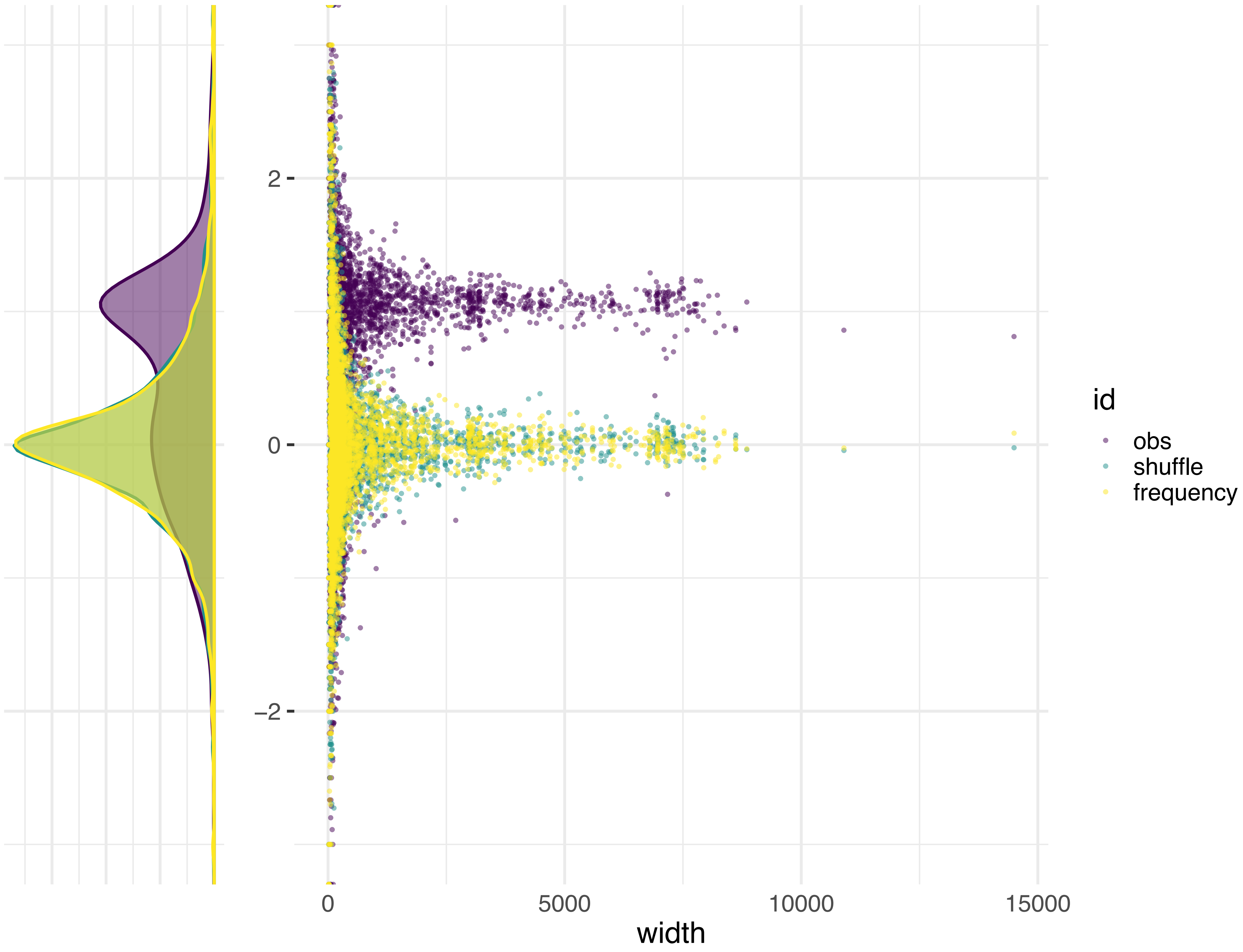

N.tet_L9_10752

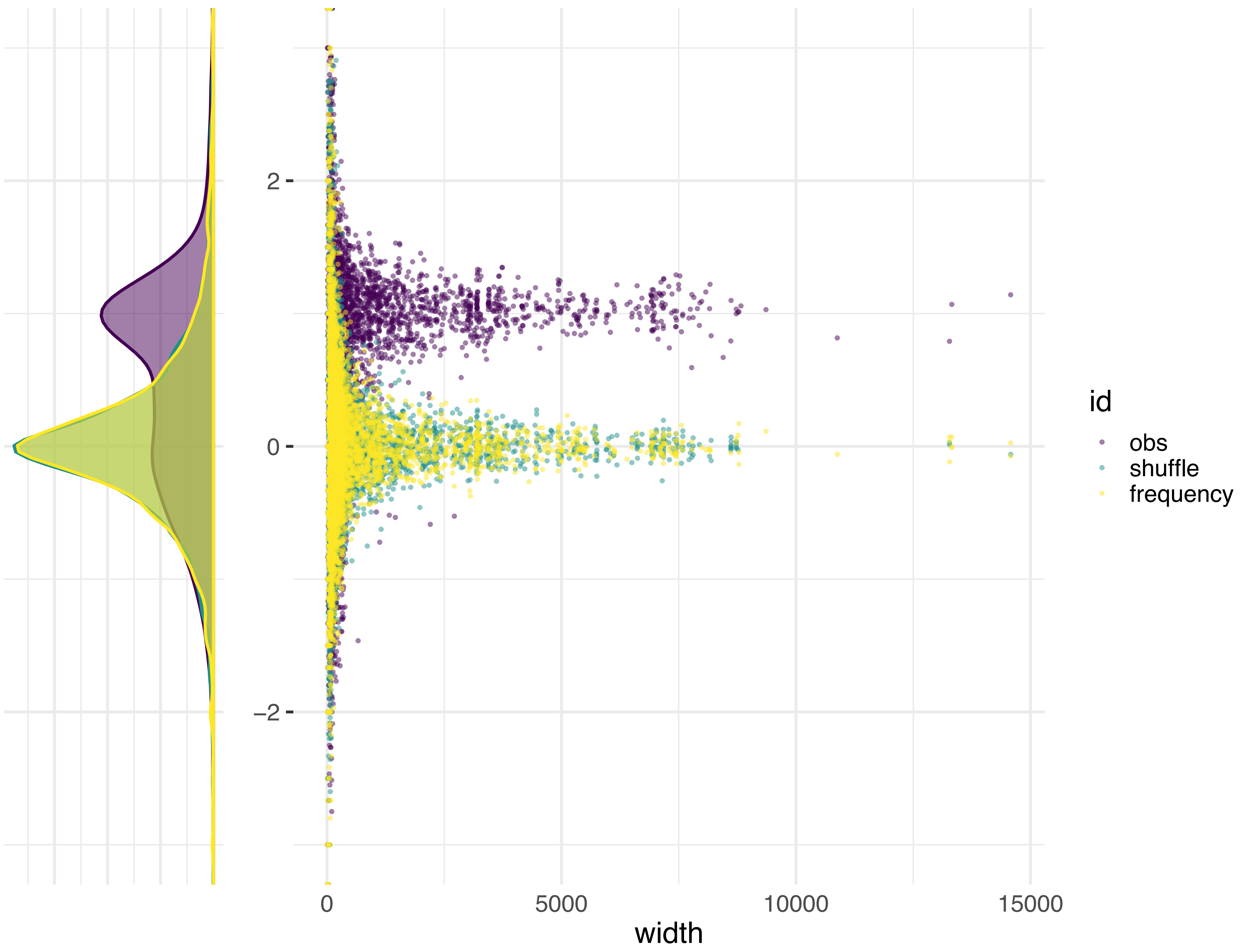

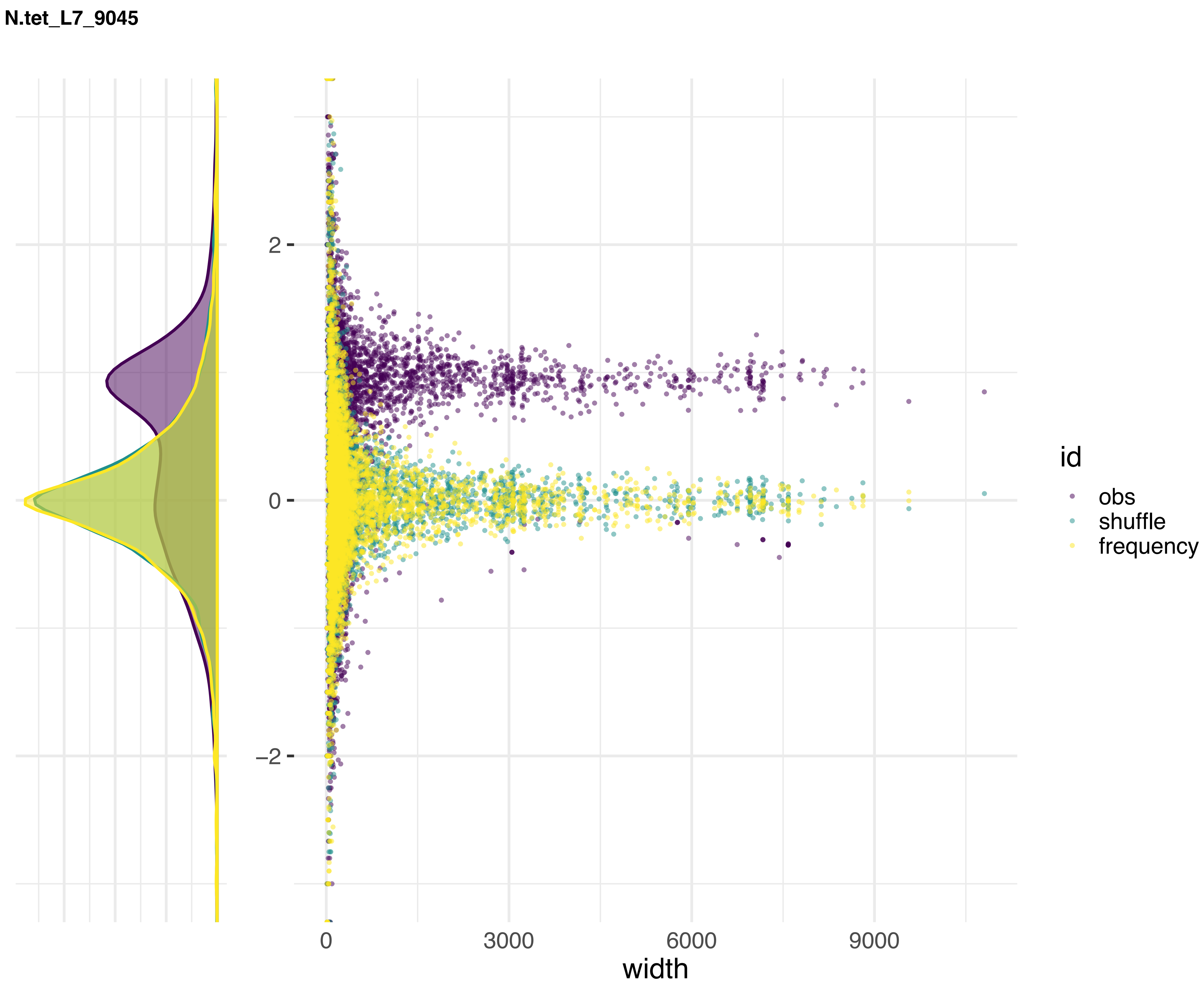
N.tet_L7_9045
2
0
−2
0
3000
6000
9000
width
id
obs
shuffle
frequency

N.tet_L7_9046

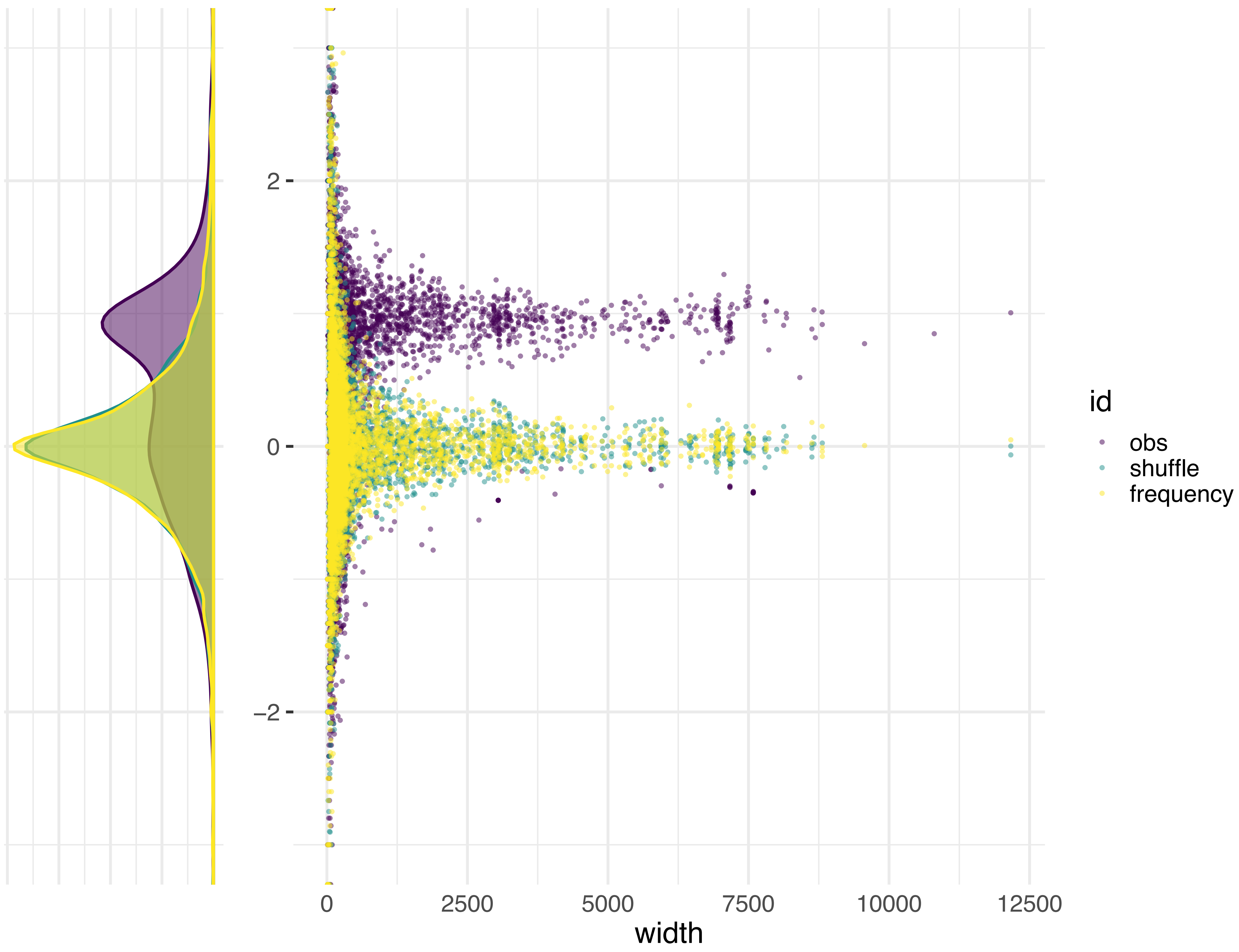
2
0
−2
0
2500
5000
7500
10000
12500
width
id
obs
shuffle
frequency

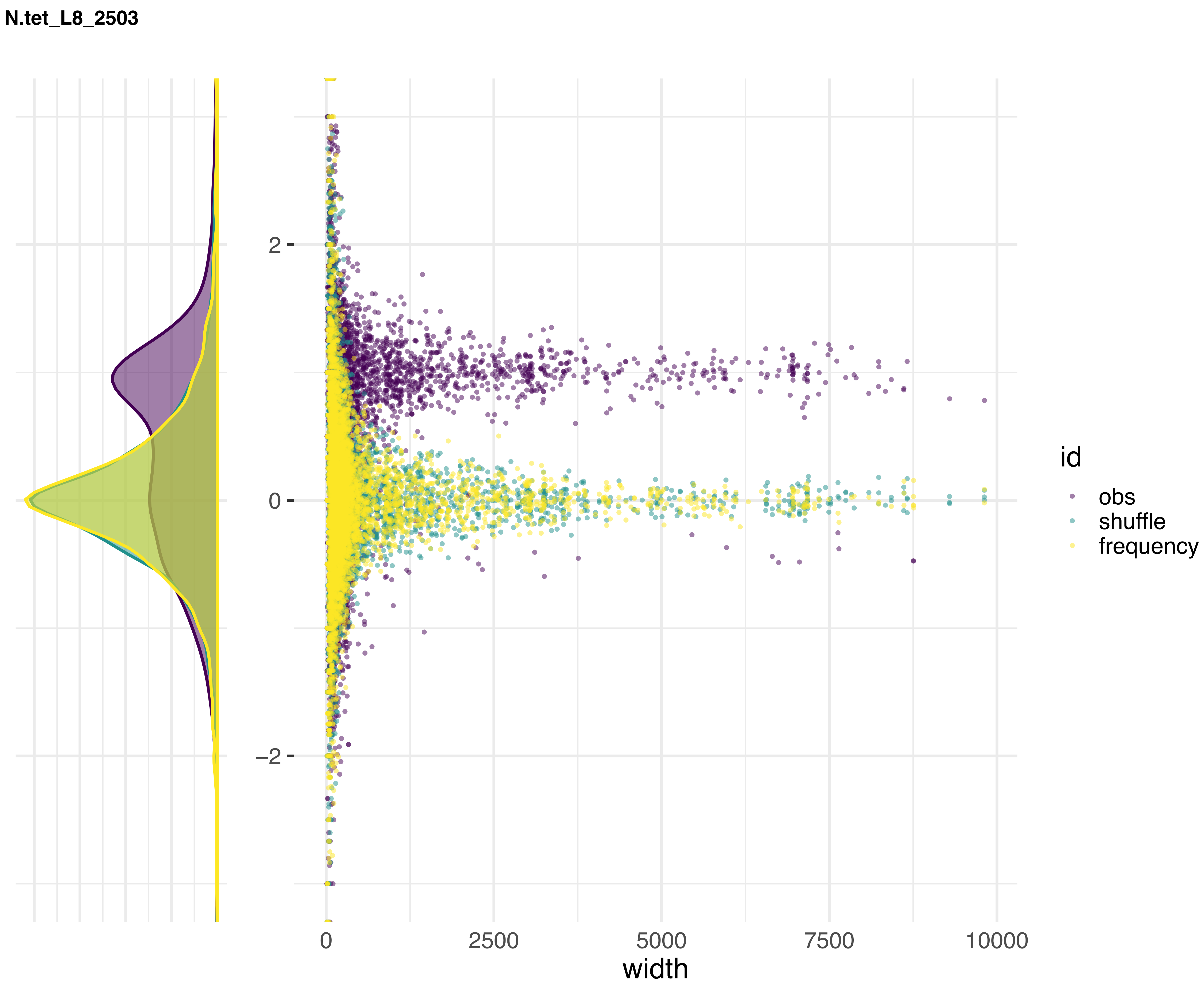
N.tet_L8_2503
2
0
−2
0
2500
5000
7500
10000
width
id
obs
shuffle
frequency

**N.tet_L8_2504**

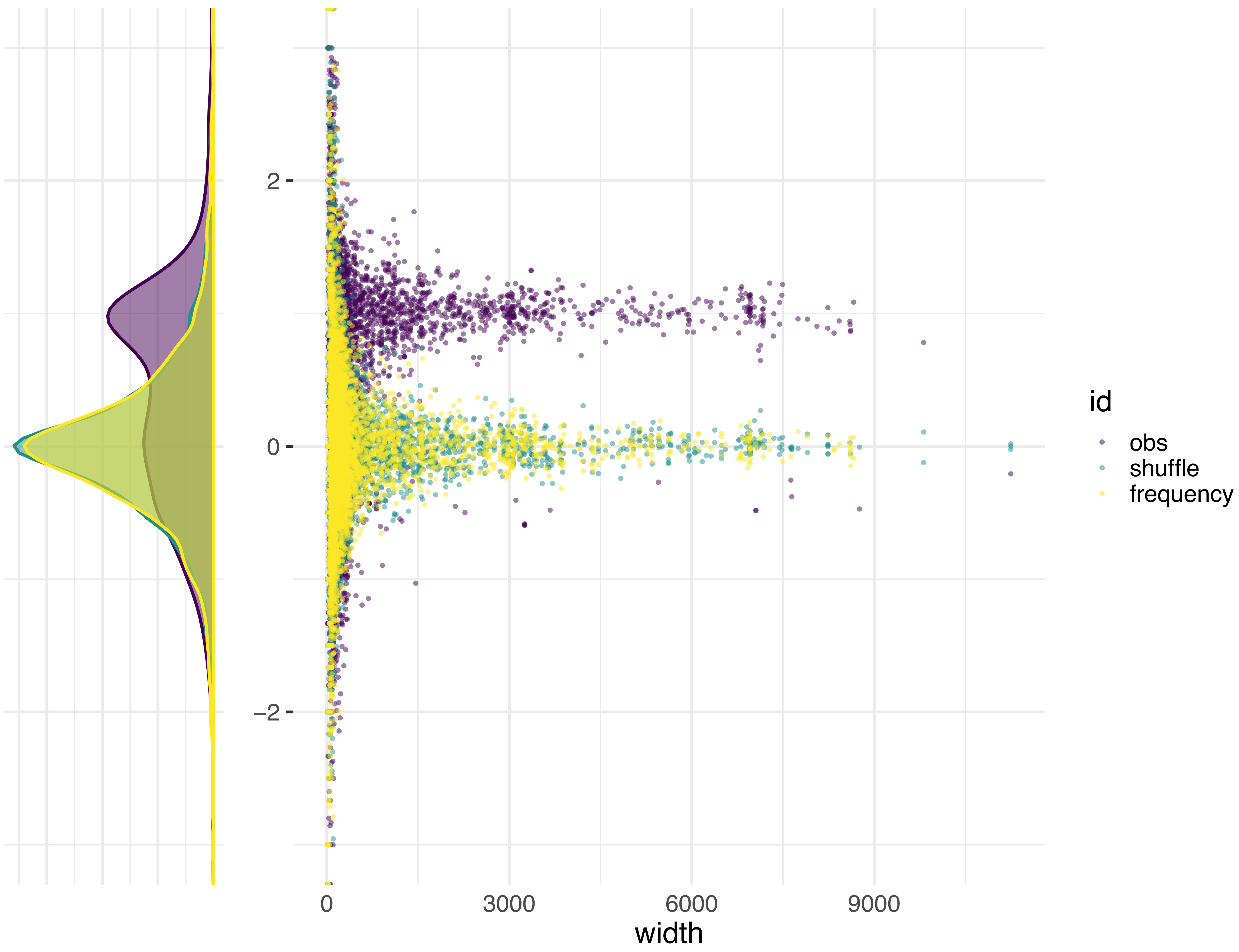

N.sit_5940

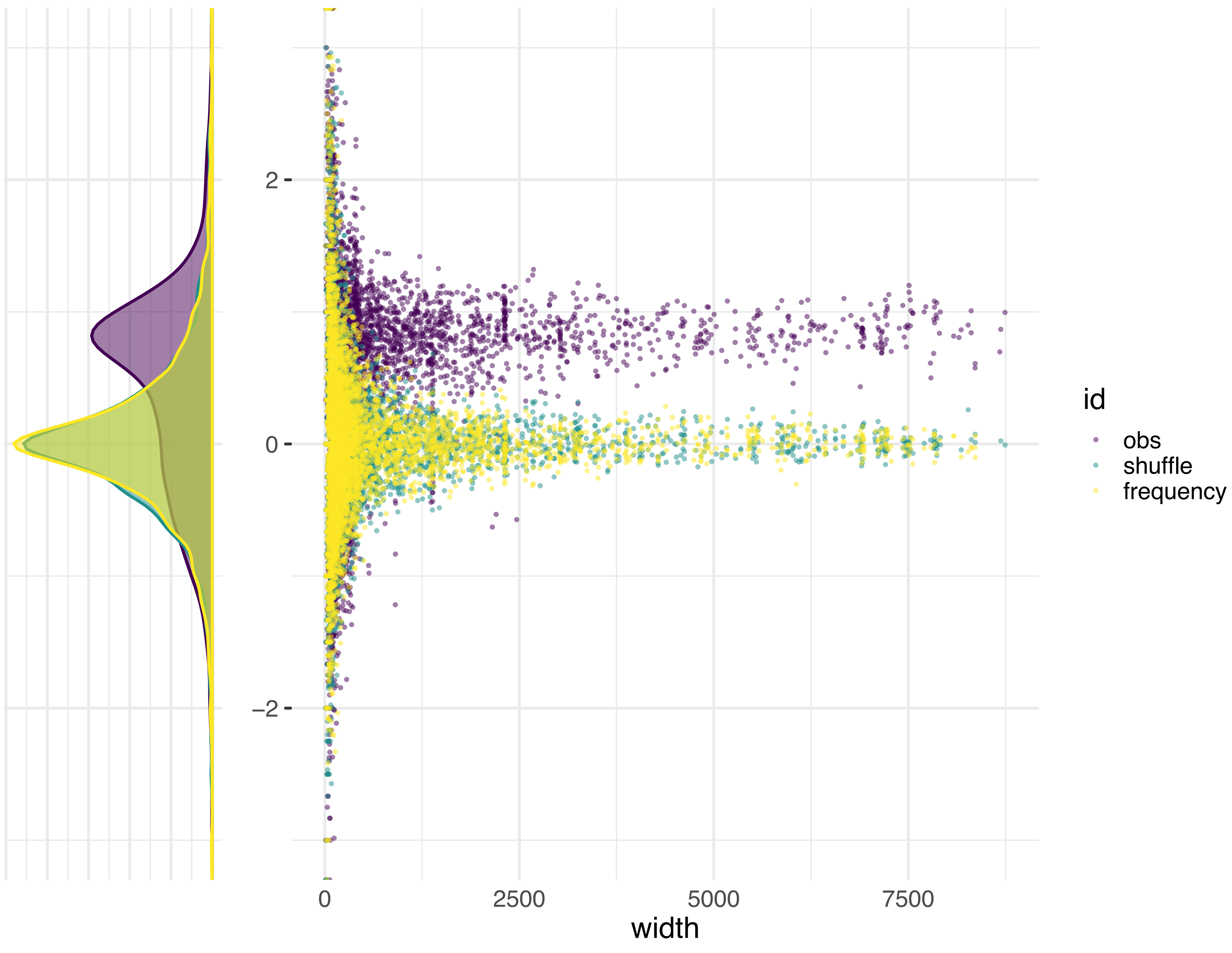

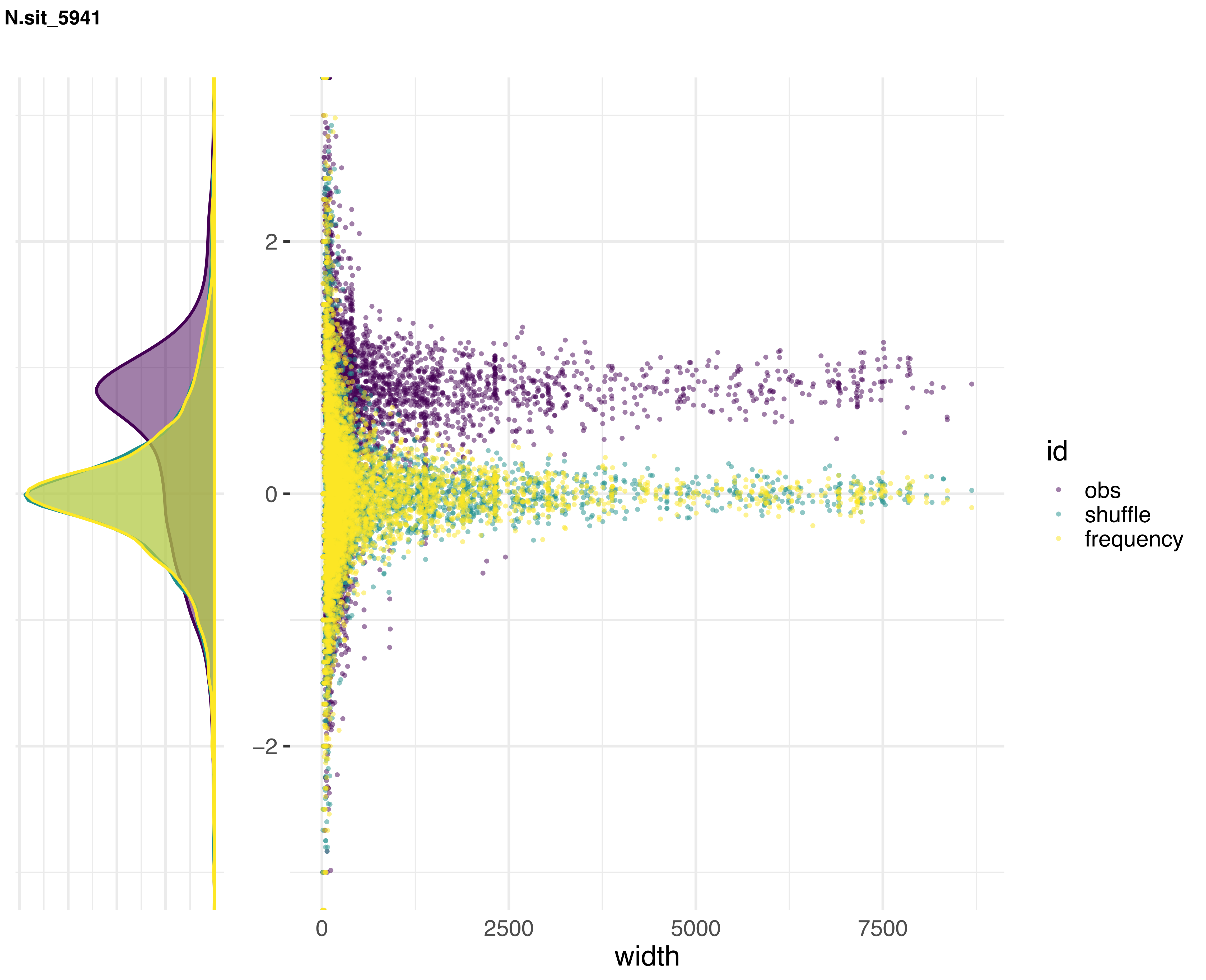
N.sit_5941
2
0
−2
0
2500
5000
7500
width
id
obs
shuffle
frequency

**N.sit_W1426**

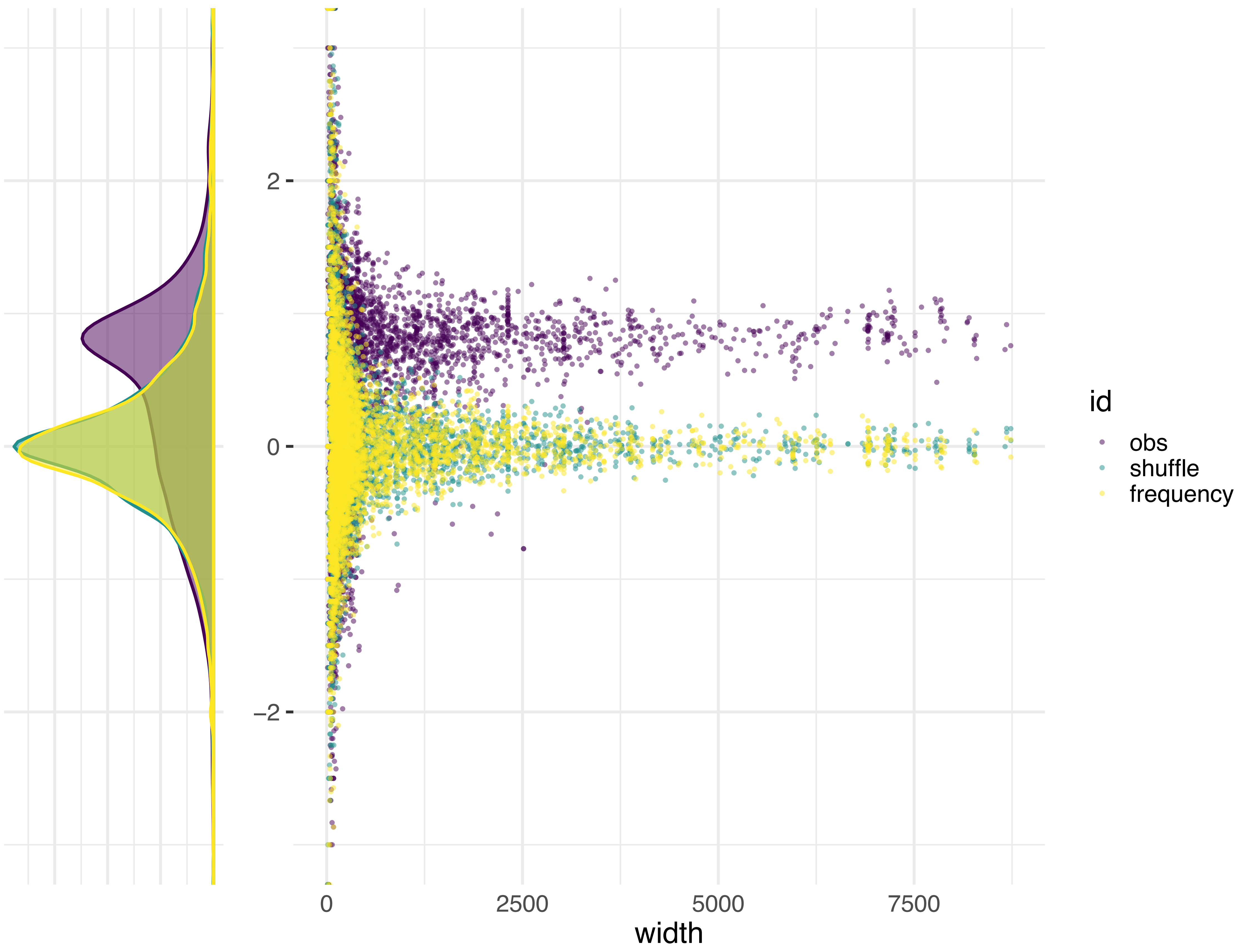

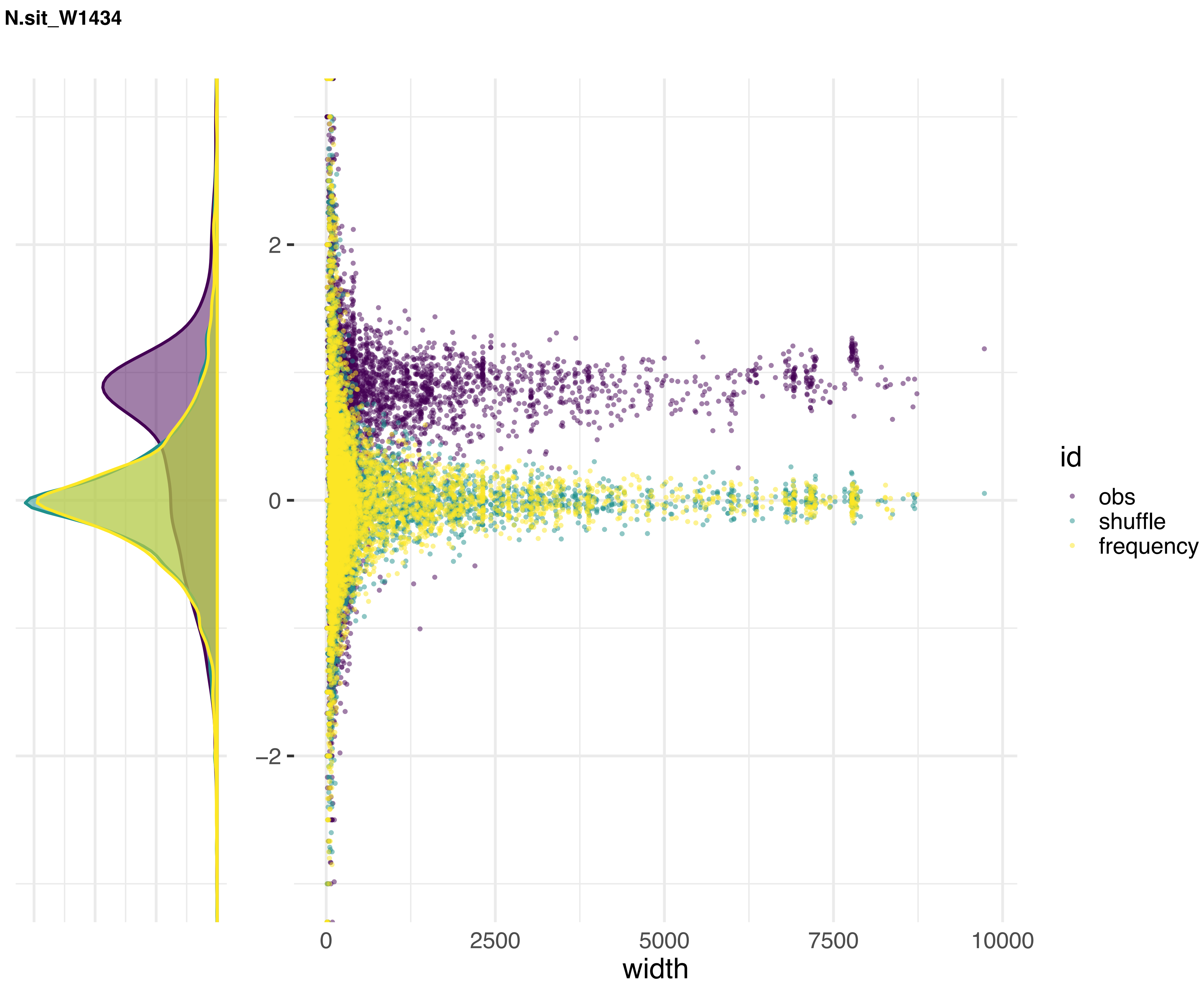
N.sit_W1434
2
0
−2
0
2500
5000
7500
10000
width
id
obs
shuffle
frequency

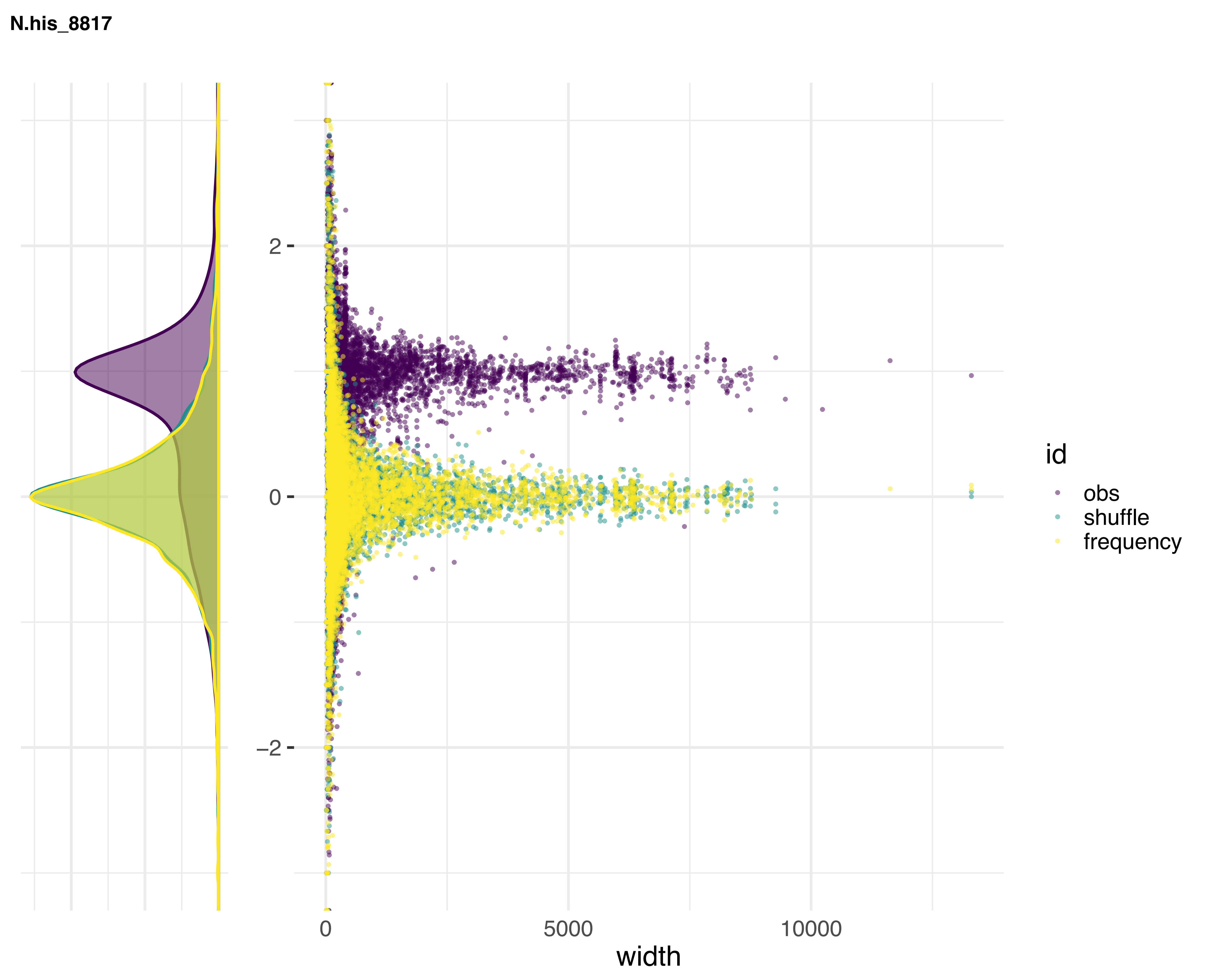

N.his_8817
2
0
−2
0
5000
10000
width
id
obs
shuffle
frequency

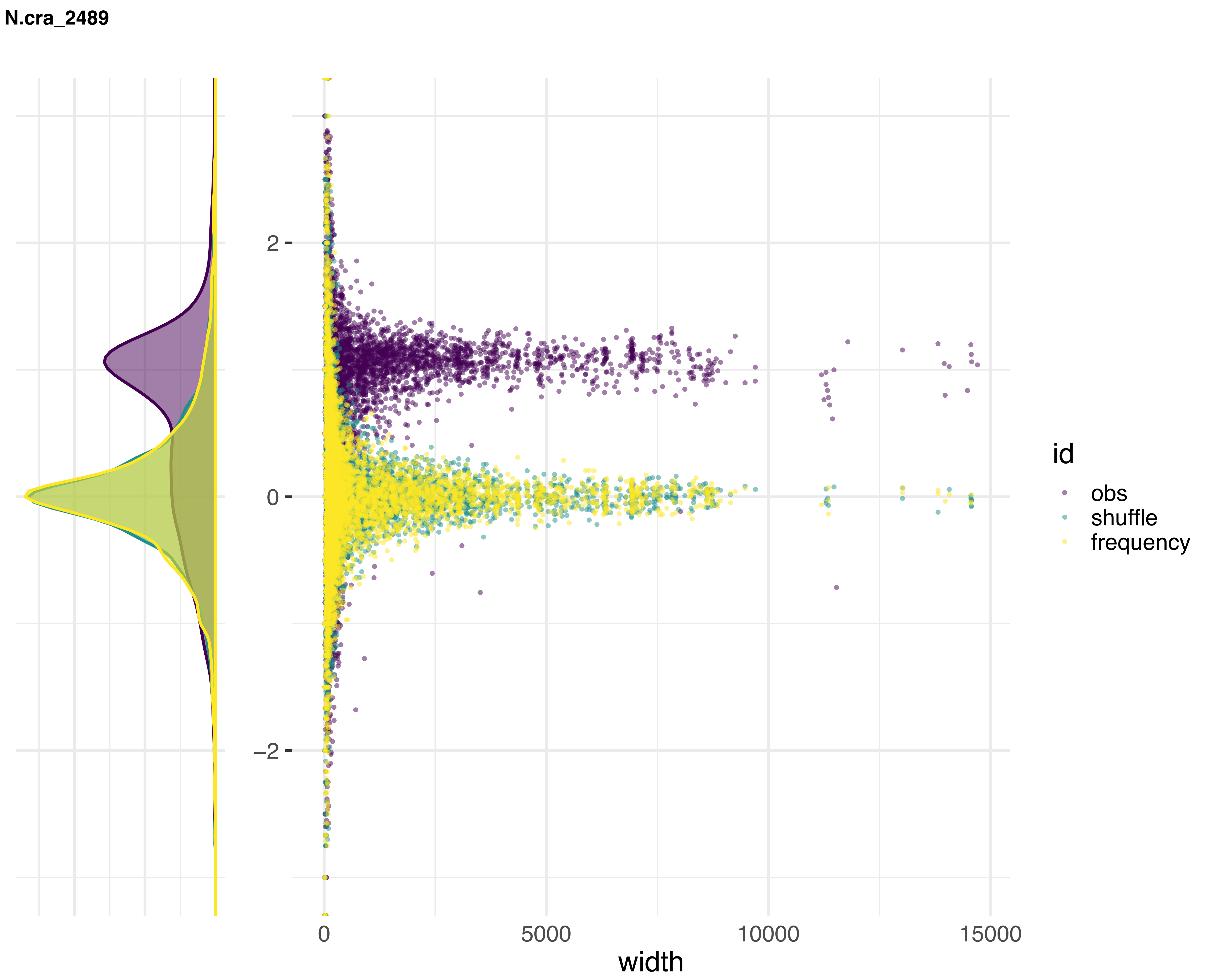

N.cra_2489
2
0
−2
0
5000
10000
15000
width
id
obs
shuffle
frequency

**N.int_8767**

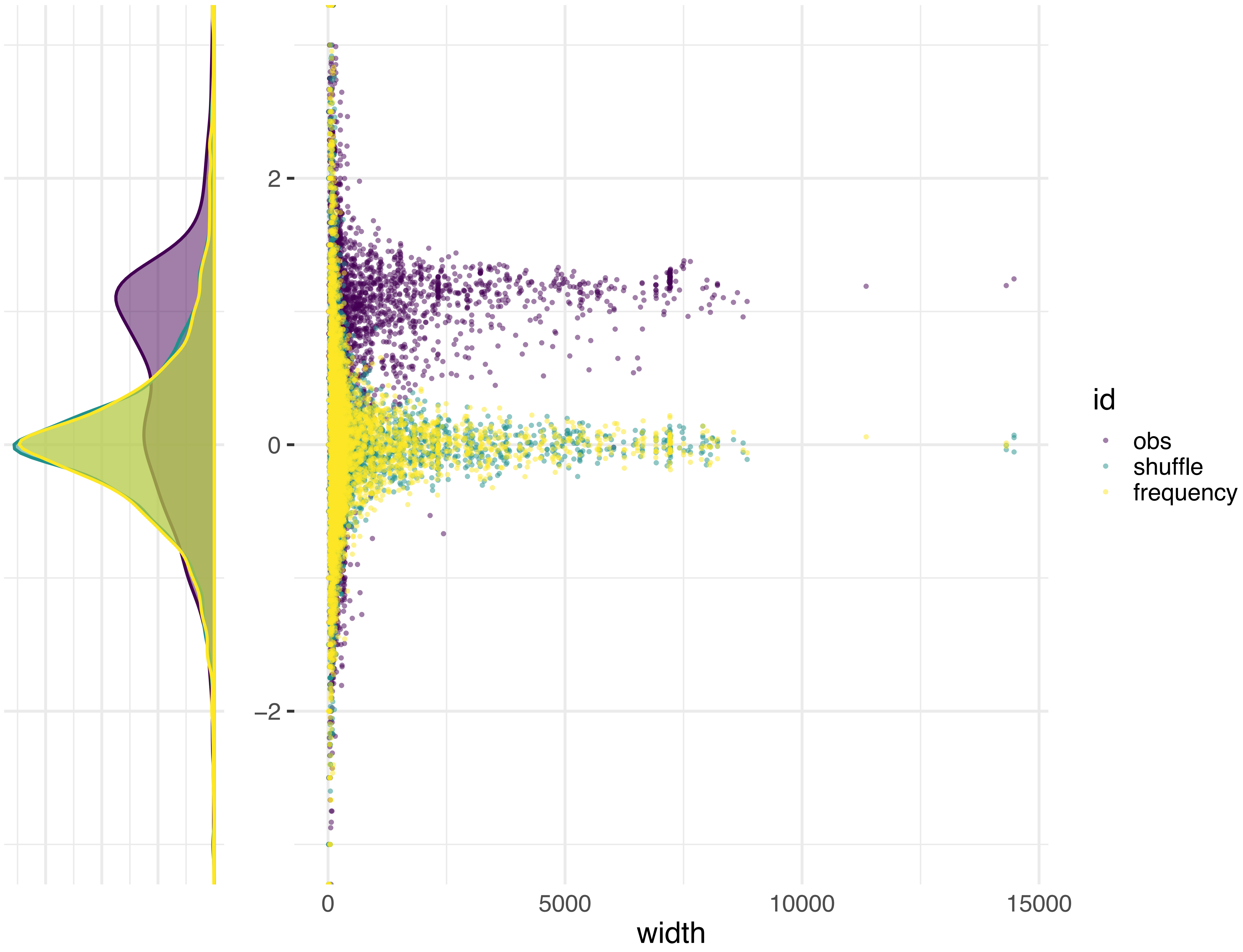

N.int_8807

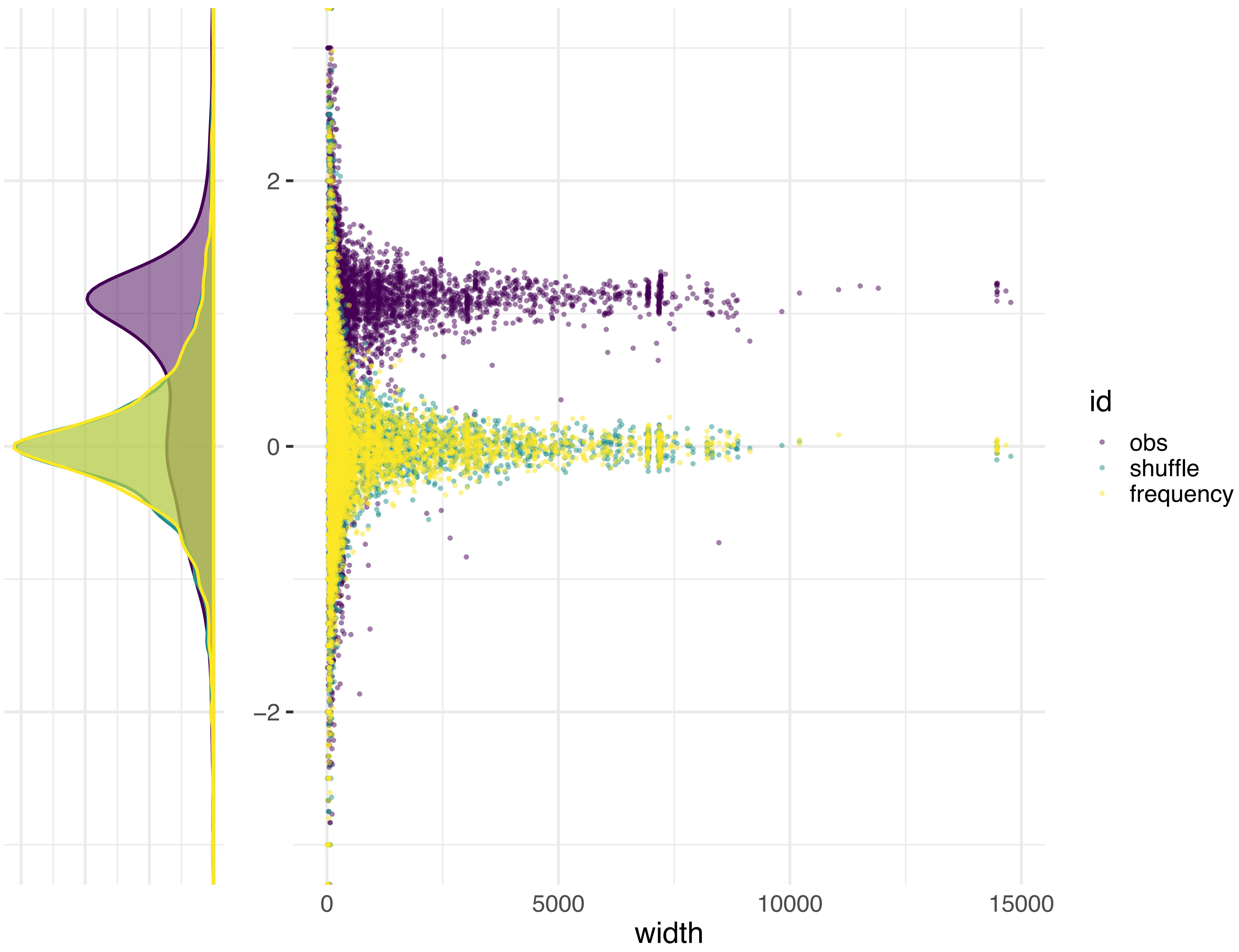

N.int_8761

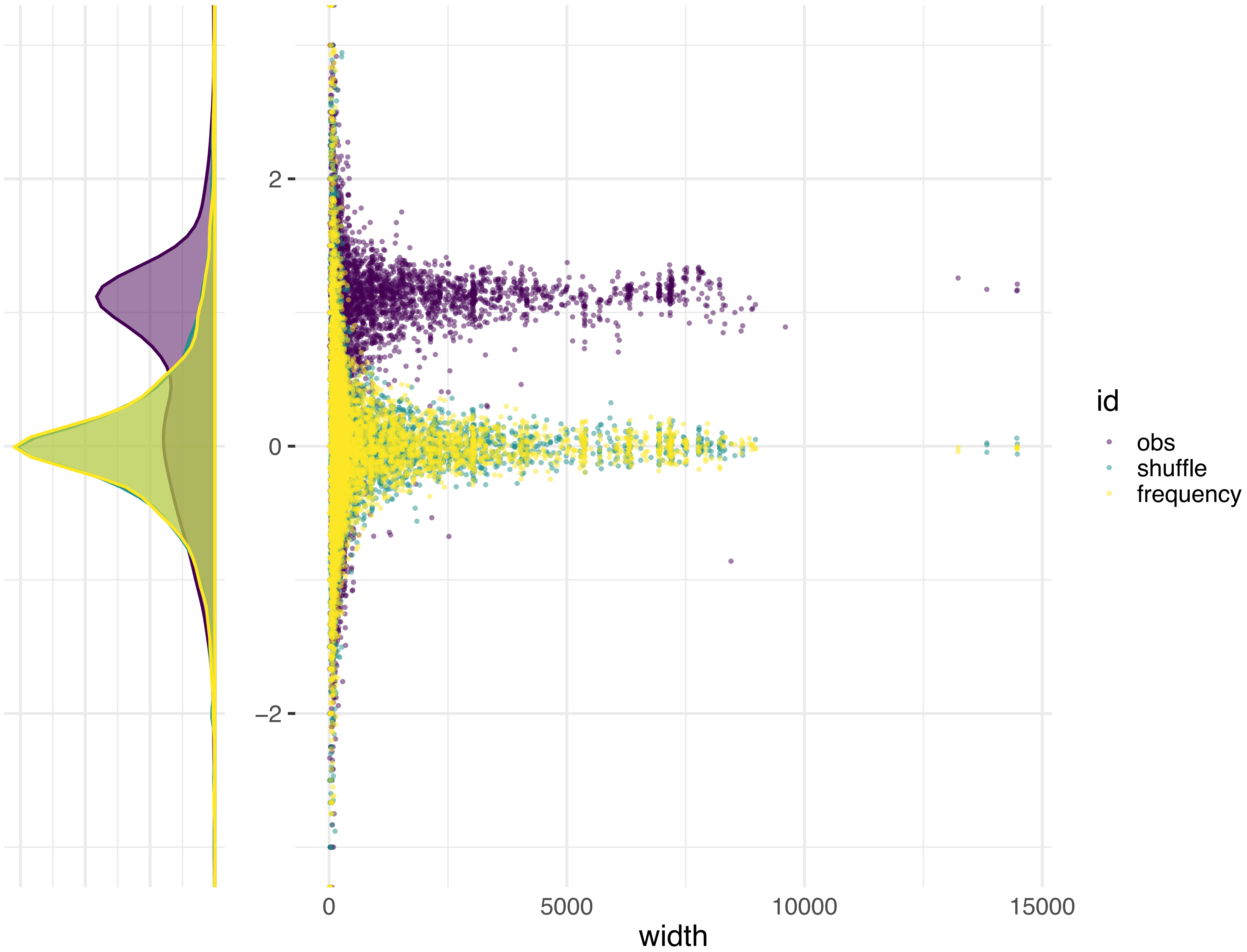

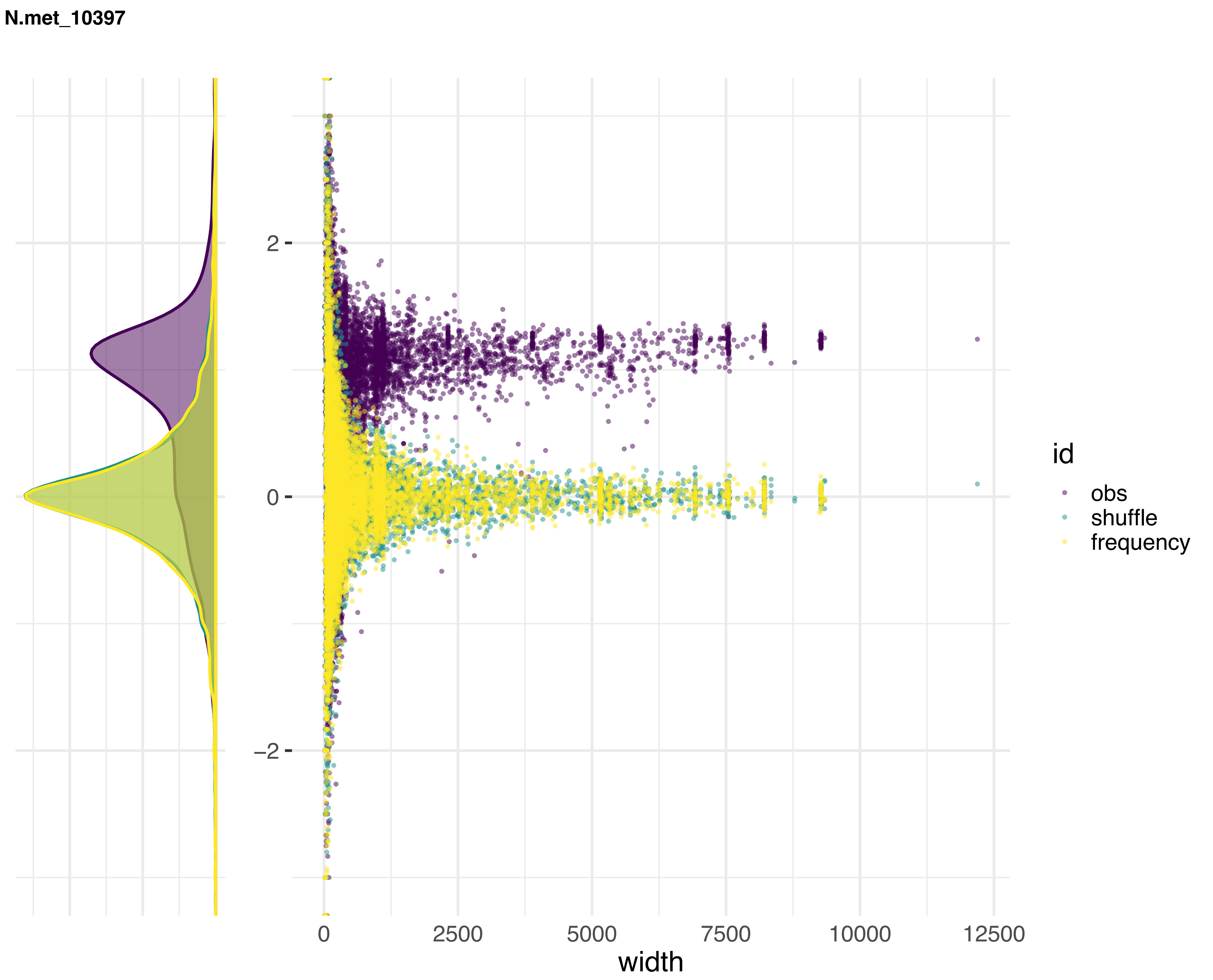
N.met_10397
2
0
−2
0
2500
5000
7500
10000
12500
width
id
obs
shuffle
frequency

**N.dis_8579**

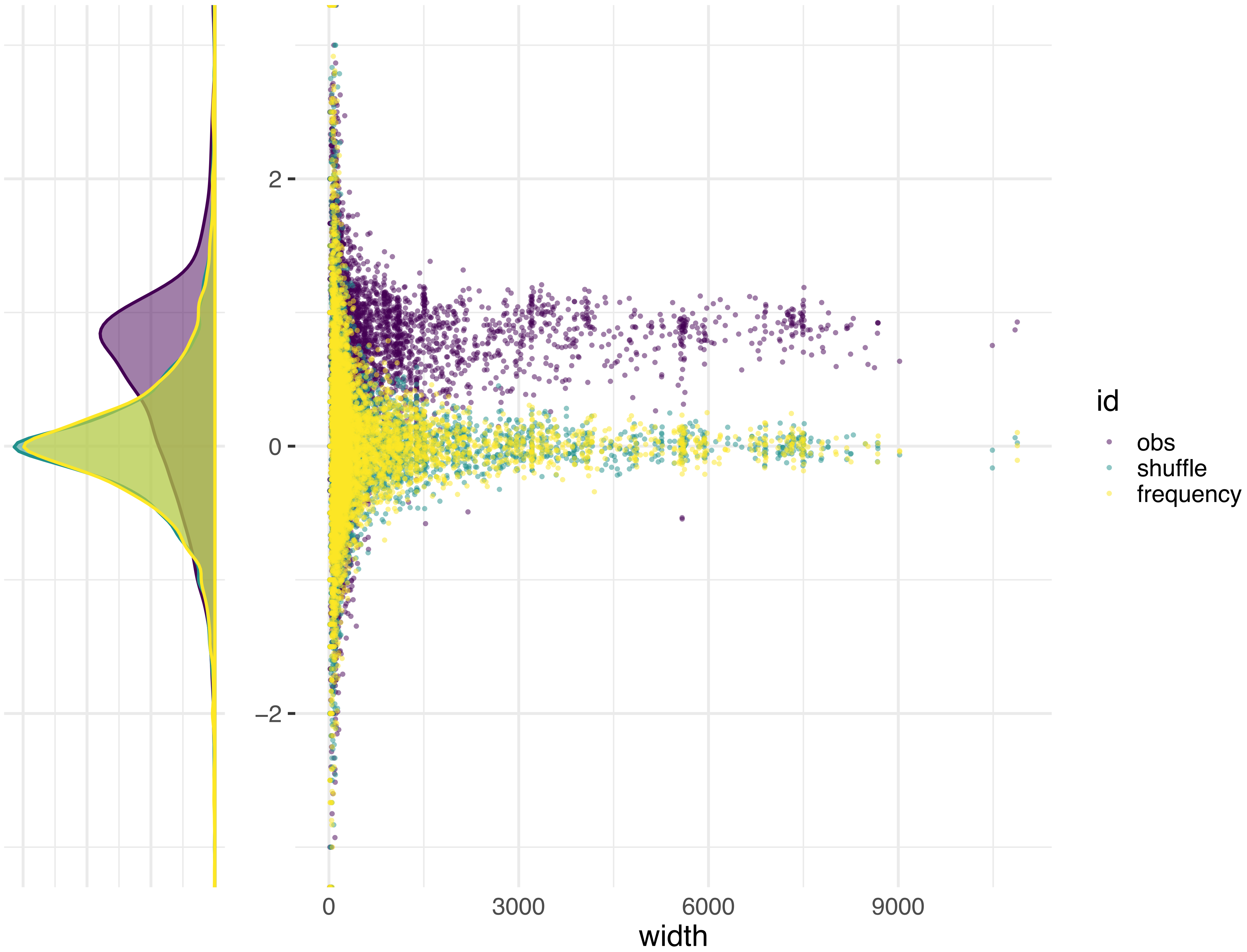
