## Supplementary material for "Transposon- and genome dynamics in the fungal genus *Neurospora*: insights from nearly gapless genome assemblies": Figure S5

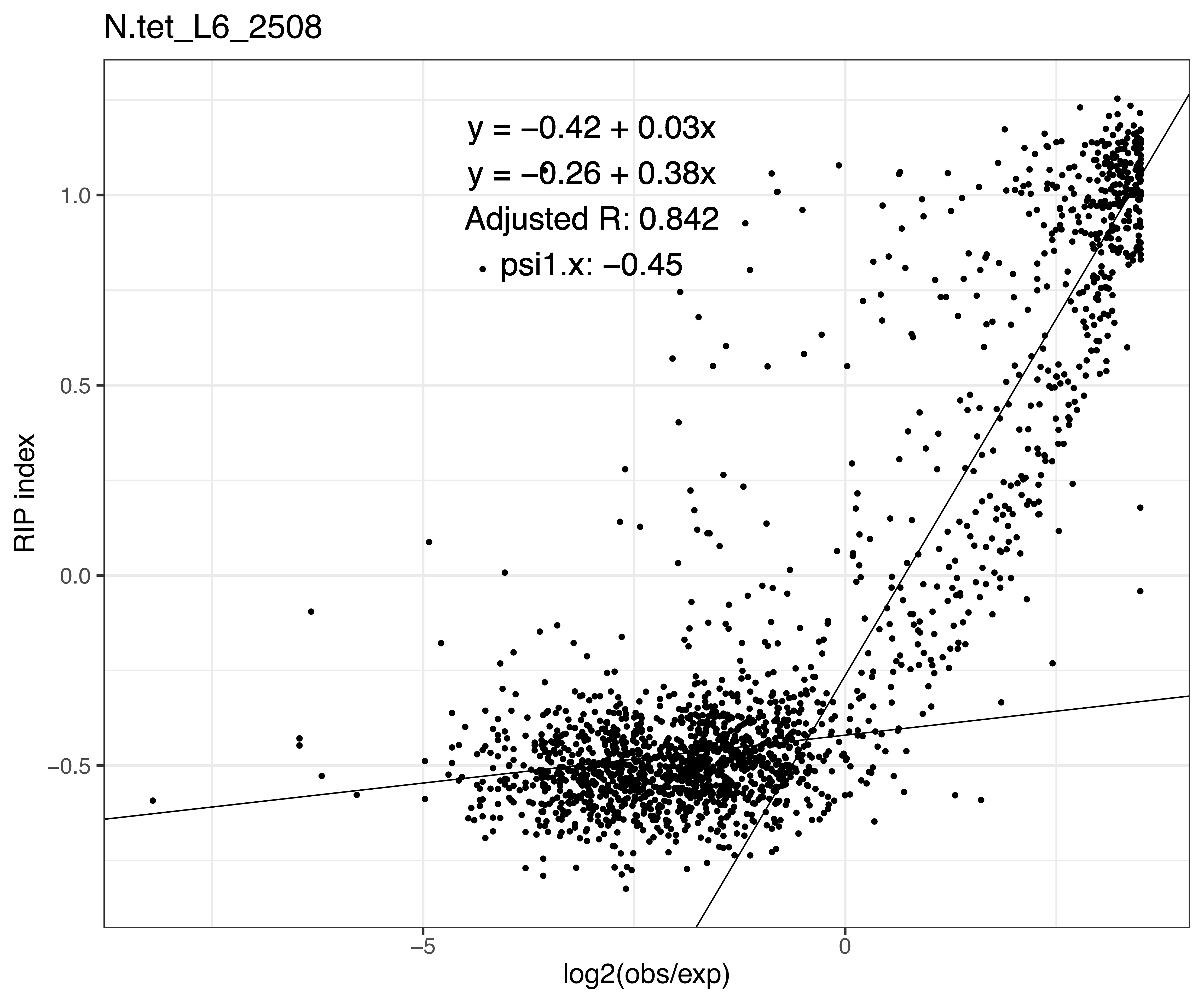

N.tet_L6_2508
y = −0.42 + 0.03x
y = −0.26 + 0.38x
Adjusted R: 0.842
psi1.x: −0.45
RIP index
1.0
0.5
0.0
−0.5
−5
0
log2(obs/exp)

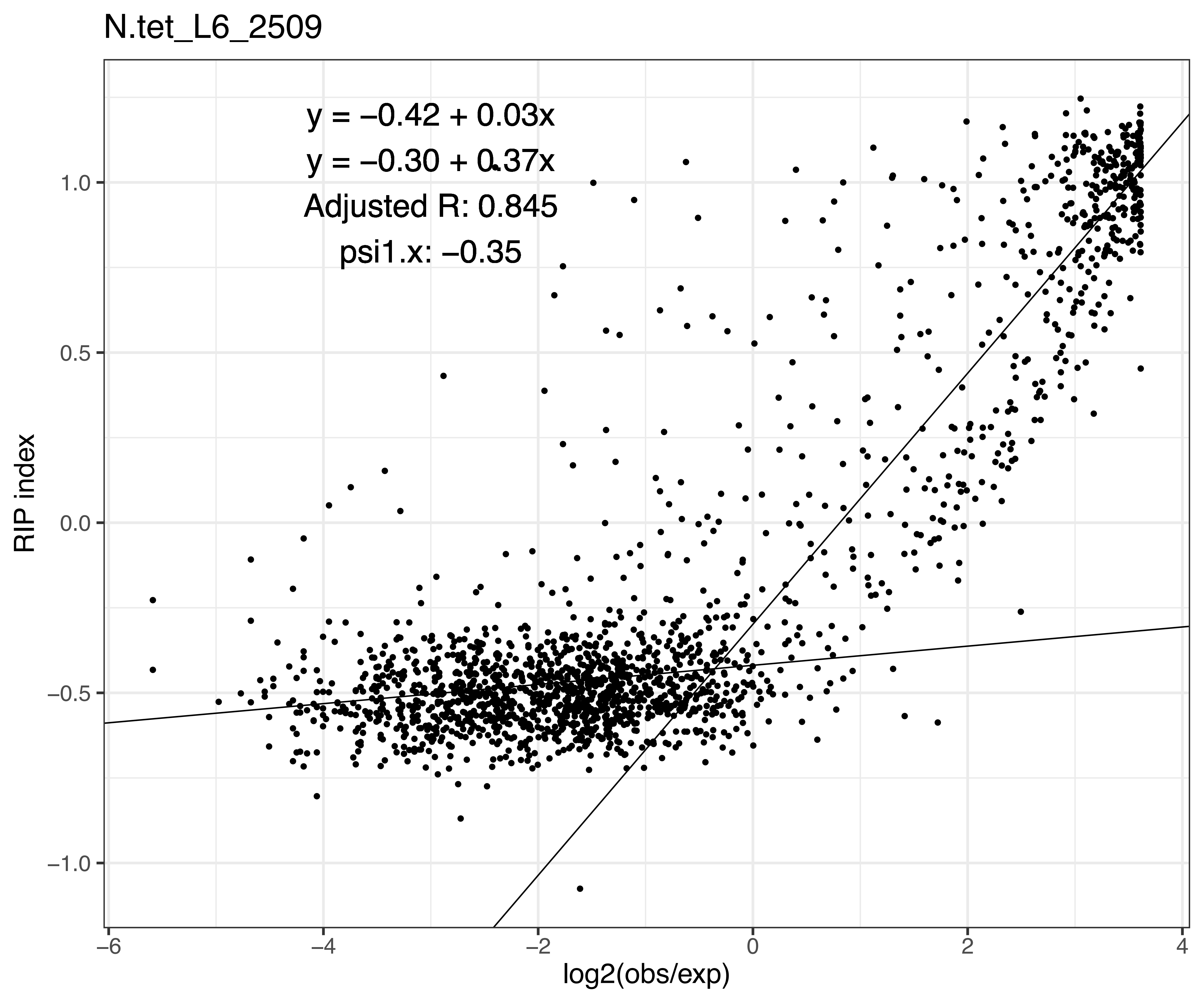

N.tet_L6_2509
y = −0.42 + 0.03x
y = −0.30 + 0.37x
Adjusted R: 0.845
psi1.x: −0.35
RIP index
log2(obs/exp)
1.0
0.5
0.0
−0.5
−1.0
−6
−4
−2
0
2
4

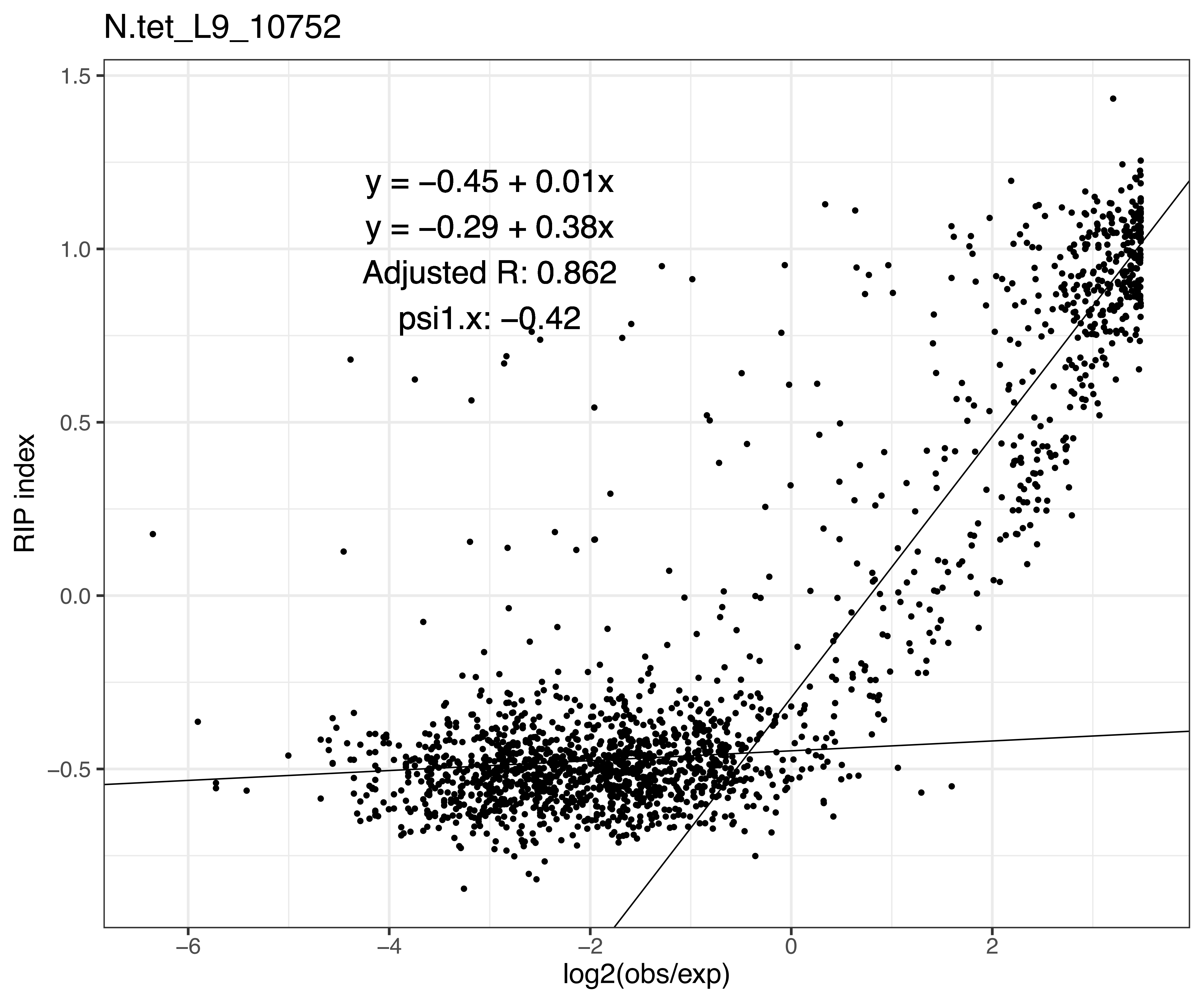

N.tet_L9_10752
y = −0.45 + 0.01x
y = −0.29 + 0.38x
Adjusted R: 0.862
psi1.x: −0.42
RIP index
log2(obs/exp)
1.5
1.0
0.5
0.0
−0.5
−6
−4
−2
0
2

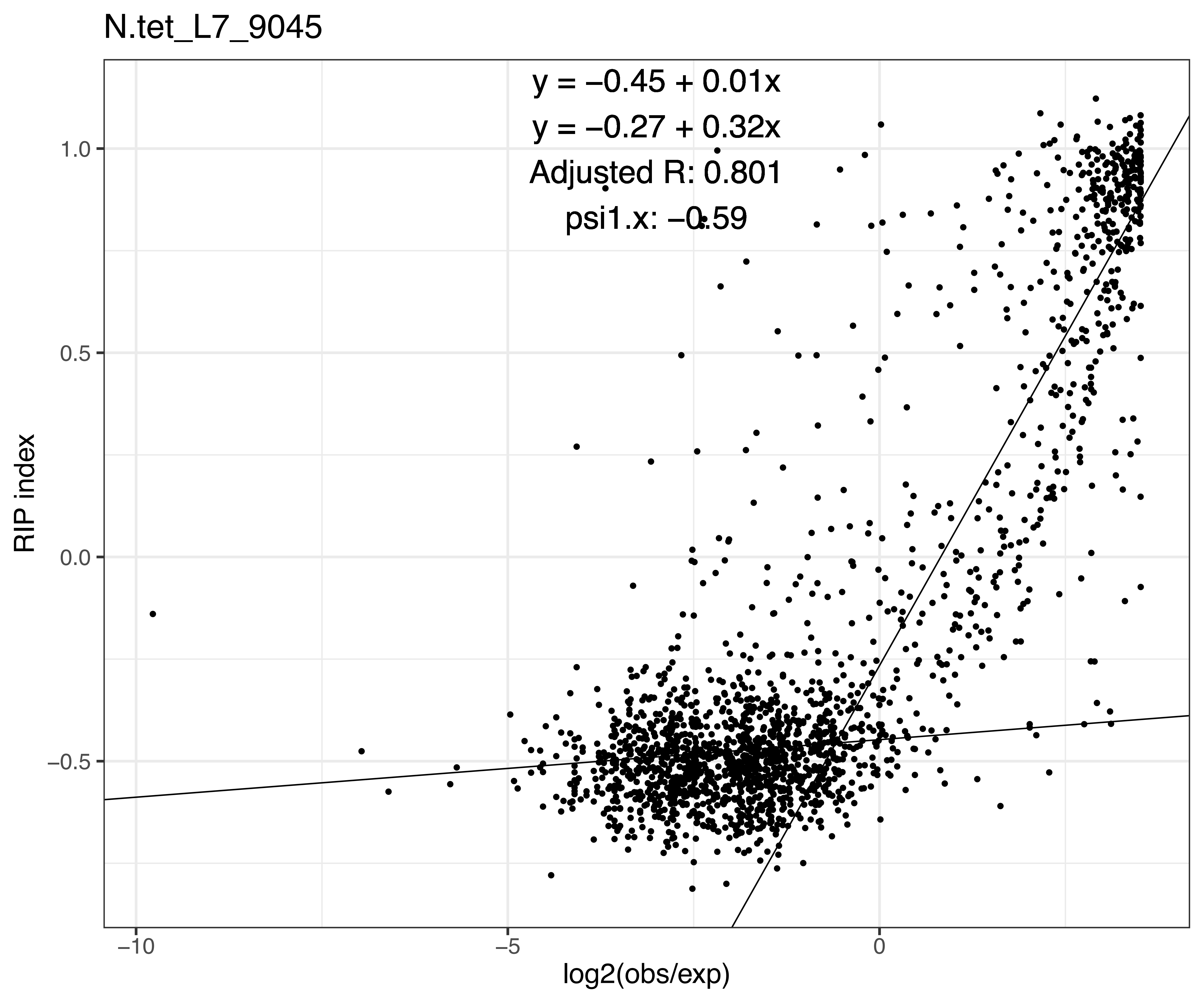
N.tet_L7_9045
y = −0.45 + 0.01x
y = −0.27 + 0.32x
Adjusted R: 0.801
psi1.x: −0.59
RIP index
1.0
0.5
0.0
−0.5
−10
−5
0
log2(obs/exp)

N.tet_L7_9046

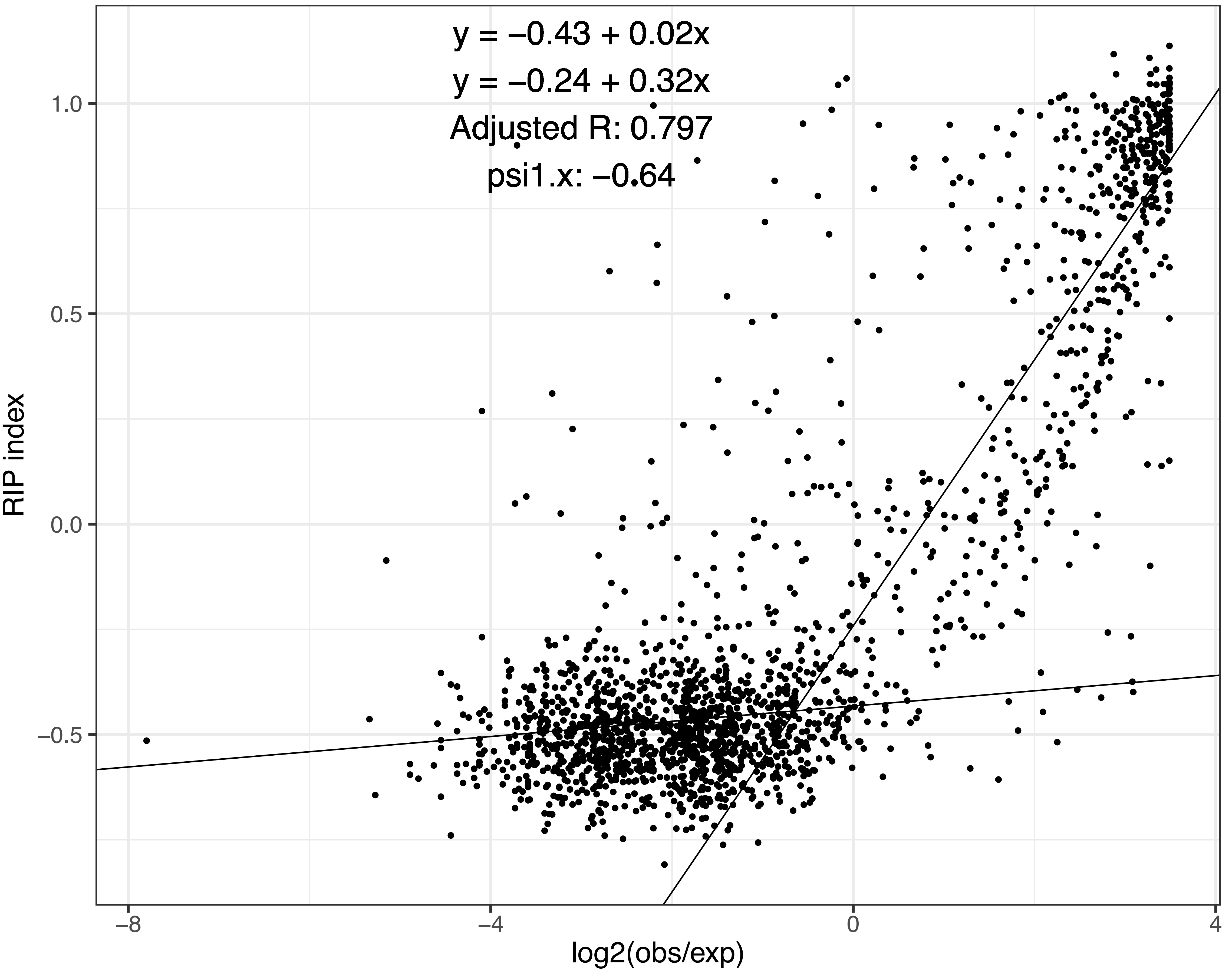

y = −0.43 + 0.02x
y = −0.24 + 0.32x
Adjusted R: 0.797
psi1.x: −0.64
RIP index
log2(obs/exp)
1.0
0.5
0.0
−0.5
−8
−4
0
4

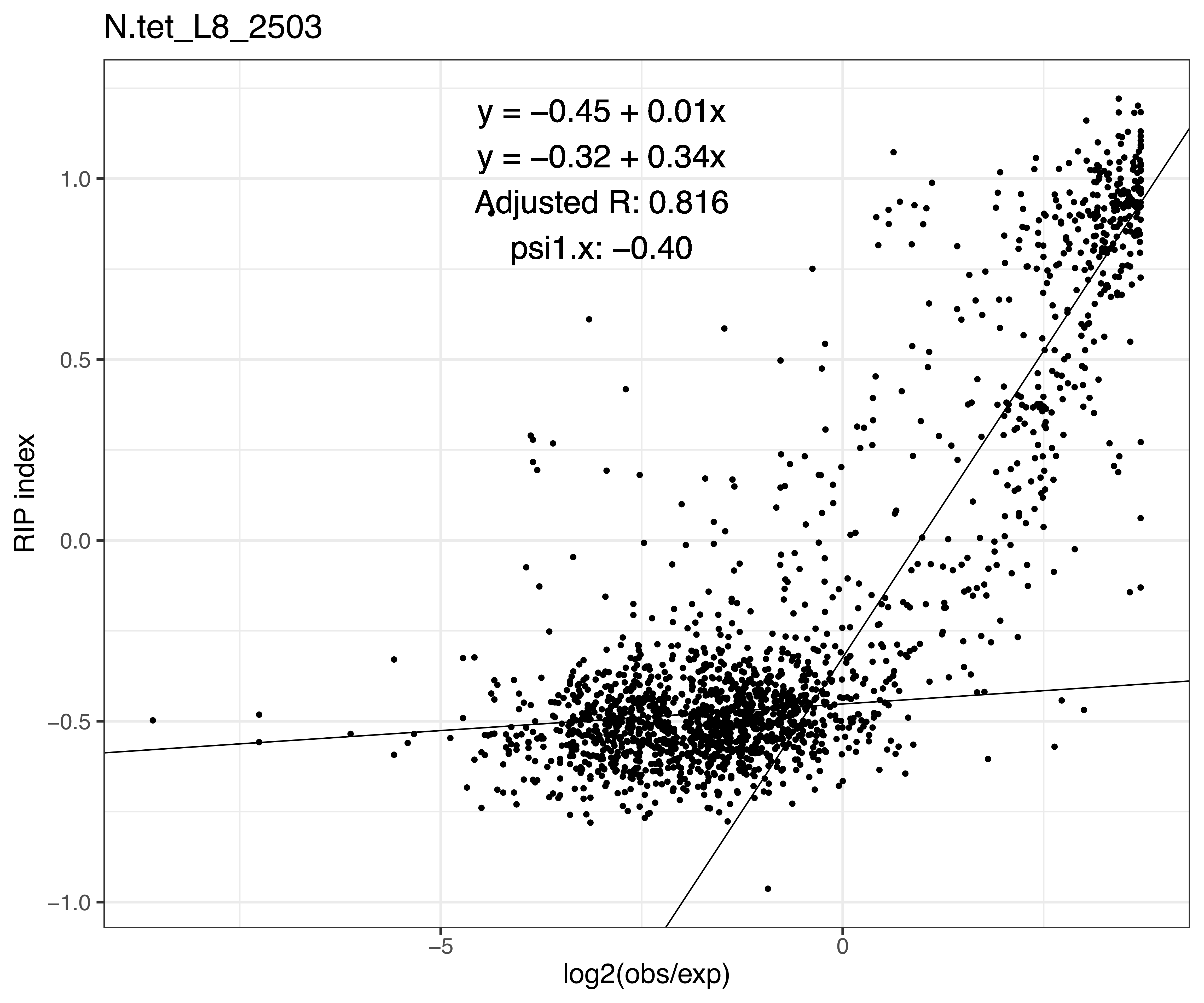

N.tet_L8_2503
y = −0.45 + 0.01x
y = −0.32 + 0.34x
Adjusted R: 0.816
psi1.x: −0.40
1.0
0.5
0.0
−0.5
−1.0
RIP index
−5
0
log2(obs/exp)

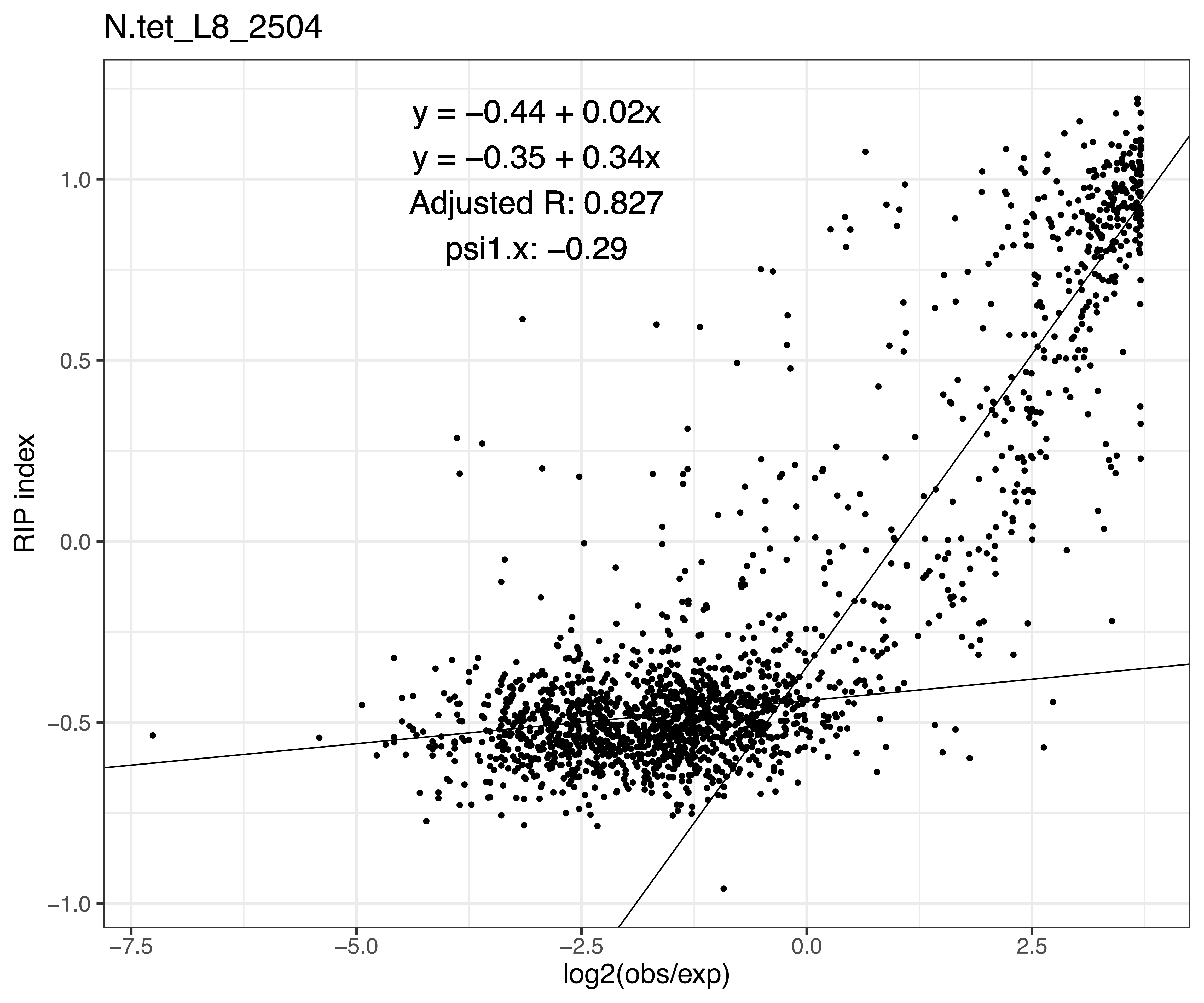
N.tet_L8_2504
y = −0.44 + 0.02x
y = −0.35 + 0.34x
Adjusted R: 0.827
psi1.x: −0.29
RIP index
1.0
0.5
0.0
−0.5
−1.0
−7.5
−5.0
−2.5
0.0
2.5
log2(obs/exp)

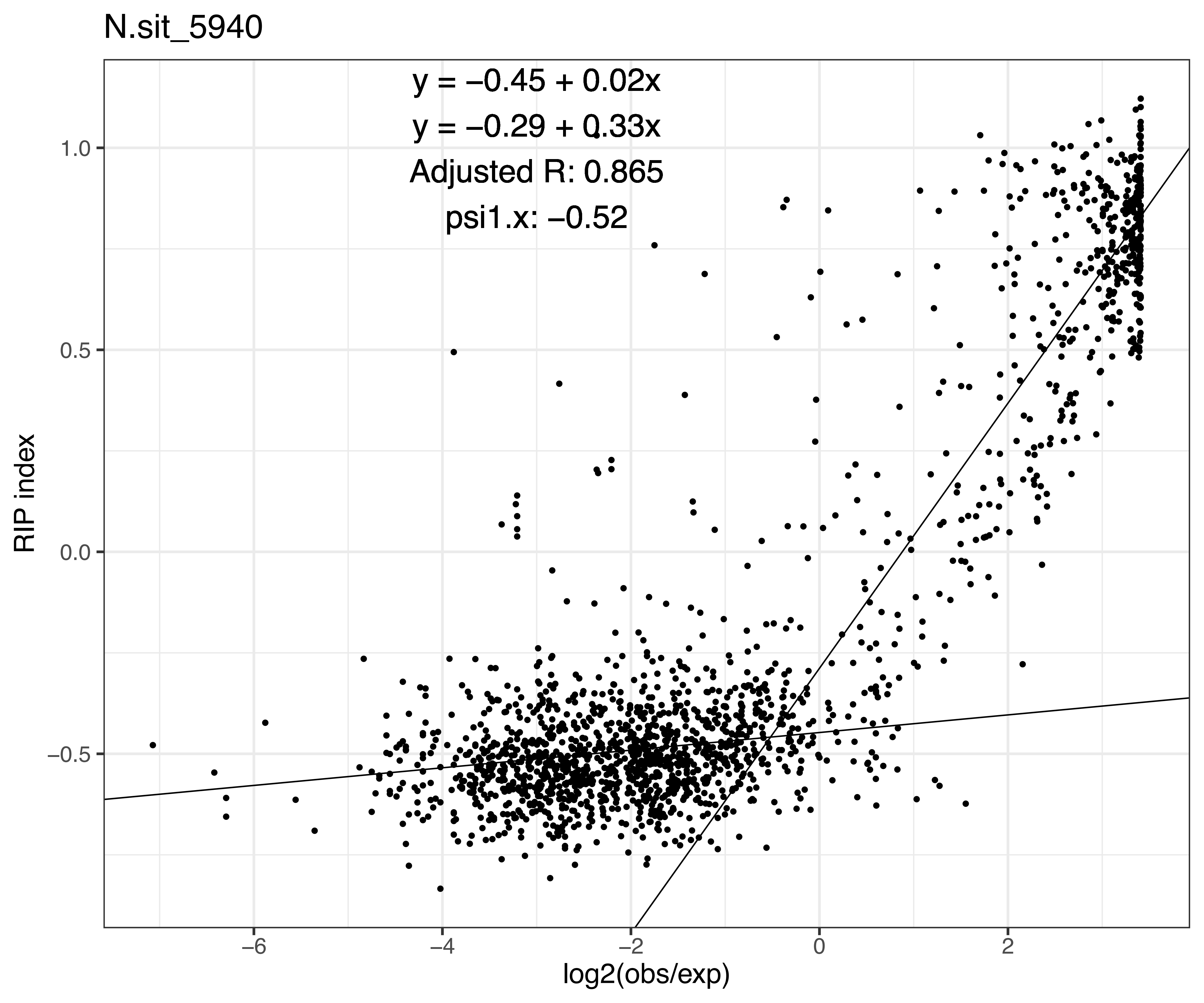
N.sit_5940
y = −0.45 + 0.02x
y = −0.29 + 0.33x
Adjusted R: 0.865
psi1.x: −0.52
RIP index
1.0
0.5
0.0
−0.5
−6
−4
−2
0
2
log2(obs/exp)

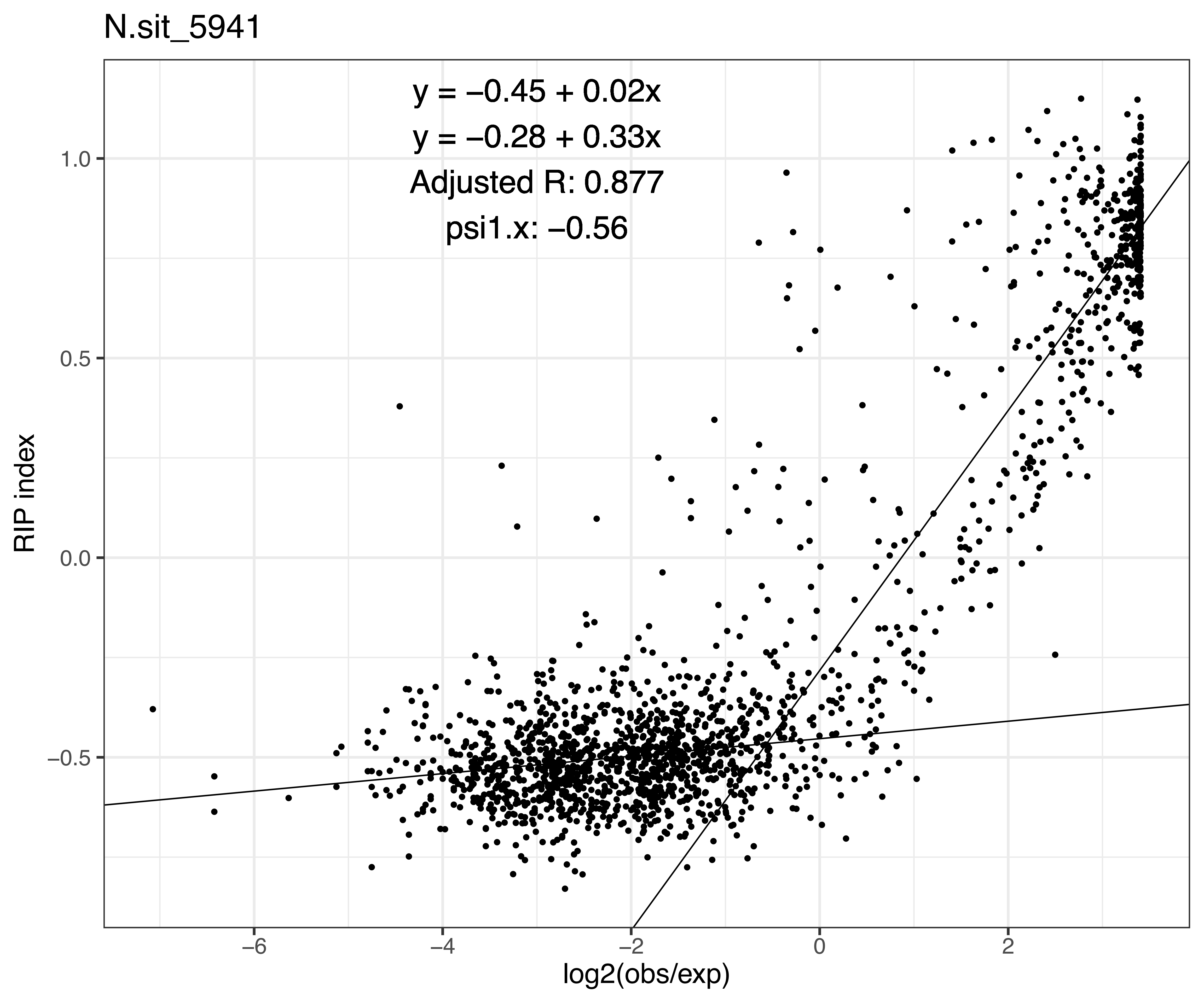
N.sit_5941
y = −0.45 + 0.02x
y = −0.28 + 0.33x
Adjusted R: 0.877
psi1.x: −0.56
RIP index
1.0
0.5
0.0
−0.5
−6
−4
−2
0
2
log2(obs/exp)

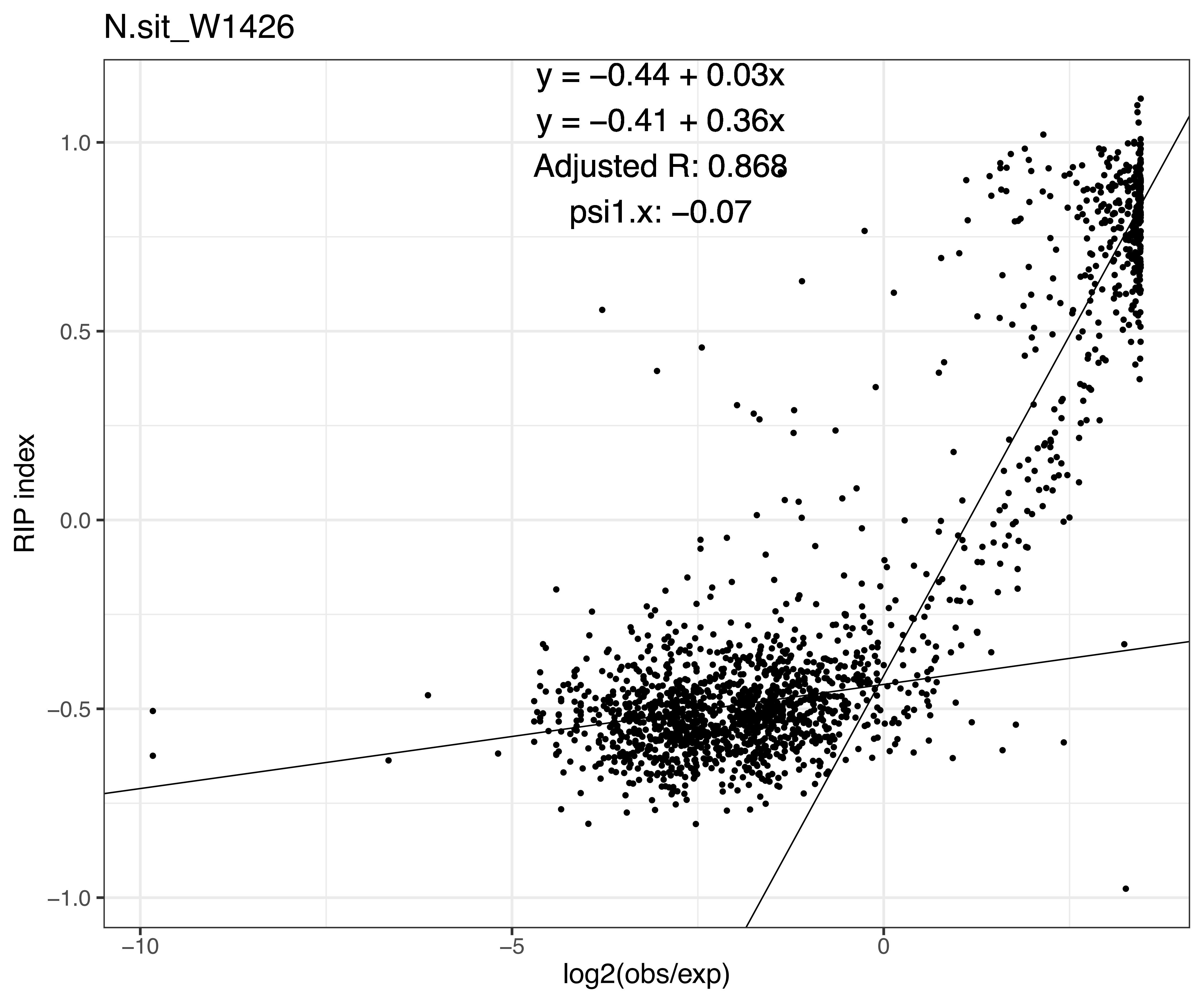
N.sit_W1426
y = −0.44 + 0.03x
y = −0.41 + 0.36x
Adjusted R: 0.868
psi1.x: −0.07
RIP index
1.0
0.5
0.0
−0.5
−1.0
−10
−5
0
log2(obs/exp)

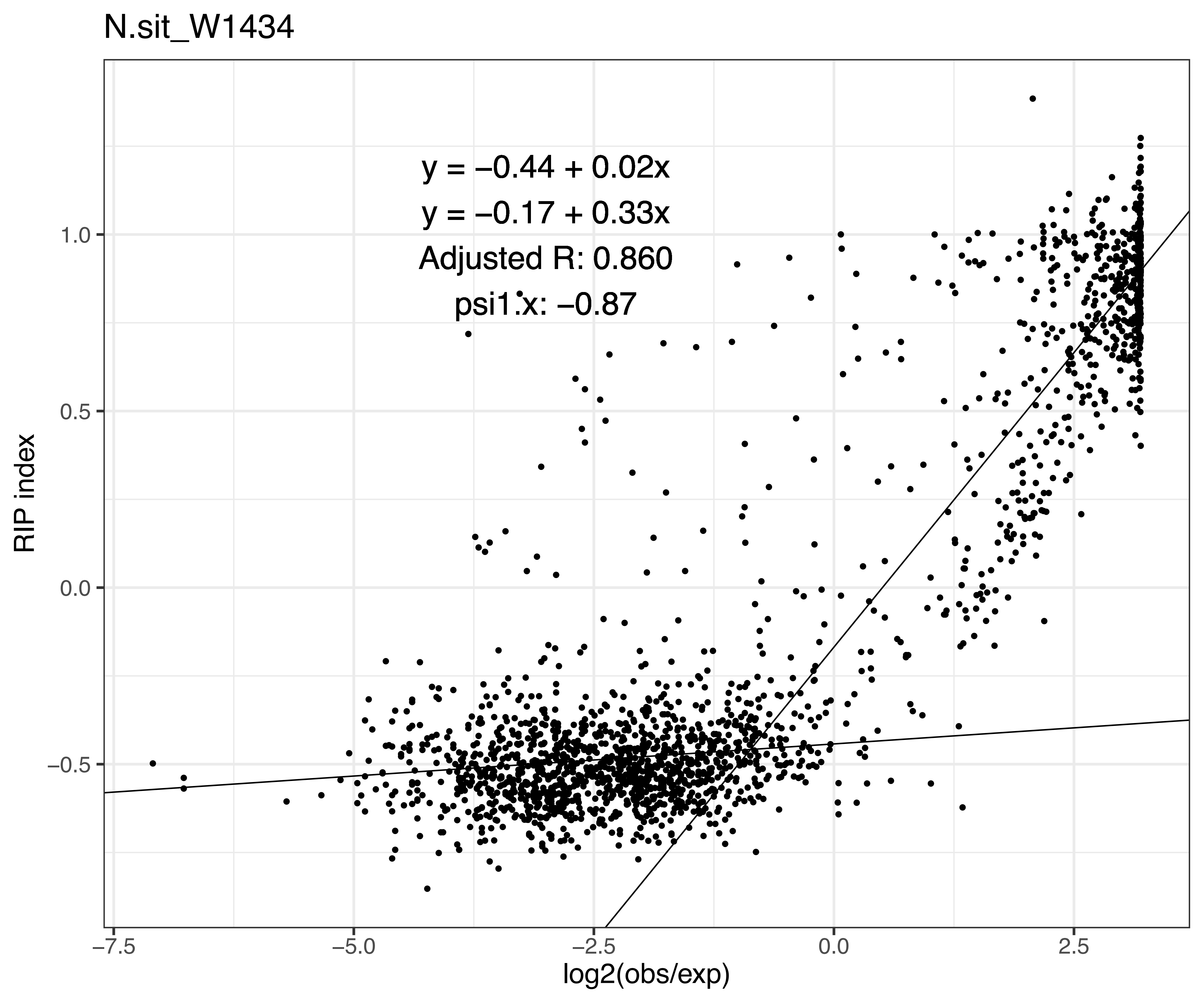
N.sit_W1434
y = −0.44 + 0.02x
y = −0.17 + 0.33x
Adjusted R: 0.860
psi1.x: −0.87
RIP index
log2(obs/exp)
1.0
0.5
0.0
−0.5
−7.5
−5.0
−2.5
0.0
2.5

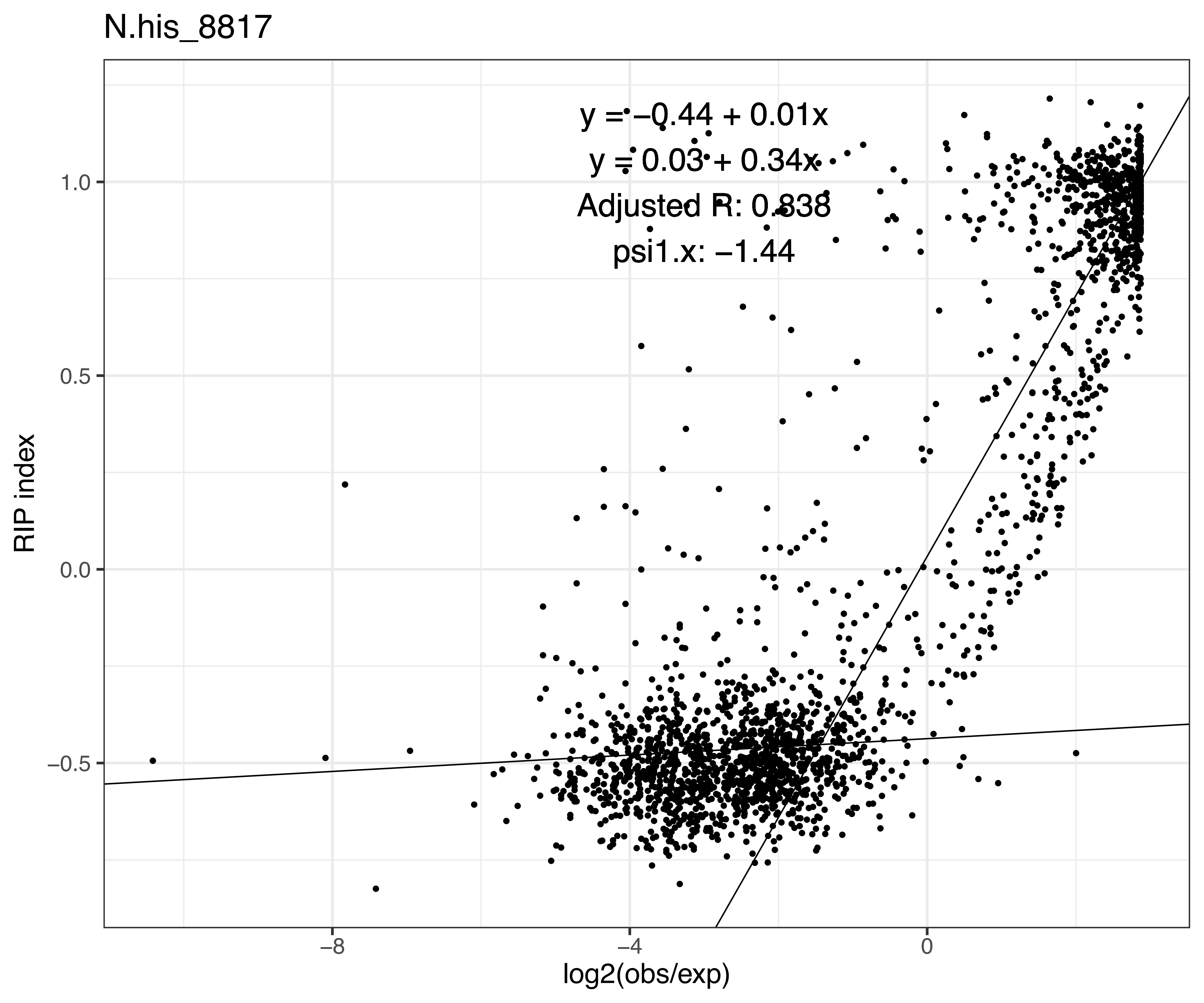

N.his_8817
y = −0.44 + 0.01x
y = 0.03 + 0.34x
Adjusted R: 0.838
psi1.x: −1.44
1.0
0.5
0.0
−0.5
RIP index
−8
−4
0
log2(obs/exp)

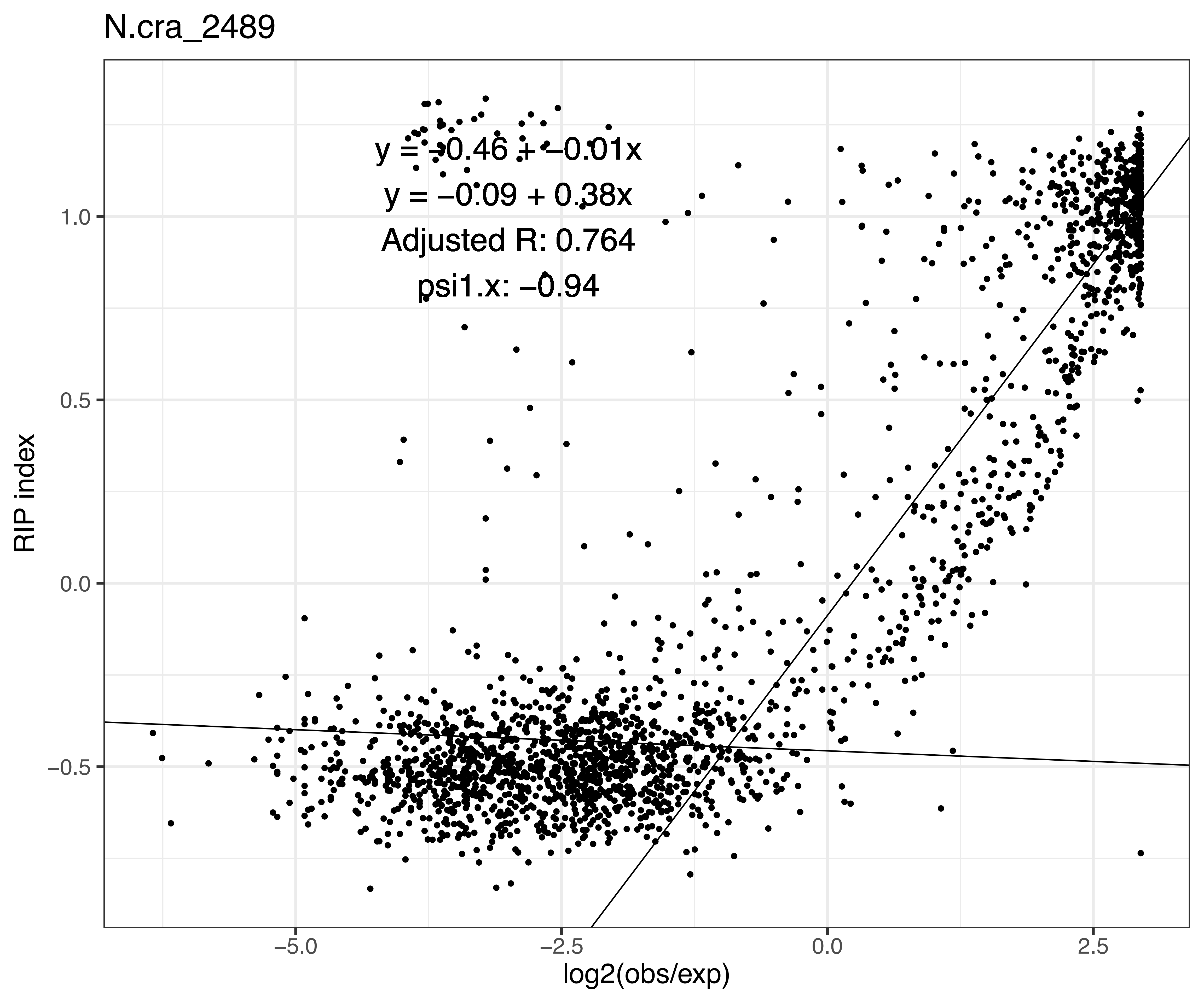
N.cra_2489
y = −0.46 + −0.01x
y = −0.09 + 0.38x
Adjusted R: 0.764
psi1.x: −0.94
RIP index
1.0
0.5
0.0
−0.5
−5.0
−2.5
0.0
2.5
log2(obs/exp)

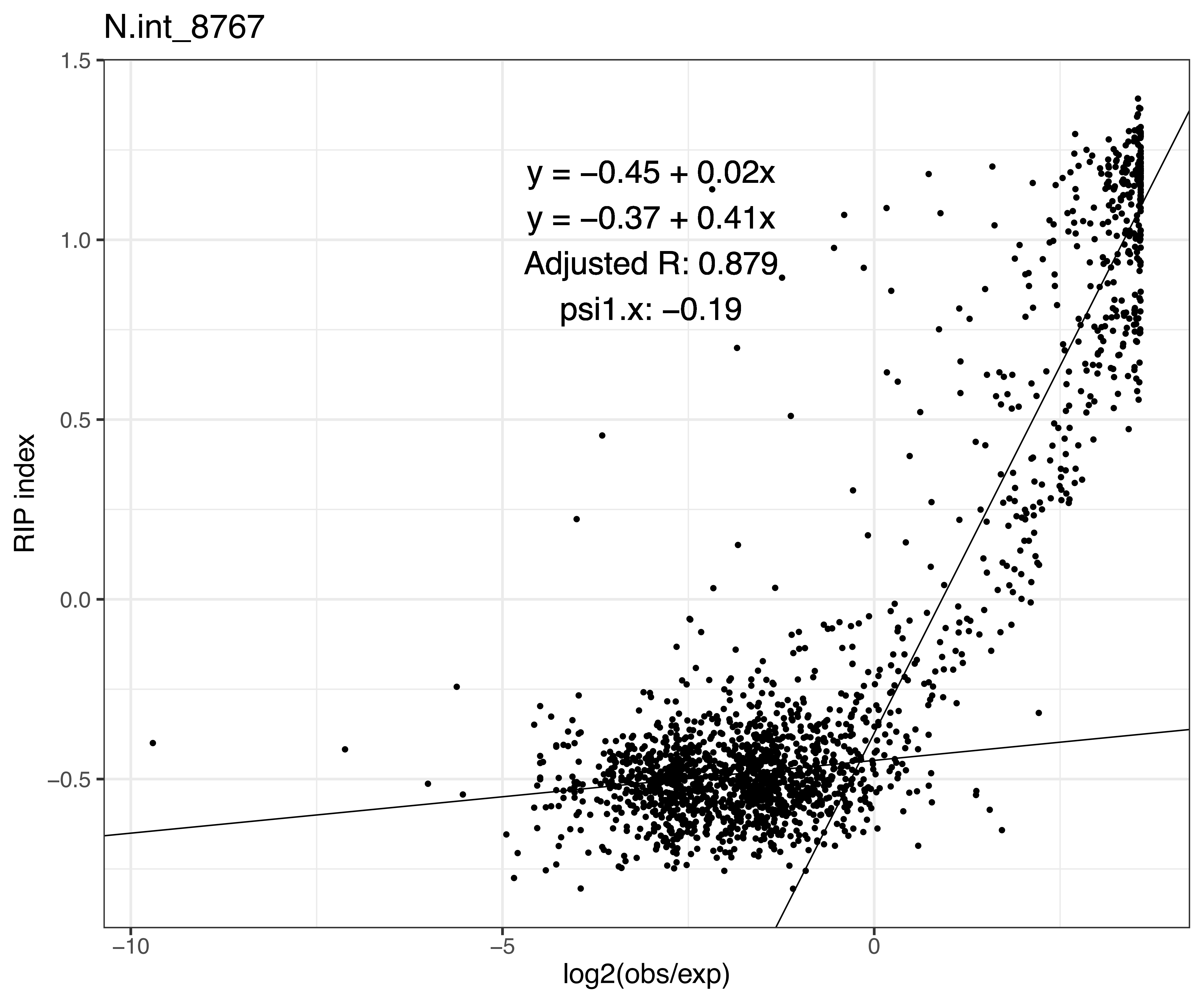
N.int_8767
y = −0.45 + 0.02x
y = −0.37 + 0.41x
Adjusted R: 0.879
psi1.x: −0.19
RIP index
log2(obs/exp)
1.5
1.0
0.5
0.0
−0.5
−10
−5
0

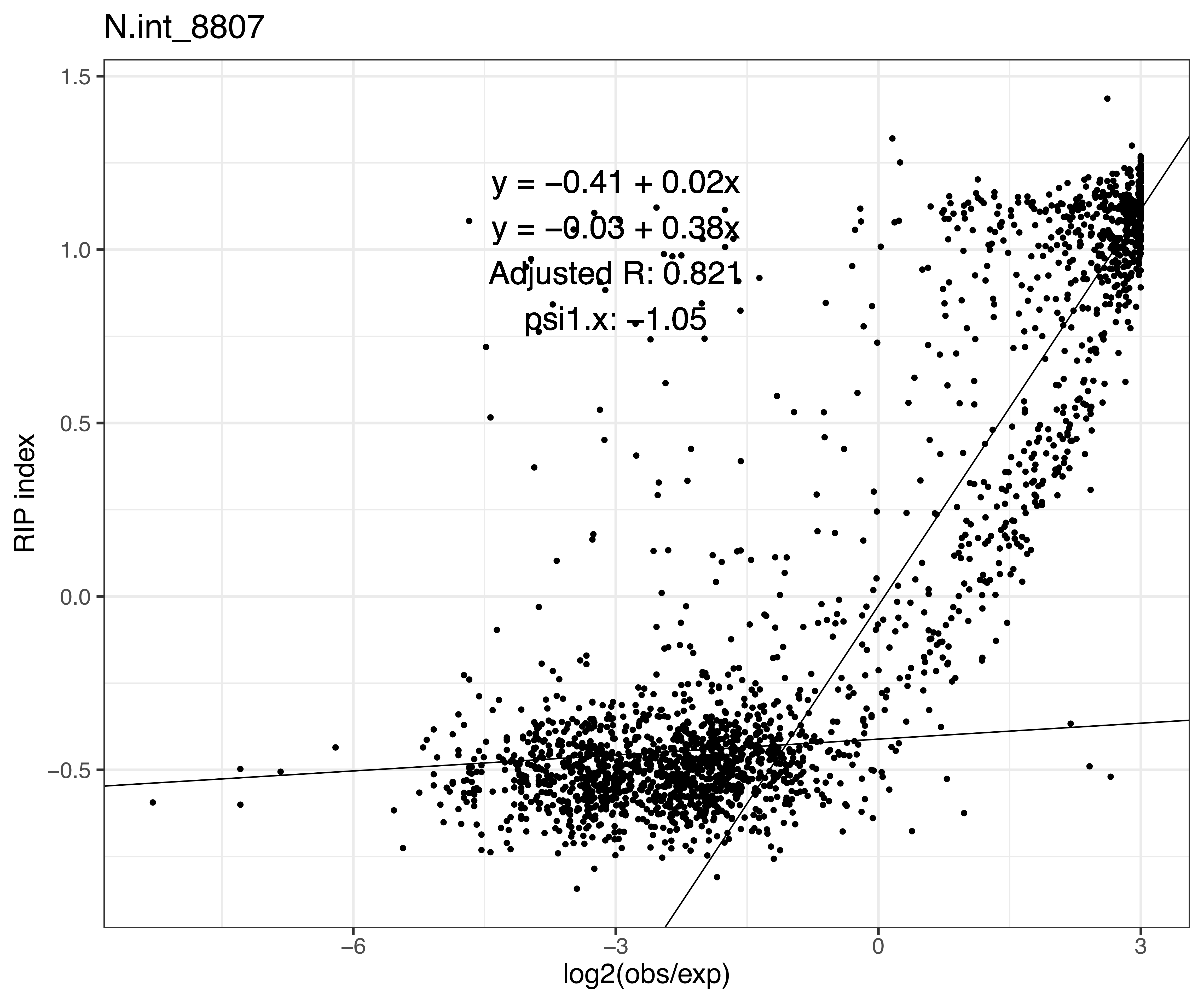
N.int_8807
y = −0.41 + 0.02x
y = −0.03 + 0.38x
Adjusted R: 0.821
psi1.x: −1.05
RIP index
log2(obs/exp)
1.5
1.0
0.5
0.0
−0.5
−6
−3
0
3

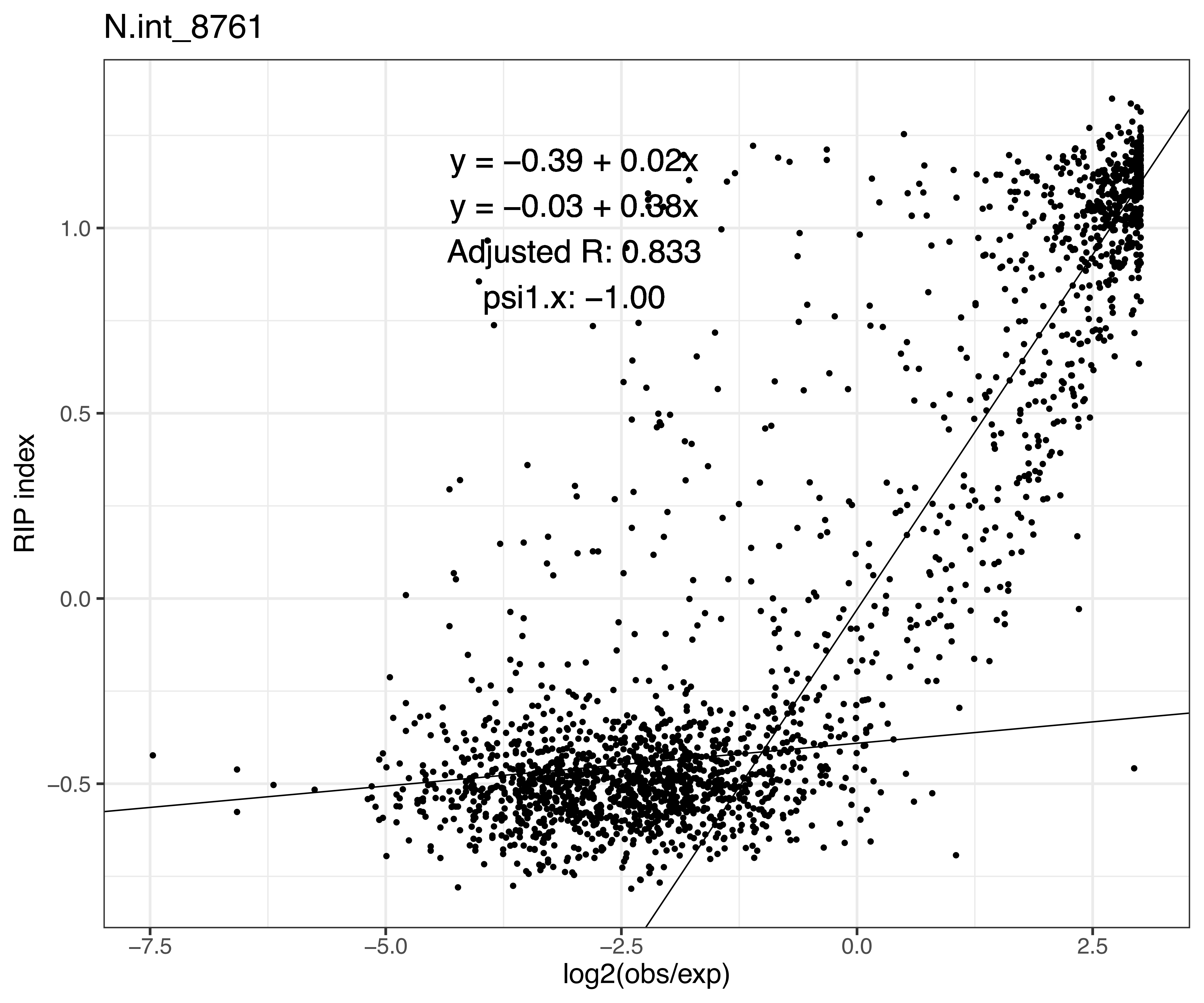
N.int_8761
y = −0.39 + 0.02x
y = −0.03 + 0.38x
Adjusted R: 0.833
psi1.x: −1.00
RIP index
1.0
0.5
0.0
−0.5
−7.5
−5.0
−2.5
0.0
2.5
log2(obs/exp)

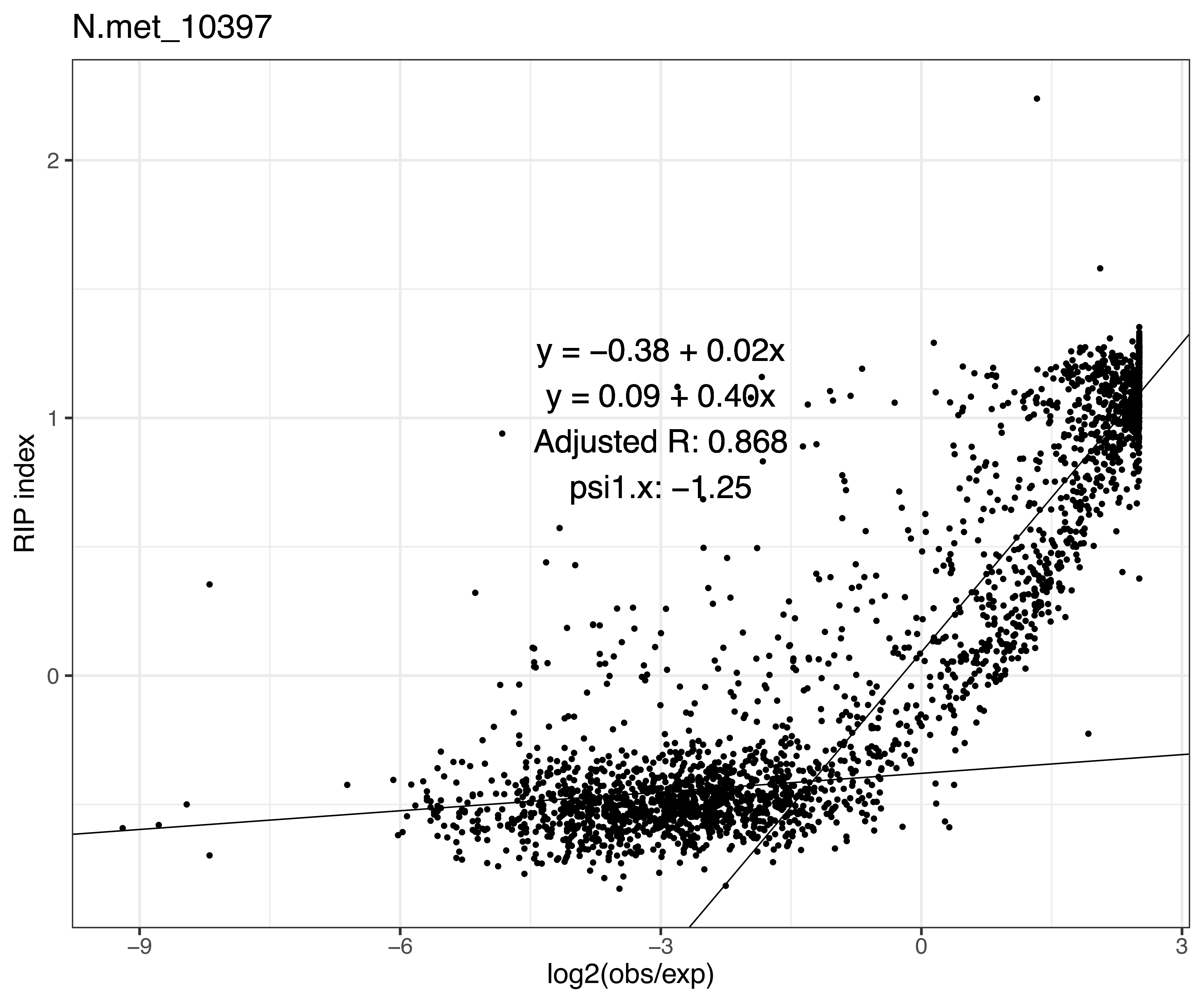
N.met_10397
y = −0.38 + 0.02x
y = 0.09 + 0.40x
Adjusted R: 0.868
psi1.x: −1.25
RIP index
log2(obs/exp)
2
1
0
−9
−6
−3
0
3

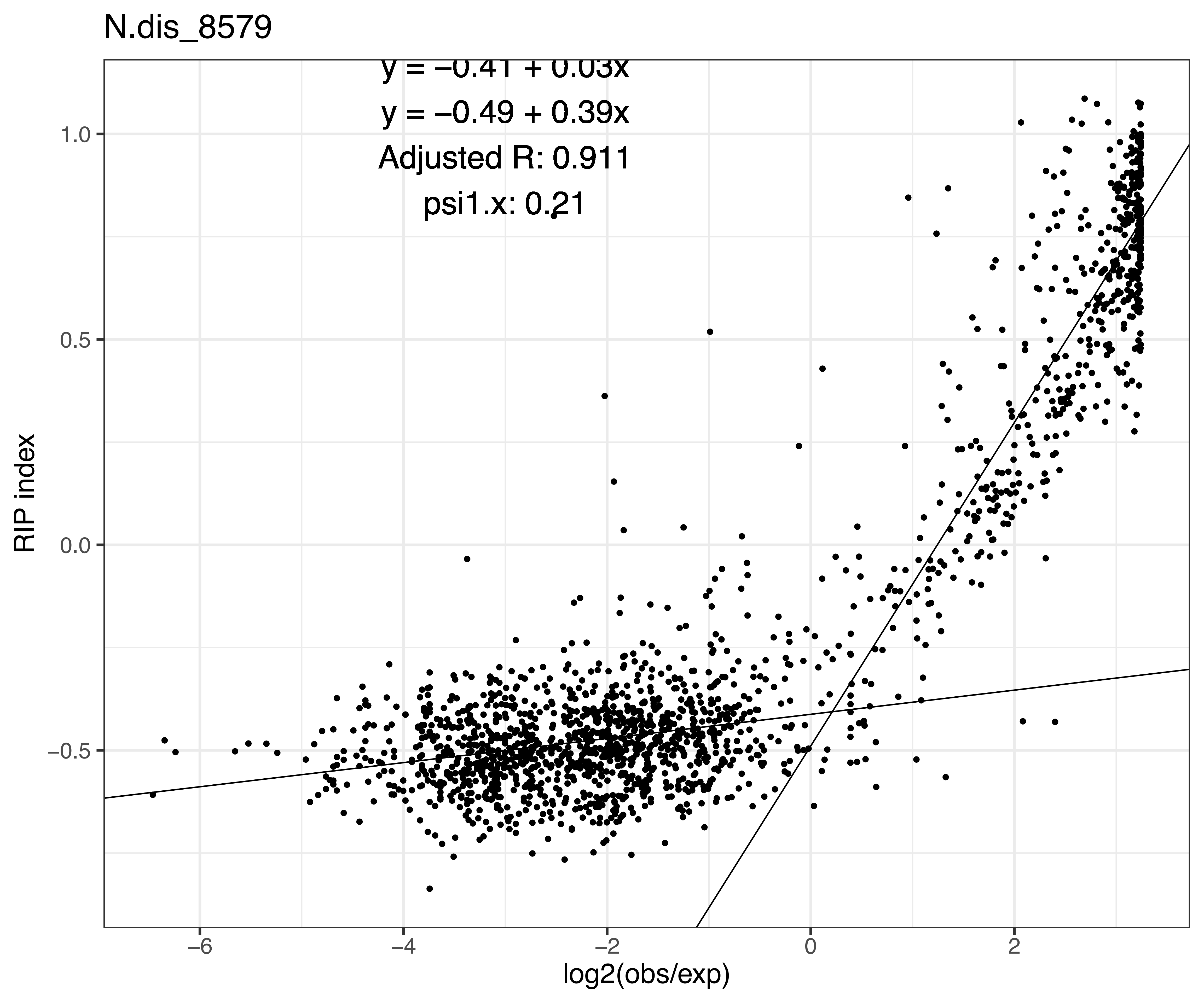
N.dis_8579
y = −0.41 + 0.03x
y = −0.49 + 0.39x
Adjusted R: 0.911
psi1.x: 0.21
RIP index
1.0
0.5
0.0
−0.5
−6
−4
−2
0
2
log2(obs/exp)
