## Supplementary Figure legend for "Transposon- and genome dynamics in the fungal genus *Neurospora*: insights from nearly gapless genome assemblies"

**Supplementary Figure Legends**

**Figure S1.** Distribution of transposable element (TE) sequences in *Neurospora* genomes. a) Composition of TEs in each genome was determined by calculating the number of TE nucleotides relative to the genome length (reflected as percent) for each of the TE families: Class I RNA retroelements which were divided into LTR, SINE, LINE; Class II DNA transposons; and TEs that were unclassified (“Unknown”). b) Total length of TEs in each genome (bp) was plotted for each of the TE families.

**Figure S2.** Correlations between total length of TE families and other repeat sequences, and *Neurospora* genome size. Total lengths of a) DNA, b) LTR, c) LINE, d) SINE, e) Unknown elements, f) low complexity, and g) simple repeat sequences were plotted again the total length of *Neurospora* genomes. Each species was color coded, and for different lineages of *N. tetrasperma*, they were grouped together. Each bubble was associated with a number, relating strain information: **1**, *N. tetrasperma* L6 FGSC 2508; **2**, *N. tetrasperma* L6 FGSC 2509; **3**, *N. tetrasperma* L9 FGSC 10752; **4**, *N. tetrasperma* L7 FGSC 9045; **5**, *N. tetrasperma* L7 FGSC 9046; **6**, *N. tetrasperma* L8 FGSC 2503; **7**, *N. tetrasperma* L8 FGSC 2504; **8**, *N. sitophila* FGSC 5940; **9**, *N. sitophila* FGSC 5941; **10**, *N. sitophila* W1426; **11**, *N. sitophila* W1434; **12**, N*. hispaniola* FGSC 8817; **13**, *N. crassa* FGSC 2489; **14**, *N. intermedia* FGSC 8767; **15**, *N. intermedia* FGSC 8807; **16**, *N. intermedia* FGSC 8761; **17**, *N. metzenbergii* FGSC 10397; **18**, *N. discreta* FGSC 8579.

**Figure S3.** TE landscape distribution in *Neurospora* genomes and genome-wide RIP pattern in *Neurospora* genomes. Top panels. Enrichment of transposable element (TE) sequences (log2(observed repeat (bp)/expected repeats (bp)) were plotted for each *Neurospora* genome in 10 kb windows. Each dot represented a 10-kb window. Values above 2 were herein described as a window enriched in TE sequences. Bottom panels. Genome-wide composite RIP index, irrespective of underlying genomic content, were determined in 10 kb windows, using a custom script. Each dot represented a 10-kb window. Positive values were herein described as a window contained sequences that experienced RIP mutation. For these plots, the alternating colors between blue and green indicate the alternation between chromosomes, following the alignment to *N. crassa*. The lighter shading indicated the presence of multiple contigs for the respective chromosome.

**Figure S4.** Background level of RIP signal was different from RIP signal detected at TE sequences. RIP scores were calculated for a *Neurospora* genome compared to mock sequences, i.e. mock sequences representing a background. For each panel, three distributions are plotted: 1) the observed RIP distribution for a genome calculated on the repeats determined by RepeatMasker (“obs”, purple distribution). Then, RIP scores were recalculated by randomly placing the observed repeat intervals onto mock genomes that were generated in two different ways: 2) a random genome obtained by shuffling the bases (“shuffle”, blue distribution) and 3) generated by sampling nucleotides from the observed nucleotide frequency distribution (“frequency”, yellow distribution). Cases 2 and 3 will disrupt all repeat regions such that no RIP signal should be observed.

**Figure S5.** Linear correlation between enriched TE windows and RIP-positive windows in *Neurospora* genomes. For each of the 18 *Neurospora* genomes, enrichment of transposable element (TE) sequences (log2(observed repeat (bp)/expected repeats (bp)) were plotted for each *Neurospora* genome in 10 kb windows. Positive values indicated that the window was enriched in TEs. Genome-wide composite RIP index, irrespective of underlying genomic content, were determined in 10 kb windows, using a custom script. Positive values were herein described as a window contained sequences that experienced RIP mutation. The regression lines were separately calculated for log2 scores <0 (top regression, left line) and log2 scores >0 (bottom regression, right line), with the adjusted R value calculated for the bottom regression, right line when the log2 scores >0.
